# Complex-trait dissection with Evolve-and-Resequence: Selection for improved flight performance in laboratory fruit fly populations

**DOI:** 10.64898/2026.10.15.759414

**Authors:** Srikant Venkitachalam, Natalie Heath, Adam Albright, Mary York, Mik Kuehnel, Kayla Robinson, Sheila Vazquez, Kevin M. Middleton, Elizabeth King

## Abstract

Evolve-and-Resequence studies combine experimental evolution with molecular sequencing, enabling associations between complex trait phenotypes and evolved genomic changes. For this, a targeted complex trait is subjected to artificial laboratory selection in genetically variable populations. Using populations of fruit flies *Drosophila melanogaster* from the DSPR background, we have conducted a short-term selection study on a highly complex trait – flight performance. We selected for improved flight performance using a custom-built flight tunnel system, with light as a motivator and airflow as a resistive factor. As a result, flight performance significantly improved in selected populations compared to controls. We also observed signatures of evolution in flight related traits, such as activity level and wing shape, and in life-history related traits such as adult dry mass and fecundity. We used whole genome pooled sequencing on the selected and control populations, revealing significantly differentiated genomic sites. Gene ontology and enrichment analyses revealed the evolution of pathways involved in response to stimuli, growth, and development. We identified several genes of interest (e.g., *Sdc*, *dally*). The range of evolved phenotypes and associated genomic changes show that flight performance is an ideal complex trait for future Evolve-and-Resequence studies.

## Introduction

A large number of traits in multicellular plants and animals are known to be complex in nature, with many underlying genes-to-phenotypes associations (Fisher et al. 1932; Laughlin 1933; Lander and Schork 1994; Glazier et al. 2002). These genes-to-phenotypes associations are further complicated by nuanced relationships with the developmental (Bonnett 1964) and ambient environments of organisms (Allard and Bradshaw 1964). Complex traits thus represent the combined actions and interactions of many levels of biological organization (Garland et al. 2016). This complexity makes robust mapping between genotypes and phenotypes extremely difficult. As a result, mechanistic studies of traits often utilize experiments in isogenic backgrounds, through inducing large-effect deleterious mutations (reviewed in Tautz et al. 2026). Such experiments necessarily overlook many facets of complex traits, such as gene-by-environment interactions, epistatic effects, pleiotropy (Tautz et al. 2026). Despite such difficulties, researchers can gain a deeper understanding of biological systems by unravelling the genotype-phenotype associations of such complex traits across multiple levels. Complex trait research has contributed to studies of human traits (Yengo et al. 2018; Conery and Grant 2023), disease biology (Frazer et al. 2009), wildlife conservation (Harrisson et al. 2014), behavior (Plomin et al. 1994; Garland et al. 2016), and evolution (Garland et al. 2016; Sella and Barton 2019).

In particular, laboratory selection experiments on populations of model organisms can be extremely useful (Burke and Rose 2009; Kawecki et al. 2012; Tautz et al. 2026), where large scale association studies are feasible (Mackay and Anholt 2006; Mackay 2009; King et al. 2012b). Laboratory populations of fruit flies of the genus *Drosophila* are a cornerstone for such ‘Evolve- and-Resequence’ studies (Long et al. 2015). The Evolve-and-Resequence (E&R) method combines traditional experimental evolution studies with molecular tools and analyses (Burke and Long 2012; Schlötterer et al. 2014). The key steps in an E&R study are as follows:

i. Researchers impose a specific selection pressure on replicate experimental populations to influence the evolution of a candidate complex trait. Simultaneously, they maintain replicate non-selected control populations.
ii. After multiple generations of selection, they compare experimental populations with non-selected controls to measure the evolution of the complex trait under selection. Additionally, the researchers typically measure a suite of other phenotypes to identify sub-traits associated with the evolution of the candidate complex trait.
iii. Finally, ‘omics’ methods (e.g., whole genome pooled sequencing) enable the analyses of molecular evolution in the selected populations. This identifies genomic variants, typically Single Nucleotide Polymorphisms (SNPs), which may be associated with the evolved complex trait. The differentiated SNPs are further mapped to genes and biological pathways that may be affected.
iv. If ancestral genomic data are available, researchers can map the genomes of selected and control populations to founder haplotypes, permitting a deeper analysis of the evolved genomes.

While the experimental evolution method has existed for at least half a century (Garland and Rose 2009), E&R research has gained popularity owing to the relative affordability of whole-genome sequencing (Kofler and Schlötterer 2014; Long et al. 2015; Vlachos and Kofler 2019; Santos et al. 2024). Examples of candidate complex traits in *Drosophila* for E&R studies include pre-adult development time (Burke et al. 2010), senescence (Graves et al. 2017), opposing life-history patterns (Greenspan et al. 2024), different nutritional environments and age of reproduction (Hoedjes et al. 2019), hot and cold daily temperature fluctuations (Tobler et al. 2014), courtship song (Turner and Miller 2012), learning and memory (Hamlin et al. 2025), temperature adaptation (Santos et al. 2024), resistance to copper (Everman et al. 2023). Researchers have also utilized other model organisms for E&R experiments, for example *Drosophila simulans* (Barghi et al. 2017), *Callosobruchus* beetles (Rêgo et al. 2025), mice (Hillis and Garland 2023), yeast of the species *Saccharomyces cerevisiae* (Burke et al. 2014b; Phillips et al. 2020), *Arabidopsis* plants (Wu et al. 2026), and bacteria (Tenaillon et al. 2016; Burghardt et al. 2018).

Given the past success of selection experiments in evolving selected traits and demonstrating genomic evolution, studying traits at very high levels of biological organization should lead to better holistic understanding of genotype-phenotype associations in complex trait evolution. Flight is one such promising candidate complex trait in *D. melanogaster*.

### Flight performance as a candidate complex trait

“If seeing is believing, then, even by casual observation, flight in insects ranks among the most complex and demanding behaviors exhibited by animals. Flies in particular occupy the pantheon of nature’s aviators.” – (Frye and Dickinson 2004, page 729)

Upon scrutinizing the plethora of traits present in flies, few would match the complexity of flight. Overall, insect flight features the integrated functioning of physiology and anatomy of many organ systems at different levels of organization and is influenced by nuanced neural control and behavioral patterns. Indeed, flight in *D. melanogaster* has been a consistent subject of study for at least the last six decades (Vogel 1967a,b; Frye and Dickinson 2004; Dickson et al. 2008; Hedrick et al. 2015). Fly flight is affected by several aerodynamic factors (reviewed in Taylor 2001; Sane 2003; Sun 2014) such as wing morphology (Vogel 1967a; Krishna et al. 2020; Flaibani et al. 2024), wing beat frequency (Lehmann and Dickinson 1997), body saccades (Cellini and Mongeau 2020), haltere-wing coupling (Deora et al. 2021). Additionally, flight muscles (reviewed in Marden 2000) as well as neuromuscular physiology and coordination (Sadaf et al. 2015; reviewed in Frye and Dickinson 2004) are critical. At a cellular scale, studies have also highlighted the role of indirect flight muscle genetics and flight metabolism (Barnes and Laurie-Ahlberg 1986). Other factors such as endurance (Graves et al. 1988) and flight experience (Hesselberg and Lehmann 2009) among others, also play important roles. Furthermore, mating songs in males are also coupled with aspects of flight (Ewing and Bennet-Clark 1968). Flight is also regulated by motivational states for flight initiation, maintenance and termination (Brembs et al. 2007).

Thus, direct selection on any aspect of flight performance could affect many traits involved in flight. At least two different reliable methods have been developed for large scale measurement of flight performance in *D. melanogaster*. One method tested the flight recovery ability of flies after a vertical free fall into a tube (Babcock and Ganetzky 2014). This method has featured extensively in molecular association studies using panels of *D. melanogaster* lines (Spierer et al. 2021; Darwin et al. 2025). The other method, first published by Weber (1988), measured ‘upwind flight ability’ in a wind tunnel. The author who developed this method also carried out a long-term experimental evolution study on improved flight performance (Weber 1996; Marden et al. 1997). The wind tunnel used to select for flight performance was a closed dark chamber with light on one end, which served to attract flies. Early studies established that *D. melanogaster* could be highly phototactic (Hirsch and Boudreau 1958; Hadler 1964), and visual cues may also be crucial to feedback in flight control in *D. melanogaster* (Tammero and Dickinson 2002). In the flight tunnel, as flies move towards the light source at the far end, a fan exerted wind resistance, impeding their flight to the light source. The authors tested several thousand flies through this tunnel at a given time, fractionating flies based on their upwind traversal in this wind tunnel. The selection process consisted of breeding the furthest traversing flies to form the next generation. As suggested by Weber (1996), the flight performance trait measured in this setup was “*actually a composite of phototaxis, activity level, flying speed, and aerial maneuvering skill.*”

### The Current Study: Evolve-and-Resequence for improved flight performance

Given the past evidence cited above, we believe that flight in *D. melanogaster* is an ideal complex trait for an E&R study. In the current study, we describe results from a laboratory selection experiment for improved flight performance on populations of *D. melanogaster* from the Drosophila Synthetic Population Resource (DSPR; King et al. 2012a,b).

We chose to select on upwind-flight ability (Weber 1988, 1996), because it showed promising selection results for the evolution of improved flight performance, but here we expanded the study to include the following advances:

1. At the time of the previous study, whole-genome sequencing tools were not available. This was our primary motivator for conducting a follow-up study.
2. While the original study conducted each flight performance run with a very large number of flies (approx. 10,000), the setup used only two sets of replicate populations. Low population replicate numbers could lead to unreliable results in an E&R study (Kofler and Schlötterer 2014). We used 4 replicate populations for selection and control.
3. The populations in the study were maintained in bottles, possibly in overlapping generations. Here, we use a design with non-overlapping generations.
4. The evolved phenotypes measured were largely restricted to measures of flight performance. One test on correlated evolution of fitness traits was carried out via a method which conflated fecundity, larval competitive ability and survivorship, making it difficult to disentangle these traits. Here, we expand the suite of traits assayed to identify the multifaceted nature of this complex trait.

Inspired by the designs of Weber, we used a custom-built flight tunnel (see Methods). In this setup, phototaxis in a dark tunnel was the motivator for fly movement, while airflow served as a deterrent, inhibiting forward flight toward the light. Only the flies traversing a certain distance within the tunnel were selected each generation. After 10 generations of selection, we compared the experimental populations to non-selected control populations along several aspects. Our central questions in this study were the following:

1. Can flight performance be improved through evolution in the timespan of a few generations of selection?
2. What contributing, lower-level phenotypes evolve alongside improved flight performance?
3. How does the genome change with selection for improved flight performance?

## Methods

### Populations used, maintenance and selection procedure

The outbred populations utilized in the selection experiment originated from 93 Recombinant Inbred Lines (RILs) from the population ‘A’ DSPR background (King et al. 2012a,b; Long et al. 2014). We have provided the exact IDs of the RILs used in supp. table S1. To create the populations used in this study, we first added 10 mated females from each of the 93 RILs (total 930 females) into two custom-built acrylic cages (20.3cm × 20.3cm × 20.3cm). Seven days later, we provided an egg collection plate to each cage and collected the eggs into polystyrene vials (25 mm outer diameter, 95 mm height, Genesee Scientific) at a density of 60-80 eggs per vial in approx. Approximately 9-10 mL of cornmeal food was added using DrosoFiller (Genesee Scientific). We split the vials collected from each cage into 2 further groupings, such that a total of 4 populations were formed from 20-25 vials each. On day 13 from egg collection, once most eggs had developed into adult flies, we transferred the adults into 4 respective acrylic cages. From these, we collected a fresh batch of eggs to form the next generation on day 21 from egg collection, after providing an egg collection plate on day 20. The subsequent schedule was the same as in the previous generation, with the addition of a thick paste made of brewer’s yeast with deionized water provided from day 18-20 from egg collection. Previous research has shown that live yeast can boost egg output in *Drosophila* (Robertson and Sang 1944). At generation 3, we further split the 4 populations into 8 populations. These became 4 replicate control populations (CON_1-4_) and 4 replicate populations selected for improved flight performance (FP_1-4_). For 10 subsequent generations, we applied the selection process to the FP populations on day 18 from egg collection. In all other respects, we maintained the CON and FP populations in the exact same way. The selection process involved subjecting adult flies from FP_1-4_ populations through a ‘flight performance’ apparatus (figure 1A, supp. fig. S2A). To form each generation, we selected only those flies which could traverse a certain distance in this tunnel. To account for the large number of flies being rejected from the selection process, we collected the eggs over a period of two days each generation (figure 1A). On each day, we collected eggs in around 30 vials. For the selection process on day 18 of a given population, we used two runs of the flight performance tunnel, sourced from around 30 vials each. We have described the maintenance and selection process in a schematic in figure 1. A recent study also contains additional information on the formation of the populations from the 93 RILs, as well as details of maintenance of CON populations (Hamlin et al. 2025).

**Figure 1.**
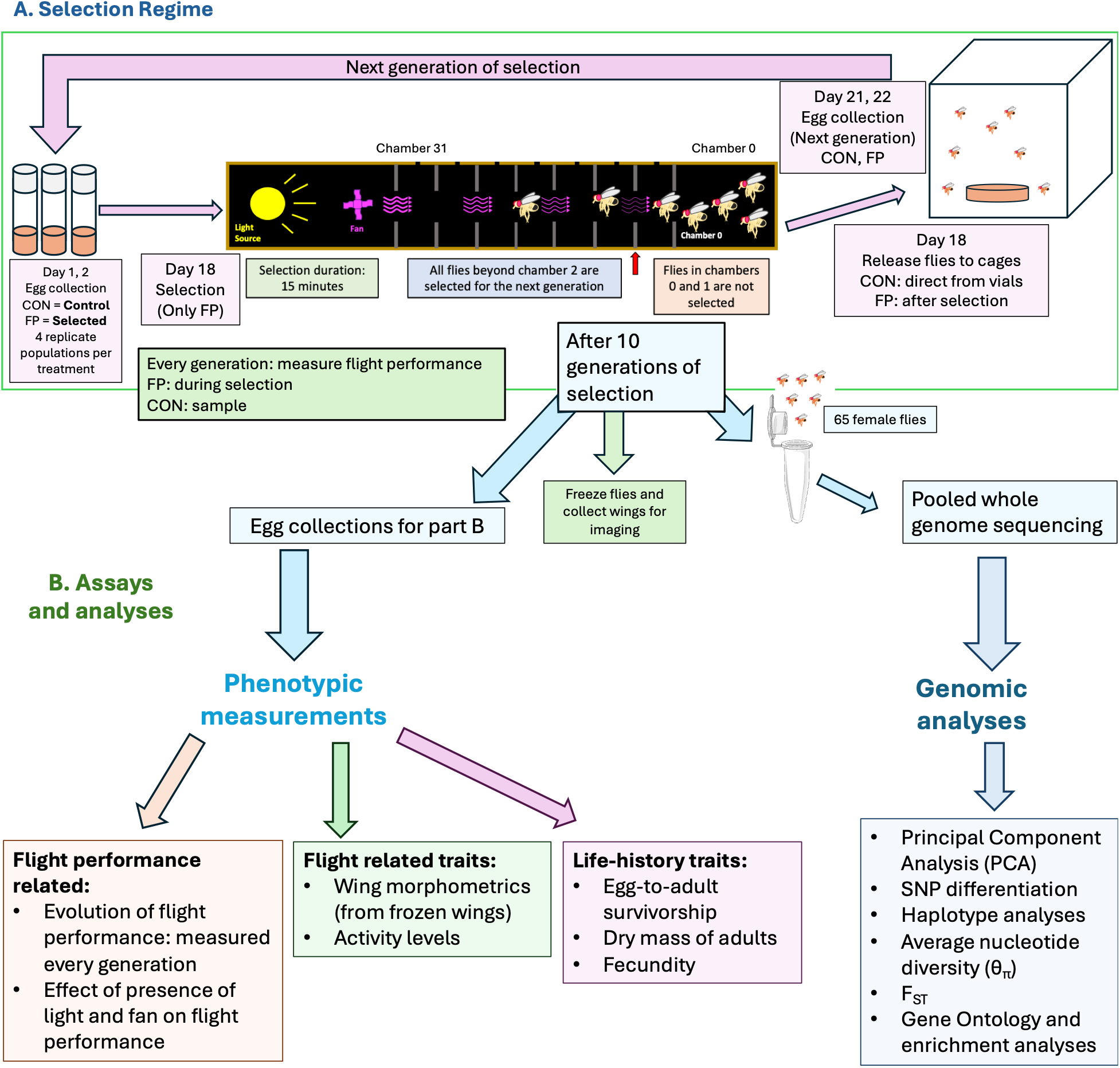
Schematic of the current study. A. The entire selection procedure in a flow chart. Both maintenance of the CON (control) populations and selection of the FP (selected) populations takes place over a 21-day generation cycle. The selection criterion was chamber 1 (C1) for the first 6 generations of FP and then increased to C2 as shown in this figure. See methods for further details. B. All the phenotypic measurement assays as well as genomic analyses carried out after the end of 10 generation of selection. We measured the evolution of flight performance each generation. We also obtained images for wing morphometrics directly from the populations at generation 10.

#### Design of the flight performance apparatus

We based the design and construction of the flight tunnel on the descriptions of the apparatus by Weber (1988), and its usage in subsequent studies (Weber 1996; Marden et al. 1997). The apparatus was a fully enclosed, dark tunnel with 32 chambers (henceforth ‘C0-C31’) and a light source and fan at the C31 end (supp. fig. S2C). To start the flight performance test, we added flies to the tunnel at C0. The light source, two fiber optic lamps, served to attract the flies through the tunnel (supp. fig. S2B). The fan, located adjacent to the light tubes, provided wind resistance to deter flight. We placed mesh-covered air outlets at the top of every chamber to let some of the air escape from each chamber, providing a wind gradient down the length of the tunnel. This meant that the flies faced wind resistance of increasing intensity at distances approaching the light source (wind resistance C31 >> wind resistance C0; supp. fig. S3). Each chamber was separated from the adjacent chambers by an ABS plastic divider with a ∼50 mm diameter hole in the center. The sides of all the dividers were coated with polytetrafluoroethylene (byFormica Ant Products, LLC) preventing flies from crawling through the openings. This setup ensured that the only way flies could travel from one chamber to the next would be to fly through the holes between chambers.

At the beginning of every flight performance run, we collected flies from around 30 vials from a population into a plastic bottle. As soon as the flies were collected, we added them to chamber C0 of the flight tunnel, immediately switching on the light source and fan. After 15 minutes, we switched off the lights and fan and immediately dosed the entire tunnel with CO_2_ for around 15 seconds. This rendered all flies unconscious close to their position at the end of the trial. We subsequently opened the cover of the tunnel, counting and collecting the flies from the selected chamber. If required, we dosed the non-selected flies with more CO_2_, to collect, freeze and count them later.

#### Selection criterion

We started with a lenient threshold of selection for the selected (FP) populations. For the first 6 generations, we selected any fly which could clear the entry chamber (C0; figure 1A). Once flight performance of FP flies visibly improved, we increased the strength of selection. Generations 7 through 10, we selected flies traversing chambers further than C1 (C2 onwards; figure 1A). At the end of the trial, we released selected flies into their respective cages.

### Measurement of phenotypes

After 10 generations of selection, we carried out several assays to study the effects of evolution for improved flight performance. Broadly, we measured direct flight performance, flight related traits and life-history related traits. All assays are listed in fig. 1B.

#### Direct flight performance measurements

##### Evolution of flight performance

Every generation, we measured the flight performance of the FP populations during selection. From generation 3 of selection, we also measured the flight performance of flies sampled from the CON populations.

To analyze the evolution of flight performance, we first fit the following model:

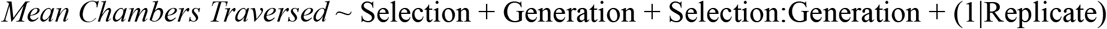

where random factor Replicate was the population replicate (1, 2, 3, or 4) nested within Selection (CON or FP). Here, *Mean Chambers Traversed* was the measure of how far flies travelled on average in the tunnel. We calculated it as the mean chamber number weighted by the proportion of flies found in each chamber. The proportion of flies was the average across replicate runs, when more than one run was used, to address the disparity in the number of runs between CON and FP (see below).

We also fit the above model for both CON and FP separately, because the number of generations differed for both (7 generations for CON, 10 for FP). Given that we measured flight performance of the FP populations as a requirement for selection, they had much more complete data compared to the CON populations. On most occasions, there were 2 runs for each measurement of an FP population, while CON populations had single run data due to logistical limitations. In the case of separate analyses of CON and FP, the model only evaluated the main effect of Generation:

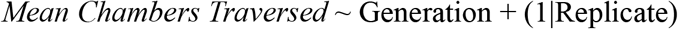

##### Factorial flight performance assay: testing the effects of light and wind stimuli

In addition to the flight performance measurement each generation, we also carried out a ‘factorial flight performance’ assay after 10 generations of selection, to test the effects of the two factors present in the flight tunnel as stimuli – light and wind.

To prepare for this experiment, we collected vials of around 60-80 eggs from both FP and CON populations, one week apart. We collected the vials over two subsequent days – 21 or 22 days from the previous generation’s egg collection. Thereafter, we sampled flies from both days of collection for wind tunnel experiments. During sampling, we noted that some vials had microbial growth in them. We sampled only those vials which showed no/minimal growth. The sampled flies were 17 or 18 days old at the time of the factorial assay.

For the factorial flight performance assay, we tested the effect of light, fan and selection in a fully factorial design. The light was either ‘ON’ or ‘OFF’. The fan was either ‘ON’ or ‘OFF’. The selection factor was either ‘CON’ for control or ‘FP’ for selected.

From 4 randomly chosen vials belonging to either CON or FP, we added flies to chamber C0 of the flight tunnel (fig. 1A, supp. fig. 1). The setup for each run was in 1 of 4 configurations, based on the cross of light status (ON/OFF) and fan status (ON/OFF). For each factorial combination, we used 2 replicate runs of 4 vials each. Some exceptions used 3 vials, and we noted these in the data file. The flies traversed the flight tunnel for 15 minutes as in the selection process of FP populations. Thereafter, we dosed the tunnel with CO2 (after turning off lights and fan, if on). Finally, we counted the number of flies in each chamber.

For statistical analysis of the mean distance traversed under each factorial combination, we carried out model comparison using the Akaike Information Criterion (AICc), via the AICcmodavg package in R (ver. 2.2.3; Mazerolle 2023). The following models were compared:

1. Only main effect of Selection

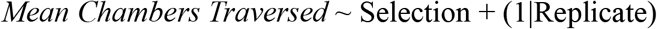
2. Only main effect of Light_Status

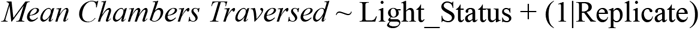
3. Only main effect of Fan_Status

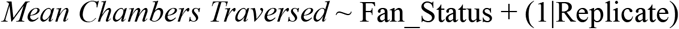
4. Additive model with all three factors

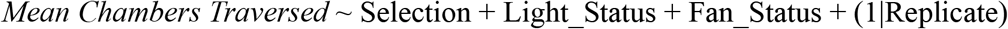
5. Interactions of Selection crossed with Light_Status and Fan_Status

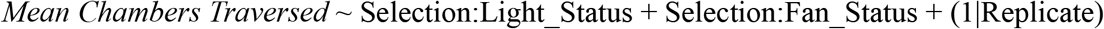
6. Fully factorial model, with all terms included

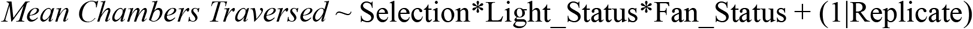

In all models being compared, random factor Replicate was the population replicate (1, 2, 3, or 4) nested within Selection (CON or FP). Light_Status and Fan_Status had 2 levels each (ON/OFF). *Mean Chambers Traversed* was the measure of how far flies travelled on average in the tunnel. We calculated it as the mean chamber number weighted by the proportion of flies found in each chamber, per run.

#### Flight-related traits

##### Activity levels

We measured the overall activity levels of CON and FP flies through their overall movement in a set duration in a custom-built setup. We assayed both individual male and female activity in individual tubes for 5 minutes each, with temperature set at 24°C. The deep learning software, DeepLabCut (Mathis et al. 2018) tracked the position of each fly during the assay and we then used a custom R script to determine the total distance traveled by each fly over the course of the assay. A previous study on thermal tolerance has described the setup and development of the deep learning model to estimate fly activity from videos of flies in individual tubes in detail (Williams-Simon et al. 2024).

To prepare for this experiment, we had two sets of fly collections, each with a different sampling schedule. For each set, we collected vials of around 60-80 eggs from both FP and CON populations, one week apart from each other.

Set 1: these were the same set of vials as the factorial flight performance assay. We collected the vials over two subsequent days – 21 or 22 days from the previous generation’s egg collection. For the assay, we mixed up and sampled the vials from both days of egg collection, discarding any vials that showed major microbial growth. The flies in Set 1 were 17 or 18 days old at the time of the activity assay.

Set 2: we collected these vials over two days, from the subsequent generation of egg collection from both CON and FP populations (no selection imposed). For the assay, we mixed up and sampled the vials from both days of egg collection, discarding any vials that showed major microbial growth. The flies in Set 2 were 17 or 18 days old at the time of the activity assay.

For each population and each assay set, we conducted 3 runs of the activity assay. Each run consisted of a total of 30 flies, 15 males and 15 females in alternating tube arrangements. The outcome variable we considered for the activity assay was the total distance traversed by a fly over the assay period of 5 minutes. During processing of the fly activity data, the deep learning model assigned each tube a likelihood score. We dropped all frames with likelihood < 95% from downstream analyses. Furthermore, if a fly had >= 5% of its total frame numbers dropped, we removed it from consideration.

For statistical analysis of the total distance traversed in the activity assay, we carried out model comparison using the Akaike Information Criterion (AICc), via the AICcmodavg package in R (ver. 2.2.3; Mazerolle 2023). The following models were compared:

1. Single main effect of Selection:

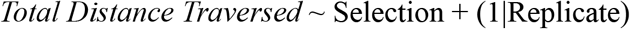
2. Single main effect of Sex:

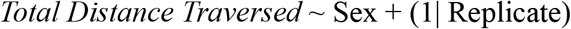
3. Additive model of Selection and Sex:

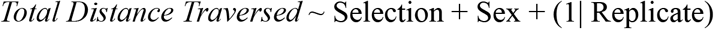
4. Factorial model of Selection and Sex:

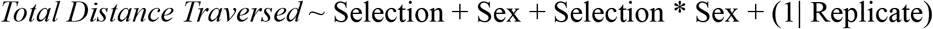
5. Additive model with Selection and Sex, with Assay_set as a fixed factor crossed with Selection:

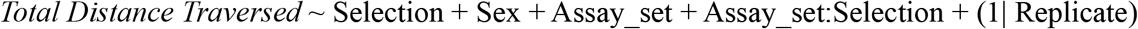
6. Factorial model of Selection and Sex, with Assay_set as a fixed factor crossed with Selection:

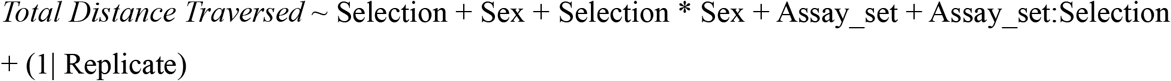
7. Full factorial model with all terms crossed:

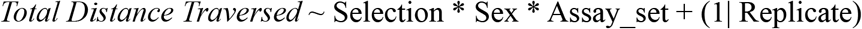
8. We additionally also fit a model where we tested the effect of Assay_set as a random factor. Here, the random terms (1|Assay_set), (1|Sex:Assay_set) and (1|Sex:Selection:Assay_set) had 0 variance, and were omitted from the fit model.

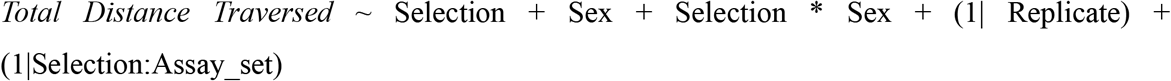

In all models being compared, random factor Replicate was the population replicate (1, 2, 3, or 4) nested within Selection (CON or FP). Sex was a fixed factor (Female or Male). Assay_set (Set1 or Set2) represented the two different sampling events we carried out, on different days of the experiment (see above). The factor Assay_set was a fixed factor for models 1-7 and was tested as a random factor in model 8. A recent study presented arguments for and against using a random factor with <5 levels (Gomes 2022).

### Wing morphometrics

#### Specimen preparation and imaging

Flies were stored for dissection by freezing. Left and right wings were dissected free of the body under a stereomicroscope. Wings were mounted on a glass slide with Permount Mounting Medium (Fisher Scientific) thinned by 50% with mixed xylenes (Fisher Scientific) and allowed to cure prior to imaging. Light microscopy images of wings (n = 598) were acquired at the University of Missouri Advanced Light Microscopy Core Facility using a Nikon Ti2 inverted microscope equipped with a PCO Edge 4.2 sCMOS camera. Image resolution was 1 pixel = ∼1.09 µm.

#### Landmarking

We used twelve landmarks for geometric morphometric analysis (supp. fig. S5), following those used by Houle et al. (2003), which encompass longitudinal and cross veins while omitting some of the proximal wing landmarks used by Gidaszewski et al. (2009). The proximal wing landmarks were in an area of the wing often damaged during dissection. This landmarking scheme was designed to detect large-scale differences in wing size and shape across sex and treatment groups, which were less likely to be obvious in the proximal wing.

To landmark the images, we developed a deep learning model using DeepLabCut (v. 3.0.1; https://deeplabcut.github.io/DeepLabCut/; Mathis et al. 2018, 2019; Nath et al. 2019). Randomly chosen images (n = 40) were manually digitized to provide the training data for transfer learning using an HRNet-32 backbone (Sun et al. 2019; Wang et al. 2020; Wightman et al. 2023). The model was then trained for 200 epochs using PyTorch as the computational backend (Paszke et al. 2019) with the AdamW optimizer (Loshchilov and Hutter 2019). Learning rate varied between 0.0001 (iterations 1 to 160), 0.00001 (iterations 161 to 190), and 0.000001 (iteration 191 to 200). After an initial training/prediction run, we manually refined landmarks in 147 additional wings. These images went back into the training pool for 187 total wings in the training set. The model was then retrained using the parameters above. The final model had a median training error of 1.86 pixels (2.03 µm) and testing error of 6.81 pixels (7.43 µm). After training, this model was used to landmark all 598 images in one batch.

#### Geometric morphometric analyses

For geometric morphometric analysis, we first filtered the sample to only include wings with complete landmark sets by dropping any landmarks with less than 0.4 posterior probability according to DeepLabCut’s assignment (i.e., landmarks that could not be placed by the algorithm with high probability). The wing images were evenly distributed by sex, selection treatment, and replicate (χ^2^ = 3.47, *df* = 31, P = 1).

We carried out two sets of geometric morphometric analyses designed to test for (1) shape and centroid size differences based on sex, selection treatment, and the sex by selection interaction using only right wings and (2) directional and fluctuating asymmetry of shape and size using paired left and right wings of the same fly.

We used the R packages geomorph (Version 4.1.1; Adams and Otárola-Castillo 2013; Collyer and Adams 2018) and RRPP (Version 2.2.0; Baken et al. 2021) for geometric morphometric analyses, lme4 (Version 2.0-6; Bates et al. 2015), lmerTest (Version 3.2-1; Kuznetsova et al. 2017), and emmeans (Version 2.0.4; Lenth and Piaskowski 2026) for ANOVA-like analyses, and the tidyverse (Version 2.0.0; Wickham et al. 2019) packages as well as readxl (Version 1.5.0; Wickham and Bryan 2026), cowplot (Version 1.2.0; Wilke 2025a), ggrepel (Version 0.9.8; Slowikowski 2026), ggsci (Version 5.2.0; Xiao 2026), ggridges (Version 0.5.7; Wilke 2025b), and gt (Version 1.3.0; Iannone et al. 2026) for data wrangling, plotting, and table creation.

##### Effects of selection and sex on shape and centroid size

Test for the effects of selection (Control and Flight Performance) and sex (Female and Male) on wing shape and size, we selected all individuals with right wings, because more individuals per group had complete right wings (301 right wings vs. 297 left wings; supp. table S11). We performed a Generalized Procrustes Analysis using the geomorph function gpagen() and extracted the resulting centroid size and shape principal components 1-3 for each individual. We used a multilevel model to test for effects of selection, sex, and the selection by sex interaction with replicate population as a random effect using the lmerTest::lmer() function from the lmerTest package which overwrites same function from the lme4 package to implement hypothesis tests using the Satterthwaite approximation of denominator degrees of freedom. To test for multivariate shape differences, we used the lm.rrpp() function from the RRPP package to carry out a MANOVA. Shape was predicted by centroid size, selection, sex, the selection by sex interaction, and the selection by population interaction, the latter to account for replicate lines. Significance was assessed by randomization (9,999 iterations) with Type III sums of squares. Centroid size was included as a covariate because preliminary analysis revealed allometric differences in wing shape. We similarly analyzed the relationship between PC1 and selection, sex, the sex by selection interaction, with centroid size as a covariate and replicate population as a random effect using lmerTest::lmer().

##### Directional and fluctuating asymmetry

To test for directional asymmetry (to answer the question “is there a consistent *left vs. right* bias in size or shape?”) and fluctuating asymmetry (to answer the question “is there non-zero random variation around perfect symmetry at the individual level?”), we used only flies that had both pairs of wings available (n = 66-75 per sex and selection treatment). Flies were divided into four groups by selection (Control and Flight Performance) and sex (Female and Male) and analyzed using the geomorph function bilat.symmetry() function, which follows the conventions of Mardia (2000) and Klingenberg et al. (2002) in parsing shape and size differences into individual-level differences and differences attributable to directional and fluctuating asymmetry using a Generalized Procrustes Analysis and hypothesis tests via randomization. Because landmark digitizing was algorithmic, repeated trials would yield identical landmark locations rather than a traditional measure of digitizing error. Without replicate digitizing trials, we are unable to separate measurement error from true variation among individuals in left vs. right wing differences. With this caveat, we refer to the individual by side parameter in the ANOVA models as “fluctuating asymmetry”, which is reported without a P-value. A secondary analysis of the effects of selection and sex on asymmetry centroid size and shape (following Klingenberg and McIntyre 1998) was carried out using a multilevel model with lmerTest::lmer() as above.

### Life-history traits

#### Preparing populations for assays

We used the same egg collection over two days which generated ‘Set 2’ vials in the Activity assay. We used flies from the first day’s collections to measure survivorship and dry mass, and the second day’s collection for the fecundity measurement. Any vials showing signs of major microbial growth were discarded.

#### Egg to adult survivorship

For each population, we added adults from 15 respective vials to a cage containing food and yeast paste, on day 14 from egg collection. On day 16, we gave an egg collection plate to each cage. To enable easy egg removal, the collection plate was a harder variant than regular food plates, using 2% agar. The formula for 1L of these custom plates was a mixture of 40g yeast, 40g sucrose, 20g agar in 1L water. On day 17, we took out the plates from the cage, and manually removed the eggs onto a thin sheet of 1% agar using a soft brush and water. We counted exactly 70 eggs into 10 replicate vials each. Each vial had approx. 10mL food. On day 14 from egg collection, we transferred the eclosed adults into empty vials and froze them at -20°C for later counting. When counting, we separated flies into male and female groups, counted them and stored them in 1.5 mL centrifuge tubes at -20°C for future dry mass measurements.

For analysis, we arcsine-square-root transformed the proportional egg to adult survivorship values. We fit the following model:

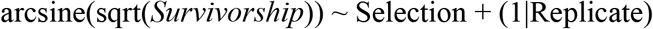

Where the random factor Replicate was the population replicate (1, 2, 3 or 4) nested within Selection (CON or FP).

#### Adult dry mass

Each centrifuge tube for dry mass measurements held groups of either 10 males or 10 females. Prior to weighing the flies, we dried them by keeping the open centrifuge tubes at 70°C for approx. 46-48 hours. Once dried, we weighed the flies on a weighing balance (Mettler Toledo NewClassic MF, model MS304S/03), in duplicate measurements.

For statistical analysis, we fit the following model:

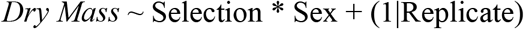

Where the random factor Replicate was the population replicate (1, 2, 3 or 4) nested within Selection (CON or FP). Selection was crossed with Sex (Female or Male). As explained in the results, we further analyzed models for both sexes separately:

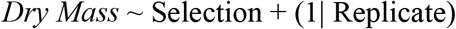

#### Fecundity

We released adults from 10 sampled vials to a cage on day 17 from a respective population’s egg collection. On day 18, we conducted the fecundity assay. We gave each cage 3 food plates successively in 2 hours intervals. Subsequently, we removed each oviposited plate and counted the number of eggs. After we had obtained and counted all 3 plates, we froze the adults in the cage at -20°C overnight. We counted the total number of adults the next day, and used total female count to calculate per capita fecundity for each plate. Prior to analyses, we log-transformed the per capita fecundity to account for a skew in values.

For statistical analysis of log-transformed per capita fecundity, we carried out model comparison using the Akaike Information Criterion (AICc), via the AICcmodavg package in R (ver. 2.2.3; Mazerolle 2023). The following models were compared:

1. Main effect of Selection:

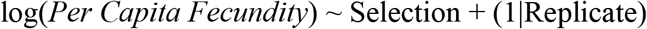
2. Plate number crossed with Selection:

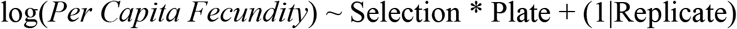

Where the random factor Replicate was the population replicate (1, 2, 3 or 4) nested within Selection (CON or FP). Plate was assigned IDs of 1, 2, 3.

### Note on False Discovery Rate (FDR) correction of *P*-values

We compiled all the *P*-values generated across phenotypic data and applied FDR correction using the Benjamini-Hochberg method (p.adjust function (method = ‘fdr’)). All *P*-values in the Results section are FDR-corrected.

### Pooled whole genome sequencing: preparation, processing and filtering

We sampled female flies from generation 10 of selection for FP_1-4_, and concurrently, female flies from the CON_1-4_ populations. For whole genome extraction, we used the Qiagen Gentra Puregen Cell kit (Qiagen, Hilden, Germany). From each population, we collected a total of 65 females for pooled sequencing; this number was determined by whichever population had the lowest number of female flies sampled (FP_3_). Once extracted, we submitted the DNA samples to the Genomics Technology Core at the University of Missouri. The Core used short-read, paired-end sequencing on an Illumina NovaSeq 6000 platform on the S4 lane, with a targeted library insert size of 550 bp and a paired-end read length of 2×150 bp. The following steps explain the alignment, cleanup and filtering of the pool-seq data.

#### Reference Genome Preparation

We indexed the *Drosophila melanogaster* reference genome (release r6.60) using SAMtools v1.20 faidx and created a sequence dictionary using GATK v4.2.6.1 CreateSequenceDictionary. We partitioned the reference into 64 evenly sized intervals using GATK SplitIntervals with INTERVAL_SUBDIVISION mode to enable parallelized variant calling.

#### Read Alignment

We aligned paired-end reads to the *D. melanogaster* reference genome (r6.60) using Minimap2 v2.28-r1209 (short-read mode, -ax sr) with read group information including sample ID, flowcell, library, lane, and platform. We used SAMtools v1.20 sort to coordinate-sort aligned reads.

#### Duplicate Marking

We identified and marked duplicate reads using GATK v4.2.6.1 MarkDuplicates, assuming query-name sort order. We coordinate-sorted and indexed marked BAMs with SAMtools v1.20. For samples sequenced across multiple lanes or libraries, we merged the sorted BAMs using SAMtools merge prior to variant calling.

#### Variant Calling

We called per-sample variants using GATK v4.2.6.1 HaplotypeCaller in GVCF mode (-ERC GVCF) with ploidy set to 8 (--sample-ploidy 8), run in parallel across 64 genomic intervals. We merged per-interval GVCFs per sample using GATK SortVcf.

#### Joint Genotyping

We imported all per-sample GVCFs into a GenomicsDB datastore using GATK GenomicsDBImport, independently for each genomic interval. We performed joint genotyping across all samples using GATK GenotypeGVCFs. We merged per-interval VCFs into a single cohort VCF using GATK SortVcf.

#### Variant Filtering

We extracted SNPs and indels separately using GATK SelectVariants, and applied hard filters using GATK VariantFiltration with the following thresholds:

**SNPs:** QD < 2.0, QUAL < 30.0, SOR > 3.0, FS > 60.0, MQ < 40.0, MQRankSum < −12.5, ReadPosRankSum < −8.0

**Indels:** QD < 2.0, QUAL < 30.0, FS > 200.0, ReadPosRankSum < −20.0

We flagged variants failing any filter, and retained passing variants using BCFtools v1.20 view -f PASS. We merged the filtered SNP and indel sets into a final cohort VCF using GATK SortVcf.

#### Pipeline Implementation

We implemented the full pipeline in Nextflow v24.04.2 (DSL2) and executed it on a SLURM HPC cluster. The computation for this work was performed on the high-performance computing infrastructure operated by Research Support Solutions in the Division of IT at the University of Missouri, Columbia MO (https://doi.org/10.32469/10355/97710).

#### Filtering based on allele frequency and read depth

We filtered the SNP data based on the following:

1. Read depth between 20 and 500 (any site with any population outside these criteria are filtered). This was largely based around the conventional filtering range, with reasoning supported by an earlier study (Ferretti et al. 2013).
2. Allele frequency: at each SNP, at least one population should have frequency >=0.02 and <=0.98.

With these filters, 1,362,292 SNPs remained. Before any filters, and after selecting relevant chromosomal arms (2L, 2R, 3L, 3R, X) and dropping sites marked NA, the SNPs numbered 1,423,464. A total of 61,172 sites were filtered out, which is approx. 4.3% of the unfiltered data (supp fig. S7A). The average read depth was 102.623 after filtering. We have shown the distribution of read depth after filtering in supp. fig. S7B.

### Principal Component Analysis

After cleanup and filtering, we carried out a PCA on the entire SNP dataset. This enabled a visual estimation of differentiation between all 8 populations. We arranged the reference allele frequency data in a matrix of 8 rows (for 8 populations) and 1,362,292 columns, for each of the loci. We performed PCA using the prcomp function in R, after centering the allele frequency to the respective mean at each locus (center = TRUE), and scaling the allele frequency at each locus to its respective unit variance (scale = TRUE). We generated a biplot of PC1 and PC2, to visualize the genomic structure across the 8 populations. We also measured the % variance captured by each of the principal components in a scree plot.

### SNP Differentiation Analysis

On the filtered SNP table, we arcsine-square-root-transformed the reference allele frequencies. We fit the following linear model:

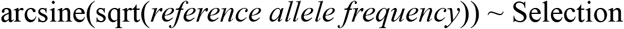

Where Selection had 2 levels, CON and FP. Each CON and FP had 4 replicates (drawn from population replicates 1, 2, 3, 4). We carried out this analysis for each SNP. The generated table of P-values was adjusted for false discovery rate (Benjamini-Hochberg correction) using the p.adjust function (method = ‘fdr’).

### Average Nucleotide Diversity (θ_π_)

We calculated the average nucleotide diversity using Grenedalf v0.6.3 (Czech et al. 2022, 2024). The input parameters were:

“--pool-sizes”, “130”: Pool size of 130 was due to 65 females per population contributing 2 sets of chromosomes each.
“--window-type”, “interval”; “--window-interval-width”, “100000”; “--window-interval-stride”, “100000”: we used a sliding window analysis with an interval width of 100,000 based on earlier recommendations (Da Silva Ribeiro et al. 2022). We kept the interval stride equal to interval width to make the windows non-overlapping (Shahrestani et al. 2021).
“--window-average-policy”, “valid-loci”; “--filter-sample-min-count”, “2”; “--filter-sample-min-read-depth”, “4”: We removed singletons as per the requirement of Grenedalf, the min-read-depth was a dummy value because we had pre-filtered with a minimum of 20x read depth.

### Population differentiation (F_ST_)

We also used Grenedalf (v0.6.3) to calculate population differentiation using F_ST_. We considered both Nei (Nei 1986) and Hudson (Hudson et al. 1992) formulations, with the primary focus on the Hudson formula as per earlier recommendations (Bhatia et al. 2013). The parameters used to calculate F_ST_ were:

“--pool-sizes”, “130”; due to 65 females used.
“--method”, “unbiased-hudson”: this was “unbiased-nei” for the Nei formulation.
“--window-type”, “interval”; “--window-average-policy”, “valid-loci”; “--window-interval-width”, “100000”; “--window-interval-stride”, “100000”: see text for nucleotide diversity.
“--filter-sample-min-count”, “2”; “--filter-sample-min-read-depth”, “4”: see text for nucleotide diversity.

### Haplotype Inference and Analysis

We created our starting population by crossing recombinant inbred lines from the DSPR. The DSPR lines resulted from an 8-way 50-generation intercross followed by 25 generations of full-sib mating to create recombinant inbred lines whose genomes are fine-scale mosaics of the original 8 inbred founder lines (King et al. 2012a,c; Long et al. 2014). Thus, at any given location in the genome, there are 8 possible ancestral haplotypes segregating in our populations. This structure provides an opportunity to detect changes in the frequency of the different haplotypes in our flight performance versus control lines by inferring the 8 haplotype frequencies at each location in the genome from our pool-seq allele frequency data and testing for differences in this set of frequencies between control and selected lines.

We used a likelihood-based method to infer the optimal combination of haplotype frequencies that would produce our observed set of allele frequencies within a sliding 100kb window. We determined the ideal method and window size to use for this inference based on simulations using the DSPR RILs (Supplementary Methods). We have previously used a similar method for inference of the DSPR haplotype frequencies but with a less formalized likelihood calculation (Burke et al. 2014a; Ward et al. 2026), and others have used similar methods in other systems (Kessner et al. 2013; Tilk et al. 2019; Linder et al. 2022; Macdonald et al. 2022). Because many SNPs contribute to this inference, these previous studies have found different estimation methods are able to estimate highly accurate haplotype frequencies, even with modest coverage pooled sequencing data. We describe the development of this likelihood-based method, comparison to our previous method, and the simulations demonstrating the ideal window size for applications using the DSPR in the Supplementary Methods. Based on our simulation results, 100kb is the smallest window size showing high performance in haplotype frequency inference.

Inference of the haplotype frequencies requires the SNP identity of each of the original founder haplotypes for all positions in the window. Thus, we first filtered our pool-seq dataset to contain only those SNPs that were called in the original DSPR founder dataset (King et al. 2012c), resulting in a total of 968,645 positions. To obtain genome-wide haplotype frequency estimates, we used the `optim` function in R to find the set of haplotype frequencies that maximizes the likelihood for the following function for a given SNP window consisting of *nsnp* SNPs:

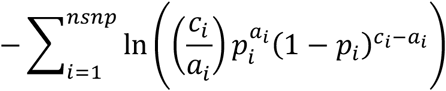

where

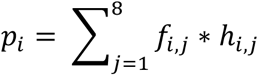

This method uses calculates the likelihood of observing *a_i_* reads (i.e. count of reference allele reads) out of a total of *c_i_* reads (i.e. coverage) at the *ith* SNP using the `dbinom` function in R where the probability is the expected pooled SNP frequency (*p_i_*) calculated based on the set of founder haplotype frequencies (*h_i,j_* is the frequency of the *jth* founder haplotype at the *ith* SNP) and the allele calls for each founder haplotype (*f_i,j_* is the allele call of the *jth* founder haplotype at the *ith* SNP) for a given window containing *nsnp* number of SNPs. The calculated pooled allele frequencies are constrained to be between 0 and 1 and each founder haplotype frequency is also bounded to be between 0 and 1. We then performed this on each window in the genome, as a sliding window with a single SNP step size. On average, a given window spanned ∼0.25cM and contained ∼875 snps. The result is an estimated set of haplotype frequencies for a 100kb window centered on each SNP location for each of our lines.

We performed a genome scan to localize the genomic regions where there are consistent differences in haplotype frequencies between the control and flight performance lines. At each location, we fit a linear model with the arcsine square root transformed haplotype frequency predicted by haplotype id, treatment, and their interaction, as follows:

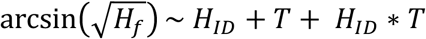

where *H_f_* is haplotype frequency, *H_ID_* is haplotype ID, and *T* is treatment. A significant interaction term indicates a position in which the flight performance and control replicates differ significantly in the set of eight haplotype frequencies. We fit this model at each of our windows in which we estimated haplotype frequencies across the genome. To account for these multiple comparisons, we used the FDR correction for multiple comparisons in R (Benjamini and Hochberg 1995) to obtain adjusted P-values.

### Gene ontology and enrichment analyses

We followed the pipeline for enrichment analyses described by a recent study (Gamboa-Santarosa et al. 2026). We separately annotated the significant SNPs from i) differentiation analysis between CON and FP using the lm function; ii) difference of FP relative to CON from ancestral haplotype composition. We obtained the gene database from TxDb.Dmelanogaster.UCSC.dm6.ensGene v3.12.0 (Bioconductor Core Team and Bioconductor Package Maintainer 2020) and used AnnotationDbi v1.74.0 (Pagès et al. 2026), and GenomicRanges v1.64.0 (Lawrence et al. 2013) to map SNPs to genes. We obtained the gene structure locations for each SNP using the locateVariants function from the VariantAnnotation package v 1.58.0 (Obenchain et al. 2014).We generated Gene Ontology terms from the annotated SNPs using the packages org.Dm.eg.db v3.22.0 (Carlson 2025) and biomaRt v2.68.0 (Durinck et al. 2005, 2009). The following low confidence GO term annotations were removed: Traceable Author Statement (TAS), Non-traceable Author Statement (NAS), Inferred by Curator (IC), No biological Data available (ND). We implemented gene enrichment using clusterProfiler v4.20.0 (Yu et al. 2012; Yu 2024). We used the enrichGO function with the following settings:

OrgDb = org.Dm.eg.db, keyType = “ENSEMBL”: database for *Drosophila melanogaster* genes. The background universe consisted of all *Drosophila melanogaster* genes.

ont = “BP”: GO terms were enriched for the Biological Process aspect (i.e., biological pathways). pAdjustMethod = “fdr”, pvalueCutoff = 0.05, qvalueCutoff = 0.05: *P*-values obtained from enrichment were corrected using the Benjamini-Hochberg method, with an adjusted *P*-value cut-off set at <0.05, and q-value cut-off at <0.05.

Following enrichment, we used the simplify function provided by clusterProfiler to remove redundant pathways. We finally plotted the enriched Gene Ontology terms using the dotplot function in the package enrichplot v1.32.0 (Yu 2026).

## Data Analysis

We used R v. 4.6.1 (R Core Team 2026) and the following R packages: AICcmodavg v. 2.3.4 (Mazerolle 2023), ComplexUpset v. 1.3.3 (Lex et al. 2014; Krassowski 2020), cowplot v. 1.2.0 (Wilke 2025a), effectsize v. 1.0.3 (Ben-Shachar et al. 2020), emmeans v. 2.0.3 (Lenth and Piaskowski 2026), ggrastr v. 1.0.2 (Petukhov et al. 2023), ggrepel v. 0.9.8 (Slowikowski 2026), grateful v. 0.3.0 (Rodriguez-Sanchez and Jackson 2025), gt v. 1.3.0 (Iannone et al. 2026), INLA v. 23.4.24 (Rue et al. 2009; Bakka et al. 2018), lme4 v. 2.0.1 (Bates et al. 2015), lmerTest v. 3.2.1 (Kuznetsova et al. 2017), modelsummary v. 2.6.0 (Arel-Bundock 2022), officer v. 0.7.5 (Gohel et al. 2026), paletteer v. 1.7.0 (Hvitfeldt 2021), pals v. 1.10 (Wright 2025), patchwork v. 1.3.2 (Pedersen 2025), RColorBrewer v. 1.1.3 (Neuwirth 2022), rmarkdown v. 2.31 (Xie et al. 2018; Allaire et al. 2026), tidyverse v. 2.0.0 (Wickham et al. 2019). We have cited additional packages in their relevant methods sections.

## Data availability

The scripts for all results and plots can be found at the GitHub link: https://github.com/EGKingLab/Flying-Flies.git

Raw sequence data is available at NCBI SRA: PRJNA1542687

Variant calling thresholds and filters are defined in supp. tables S23, S24.

Data files are available at Zenodo: 10.5281/zenodo.23356991

## Results

### Evolution of flight performance

#### Flight performance across 10 generations of selection

ANOVA on the fit model revealed a significant main effect of Selection, which also had a large effect size (F_1, 8.19_ = 22.8834, *P* = 0.0036; partial η^2^ = 0.7364). On average, flies from the selected populations (FP) traversed a greater distance than flies from the control populations (CON). Additionally, Generation showed a significant main effect and a large effect size (F_9, 48.02_ = 8.0166, *P* < 0.0001; partial η^2^ = 0.6004). The interaction of Selection and Generation was also significant (F_7, 47.83_ = 4.4916, *P* = 0.0019; partial η^2^ = 0.3966). Figure 2A shows the actual chamber positions of 2500 randomly sampled flies from each Selection level, across generations. We sampled these flies without replacement, pooling population replicates and multiple runs. The entire dataset is available in supp. fig. S4. These plots indicate that flies in the FP populations traversed further than CON flies across generations. Fig. 2B shows the mean chamber traversed by flies of either FP or CON populations, at each generation. We calculated the mean chamber traversed by weighting the chamber number by the proportion of flies found in each chamber (averaged across runs in case of FP). The data show an observable difference between FP and CON at > 6 generations of selection, with the FP populations showing traversing a greater distance in the flight tunnel (fig. 2). As the number of runs as well as the completeness of data differed between CON and FP, we carried out separate analyses for both. The results reflected the observations from fig. 2. When analyzed on its own, FP showed a significant main effect of generation and a very large effect size (F_9, 26.08_ = 12.08, *P* < 0.0001; partial η^2^ = 0.8065). However, CON populations showed no significant main effect of generation, with a large effect size likely due to between-generation fluctuations (F_7, 24_ = 1.0325, *P* = 0.5415; partial η^2^ = 0.2314). The ANOVA tables are available in supp. table S2, and effect size tables in supp. table S3.

**Figure 2:**
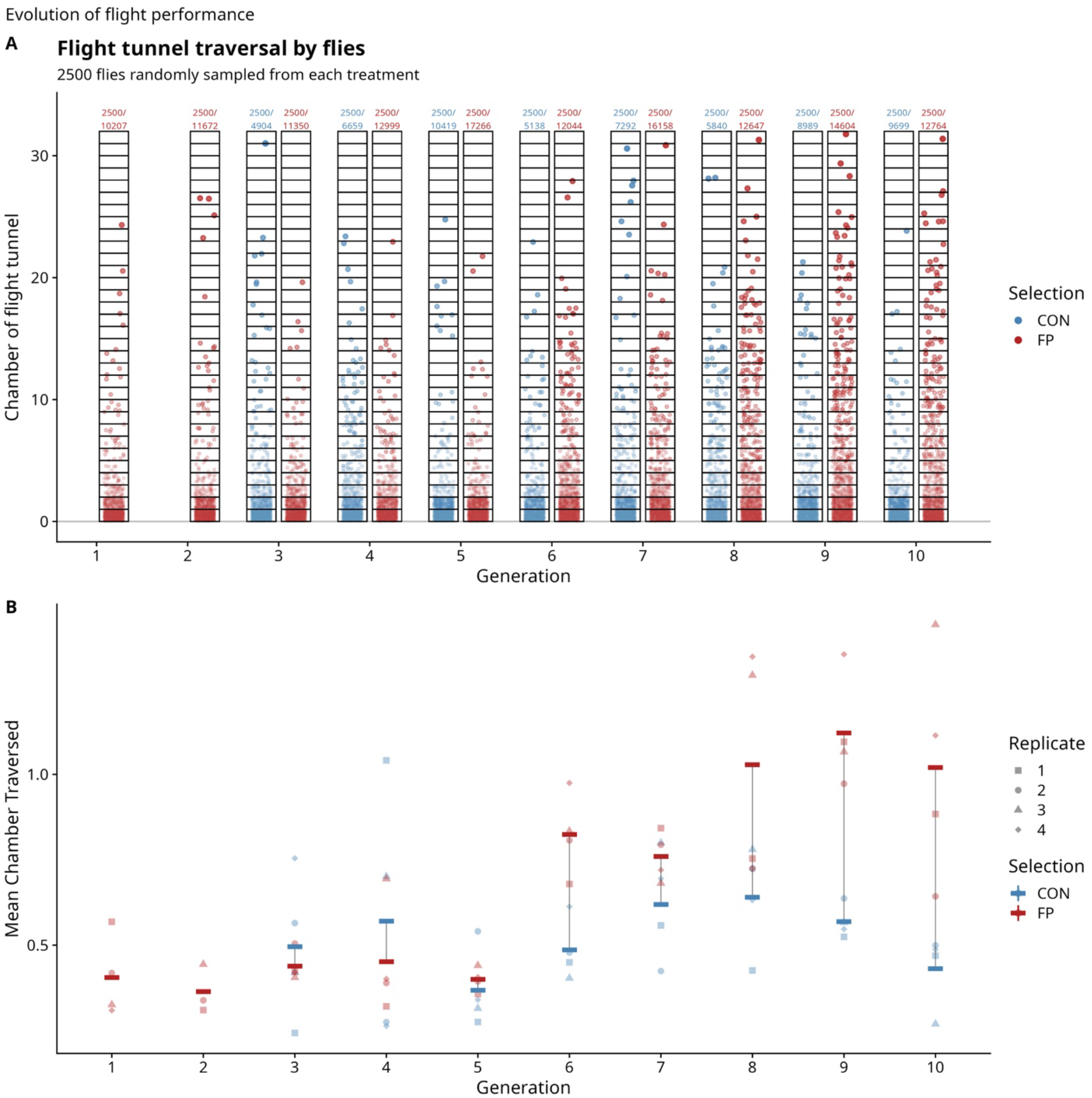
The evolution of flight performance across 10 generations of selection. A. The distance traversed by flies within the flight tunnel, pooling across replicate populations and runs. The X-axis shows generation number of selection. The Y-axis denotes the chamber position for 2500 randomly sampled flies (without replacement) for each combination of Selection × Generation. Above each scatter, the “2500/n” denotes the sampling of 2500 from a pool of n flies. For the complete dataset of n flies, see supp. fig. S4. B. Mean flight performance across 10 generations of selection. The X-axis shows generation number of selection, the Y-axis shows mean chamber traversed, which is an indication of flight performance in the wind tunnel. We calculated the mean chamber by taking the average chamber number, weighted by the proportion of flies found in each chamber (averaged across runs in case of FP). Point shape denotes replicate population. Generations 1 and 2 of selection did not have the respective flight performance data for CON. Analysis showed that there was no detectable difference between CON flight performance across generations. FP flight performance significantly improved between generations 1 and 10.

#### Factorial flight performance assay

Model comparison revealed that the purely additive model incorporating the main effects of selection, light status and fan status had the maximum support for mean distance (*Mean Chambers Traversed*) traversed within the wind tunnel (AICc weight 0.8071; supp. table S4):

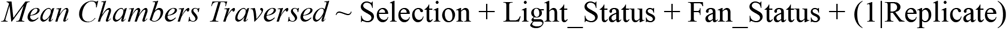

Where the random factor Replicate was the population replicate (1, 2, 3 or 4) nested within Selection. *Mean Chambers Traversed* was the measure of how far flies travelled on average in the tunnel. We calculated it as the mean chamber number weighted by the proportion of flies found in each chamber, per run. The results altogether indicate a major effect of selection on mean distance travelled, with flies from selected FP populations traversing, on average, significantly more than those from CON (F_1, 6_ = 13.9243, *P* = 0.0214; partial η^2^= 0.6989; fig. 3A). Light status showed a moderate effect size on mean distance traversed (partial η^2^= 0.0816), with ‘light ON’ inducing a non-significant pattern of greater mean distance traversed than ‘light OFF’ (F_1, 54_ = 4.7997, *P* = 0.0577; fig. 3A). The effect size for fan status was moderate, similar to light status (partial η^2^= 0.0862). Flies on average showed a non-significant pattern of travelling a greater distance when the fan was ‘OFF’ than when it was ‘ON’ (F_1, 54_ = 5.0947, *P* = 0.0529; fig. 3A). The AIC comparison did not show support for any model with interactions, indicating that interactions were unlikely to have enough information to include their extra parameters into the fitted model (supp. table S4). We have shown the complete factorial framework for mean chamber traversed by flies in fig. 3B. The ANOVA tables are available in supp. table S5, and effect size tables in supp. table S6.

**Figure 3:**
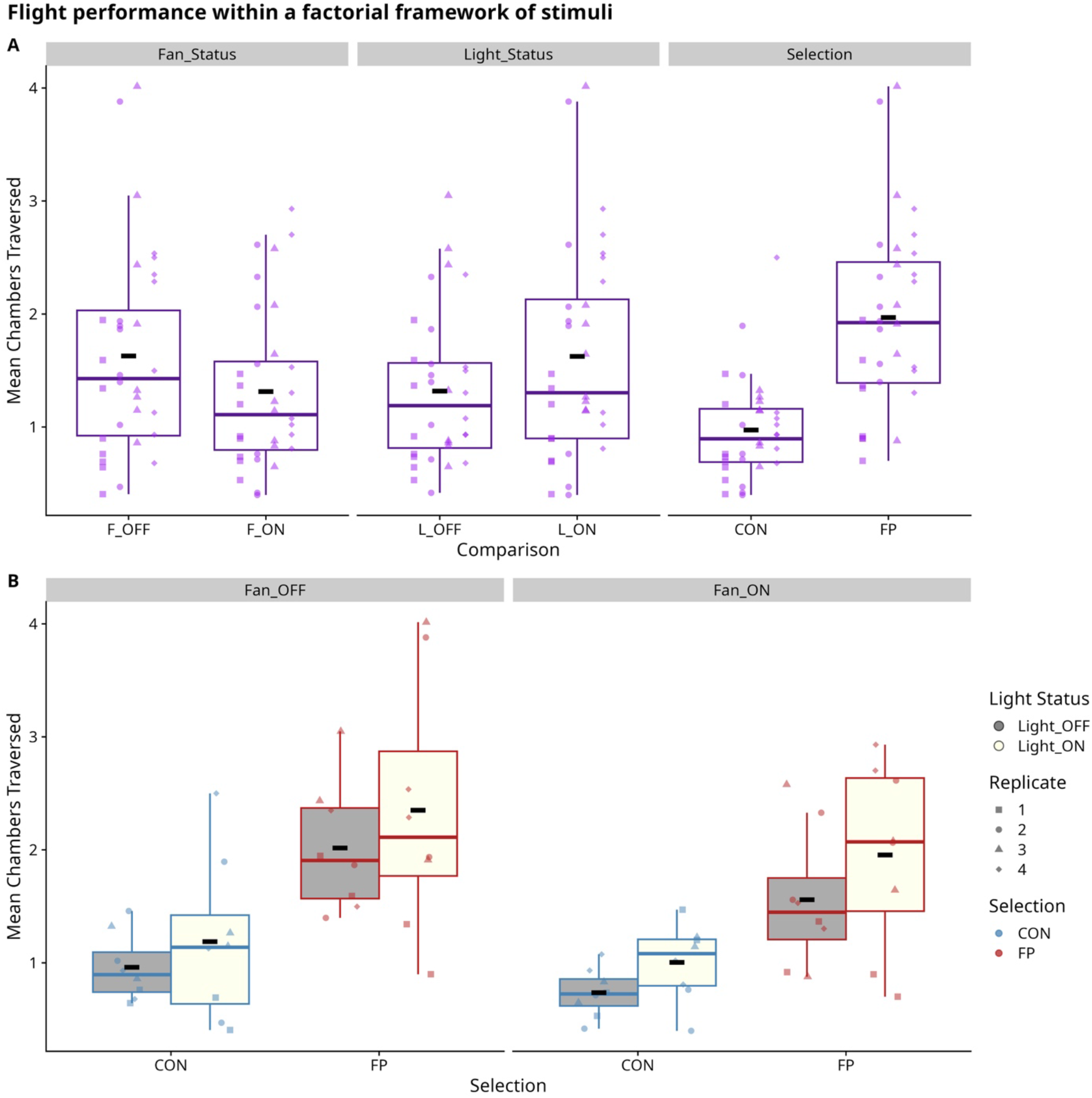
Measuring the effect of factorial framework of stimuli on the evolution of flight performance in selected (FP) populations compared to control (CON) populations. In this assay, for each run, fan and light were each in one of two configurations (ON or OFF), in a factorial design. In both plots, the Y-axis denotes mean flight performance represented as the mean chamber traversed by flies. We calculated this using the weighted average of the number of flies in each chamber. In both plots, point shape denotes population replicate. A. Main effects of Selection, Light or Fan as factors on mean flight performance in the assay. The facets show mean flight performance for a given factor, averaging across other factors. The X-axis across facets shows the comparisons between: Fan status, Fan OFF (F_OFF) vs. Fan ON (F_ON); Light status, Light OFF (L_OFF) vs. Light ON (L_ON); Selection, FP vs. CON. The black ‘-’ within each box plot shows the overall mean across runs and population replicates. The main effects showed significant effects for each of Selection (FP > CON), Light status (L_OFF < L_ON) and Fan status (F_OFF > F_ON). B. Mean flight performance for all the factors plotted together. The X-axis shows Selection (FP vs. CON), the plots are facetted by Fan status, and the boxplots colors are filled according to Light status. The black ‘-’ within each box plot shows the overall mean across runs and population replicates.

### Flight-related traits

#### Activity levels

The results of AICc showed the strongest support for the additive model with Selection and Sex, with Assay_set as a fixed factor crossed with Selection (AICc weight = 0.5518; supp. table S7):

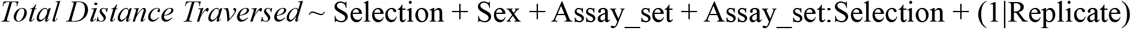

The model with the second highest support was the full cross of Selection and Sex, with Assay_set as a fixed factor crossed with Selection (AICc weight = 0.3599; supp. table S7):

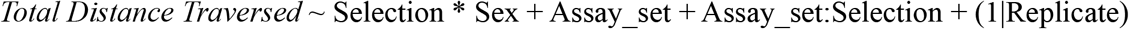

In all models being compared, random factor Replicate was the population replicate (1, 2, 3, or 4) nested within Selection (CON or FP). Sex was a fixed factor (Female or Male). Assay_set (Set1 or Set2) represented the two different sampling events we carried out, on different days of the experiment. We carried out an ANOVA on the model with the cross between Selection and Sex, because it contained all the terms of the additive model. Results showed that Selection had a very large effect size (partial η^2^ = 0.5905), with a marginally significant pattern of flies of the FP populations on average traversing greater distances than flies of CON populations (F_1,6_ = 8.5665, *P* = 0.0518). The ANOVA also revealed a significant main effect of Sex, with a moderate effect size (F_1,1414_ = 233.2784, *P* < 0.0001; partial η^2^ = 0.1416), with male flies on average having greater activity than female flies (fig. 4A). The opposite directions of significance value and effect size for Selection and Sex are likely due to differences in degrees of freedom in the denominators used for calculating respective F values. The interaction effect of Selection and Sex was not significant and had a negligible effect size (partial η^2^ = 0.0008).

**Figure 4:**
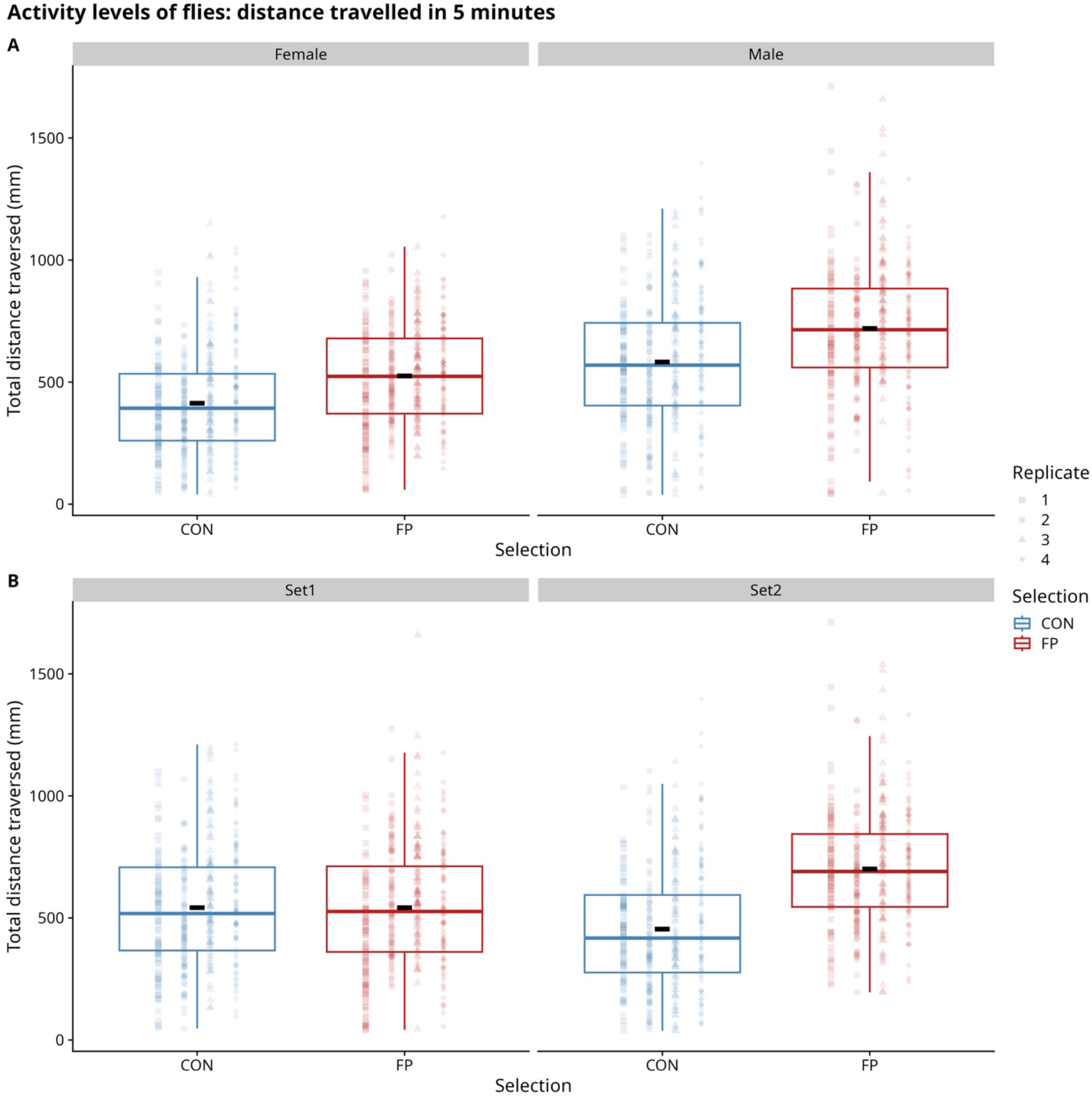
Distance traversed in the activity measurement assay. In both plots, the Y-axis shows the cumulative distance traversed by each fly in the activity tube over 5 minutes. The black ‘-’ within each box plot shows the overall mean across runs and population replicates. A. The effects of both Selection (X-axis) and Sex (facets). B. The effects of both Selection (X-axis) and assay set (facets). The results show that the second assay (Set 2) showed differences between CON and FP activity but not Set 1 (supp. tables S8, S10).

Interestingly, the two different assay sets also displayed statistically significant main effect (F_1,1414_ = 8.7292, *P* = 0.0078; partial η^2^ = 0.0061) and a significant interaction of assay set with Selection (F_1,1414_ = 105.2515, *P* < 0.0001; partial η^2^ = 0.0693). While significant, these terms displayed relatively small effect sizes.

A post-hoc analysis using Tukey’s HSD revealed that the total distance traversed by FP (selected) flies was significantly greater than the distance traversed by CON (control) flies only in the second experiment (CON Set2 – FP Set2; t.ratio = -5.6141, d.f. = 6.97 (Kenward-Roger), *P* = 0.0082; fig. 4B). We did not see a significant difference in the first experiment (CON Set1 = FP Set1; t.ratio = -0.0498, d.f. = 6.97 (Kenward-Roger), *P* = 1.0; fig. 4B). The AICc results are available in supp. table S7, ANOVA results in supp. table S8, effect size tables in supp. table S9, and Tukey’s HSD results in supp. table S10.

#### Wing morphometrics

##### Effects of sex and selection on shape and centroid size

Geometric morphometric revealed differences in centroid size and shape for both sex and selection effects. The two-way factorial multilevel model *Centroid Size ∼ Selection × Sex + (1 | Replicate)* showed that wings of female flies are, on average, larger than those of male flies, and selection for flight performance resulted in increased centroid size (i.e., larger wings) in both sexes (Tables 1, 2; Figure 5a). The selection effect is mildly modulated by sex (P = 0.0413), indicating a differential response in centroid size to selection. Although interpretation of main effects in the presence of a significant interaction can be problematic, we feel that the interaction effect is relatively weak and thus remain confident in the main effects.

**Figure 5.**
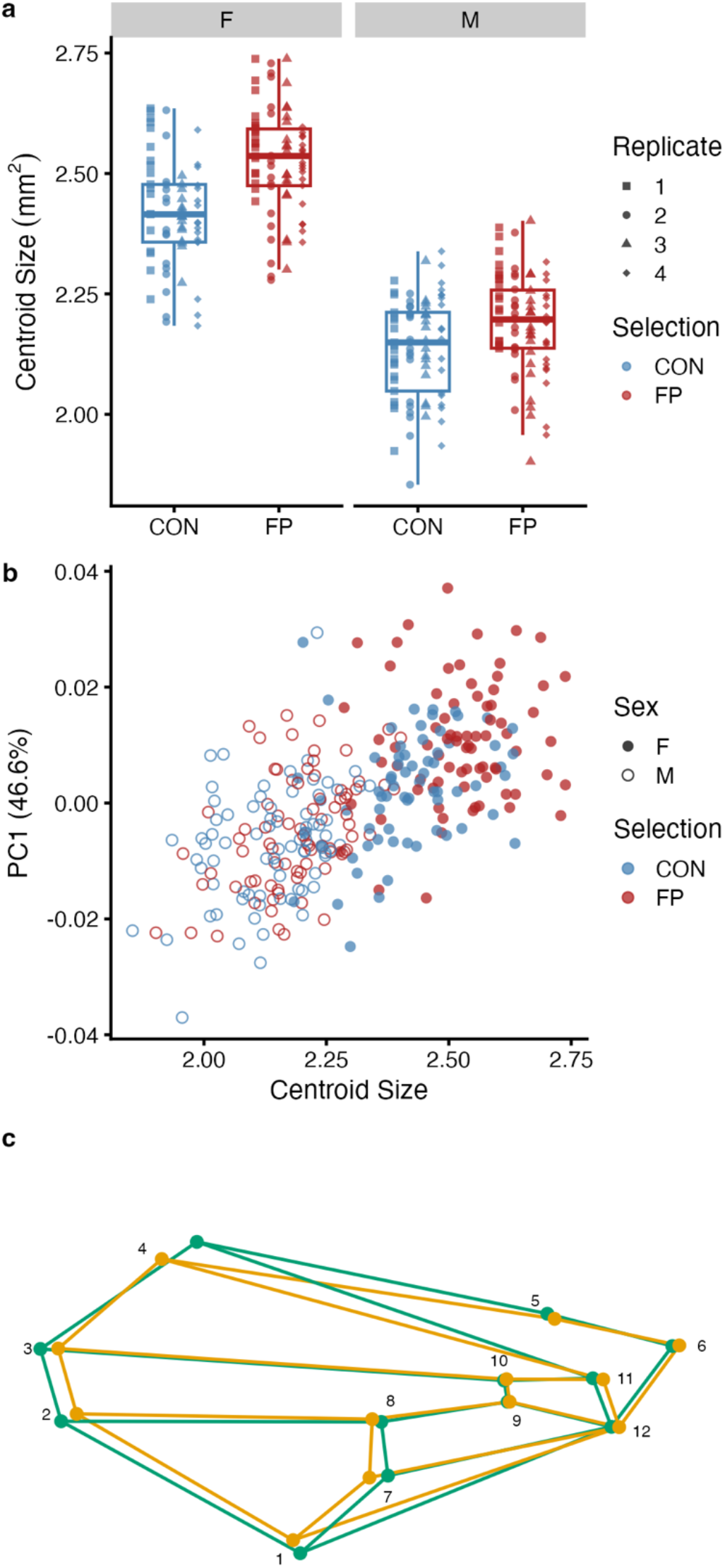
(a) Wing centroid size in replicate flight selected (SEL) and control (CON) flies. Across all replicates, females have larger wings and selection for flight performance resulted in increased wing centroid size. (b) PC1 plotted against centroid size. Females (closed circles) are generally larger than males (open circles), but the effects of selection for flight performance are less clear, with substantial overlap in SEL and CON individuals. (c) Wireframe comparing the shapes of the minimum (green) and maximum (orange) values along the PC1 axis. Males generally have relatively long and narrow wings, while females have relatively short and broad wings.

**Table 1:** Least-squares means and 95% confidence intervals of wing centroid size.

| Sex | Selection | Centroid Size ($mm^2$) | SE | df | Lower CI | Upper CI |
| --- | --- | --- | --- | --- | --- | --- |
| F | CON | 2.417 | 0.017 | 12.38 | 2.381 | 2.453 |
| F | FP | 2.526 | 0.016 | 11.14 | 2.490 | 2.561 |
| M | CON | 2.137 | 0.016 | 10.98 | 2.102 | 2.172 |
| M | FP | 2.190 | 0.016 | 10.69 | 2.155 | 2.225 |

**Table 2:** Results of the multilevel ANOVA model *Centroid Size* ∼ Selection + Sex + Selection:Sex + (1 | Replicate)

| Effect | SS | MS | df<sub>num</sub> | df<sub>den</sub> | F | P-value |
| --- | --- | --- | --- | --- | --- | --- |
| Selection | 0.18 | 0.18 | 1 | 6.1 | 17.13 | 0.0132 |
| Sex | 7.12 | 7.12 | 1 | 291.4 | 671.39 | <<0.0001 |
| Selection × Sex | 0.06 | 0.06 | 1 | 291.4 | 5.47 | 0.0413 |

Multivariate analysis of shape with centroid size as a covariate showed significant allometric (i.e., centroid size associated) differences in shape (P = 0.0003; Table 3). After accounting for allometric differences, wing shape differed significantly by both sex (P = 0.0383) and selection (P = 0.0003) with significant differences among replicate lines (P = 0.0003). The selection-by-sex interaction term was not significant (P = 0.4166; Table 3).

**Table 3:** Results of the MANOVA model for multivariate shape: *Wing Shape Coordinates ∼ Centroid Size + Selection + Sex + Selection:Sex + Selection:Replicate*. Sums of squares are type III, and P-values are determined empirically, based on 9,999 iterations.

| *Effect* | *df* | *SS* | *MS* | *R*<sup>2</sup> | *F* | *Z* | *P-value* |
| --- | --- | --- | --- | --- | --- | --- | --- |
| Centroid Size | 1 | 0.00473 | 0.00473 | 0.026 | 9.36 | 5.58 | 0.0003 |
| Selection | 1 | 0.00381 | 0.00381 | 0.021 | 7.54 | 5.12 | 0.0003 |
| Sex | 1 | 0.00114 | 0.00114 | 0.006 | 2.26 | 2.05 | 0.0383 |
| Selection x Sex | 1 | 0.00058 | 0.00058 | 0.003 | 1.15 | 0.51 | 0.4166 |
| Selection x Replicate | 6 | 0.01047 | 0.00175 | 0.058 | 3.45 | 6.82 | 0.0003 |
| Residuals | 290 | 0.14658 | 0.00051 | 0.808 |  |  |  |

Analysis of PC1 predicted by selection, sex, and the selection-by-sex interaction with centroid size as a covariate and replicate as a random effect revealed significant allometry of PC1 (P = 0.0003; Figure 5b; Table 4). In contrast to the overall MANOVA shape analysis above, neither sex nor selection was significant in this analysis, likely because RRPP uses all the multivariate shape information. For visualization purposes (Figure 5b), plotting PC1 vs. centroid size shows the allometric differences along with wireframes (Figure 5c) emphasizing the differences in wing shape between males (low PC1 values; green outline) and females (high PC1 values; orange outline). Males tend to have higher aspect ratio wings (relatively long and narrow), while females have lower aspect ratio wings (relatively short and broad anteroposteriorly).

**Table 4:** Results of the multilevel ANONA model: PC1 ∼ Selection + Sex + Selection:Sex + Centroid Size + (1 | Replicate)

| *Term* | *Estimate* | *SE* | *df* | *t* | *P-value* |
| --- | --- | --- | --- | --- | --- |
| Intercept | -0.0725 | 0.0128 | 292.8 | -5.65 | <<0.0001 |
| Selection (FP) | 0.0047 | 0.0020 | 17.0 | 2.38 | 0.0537 |
| Sex (M) | -0.0011 | 0.0022 | 295.0 | -0.51 | 0.7341 |
| Centroid Size | 0.0310 | 0.0053 | 294.9 | 5.87 | <<0.0001 |
| Selection × Sex | -0.0039 | 0.0022 | 290.6 | -1.76 | 0.1319 |

##### Directional and fluctuating asymmetry

As expected, we found widespread differences in wing centroid size among flies within groups, (Table S12; “Individual Variation” row; P = 0.0003), which represents fly to fly variation. We found no evidence for directional asymmetry in wing centroid size (P range from 0.4837 to 1). Due to the automated wing landmarking protocol, we were unable to directly test for fluctuating asymmetry in centroid size. However, the R^2^ for individual variation in above 0.98 for all groups, with a proportionately small amount (∼1%) of the variation attributable to fluctuating asymmetry (Table S12; R^2^ = 0.007 to 0.016). In contrast, we do find evidence for significant directional asymmetry in wing shape in two of the four groups (Table S13): female control line flies (P = 0.0037) and male flight performance flies (P = 0.0041), although we note that the proportion of variance explained by each is relatively small (R^2^ = 0.009 in both groups). Pooled multilevel analyses of all groups were used to test for effects of selection and sex on directional asymmetry in centroid size and fluctuating asymmetry in centroid size and multivariate shape. Overall, we found no significant differences between treatments or sexes (Table S14).

### Life-history traits

#### Egg to adult survivorship

Performing an ANOVA on arcsine-square-root-transformed survivorship revealed no significant effect of selection (F_1, 5.829_ = 0.0414, *P* = 0.9184), with both control and flight performance selected populations showing similar mean egg to adult survivorship of around 88% (fig. 6B). The effect size for Selection was also negligibly small (partial η^2^ = 0.007). The ANOVA table is available in supp. table S15, and effect size table in supp. table S16.

**Figure 6:**
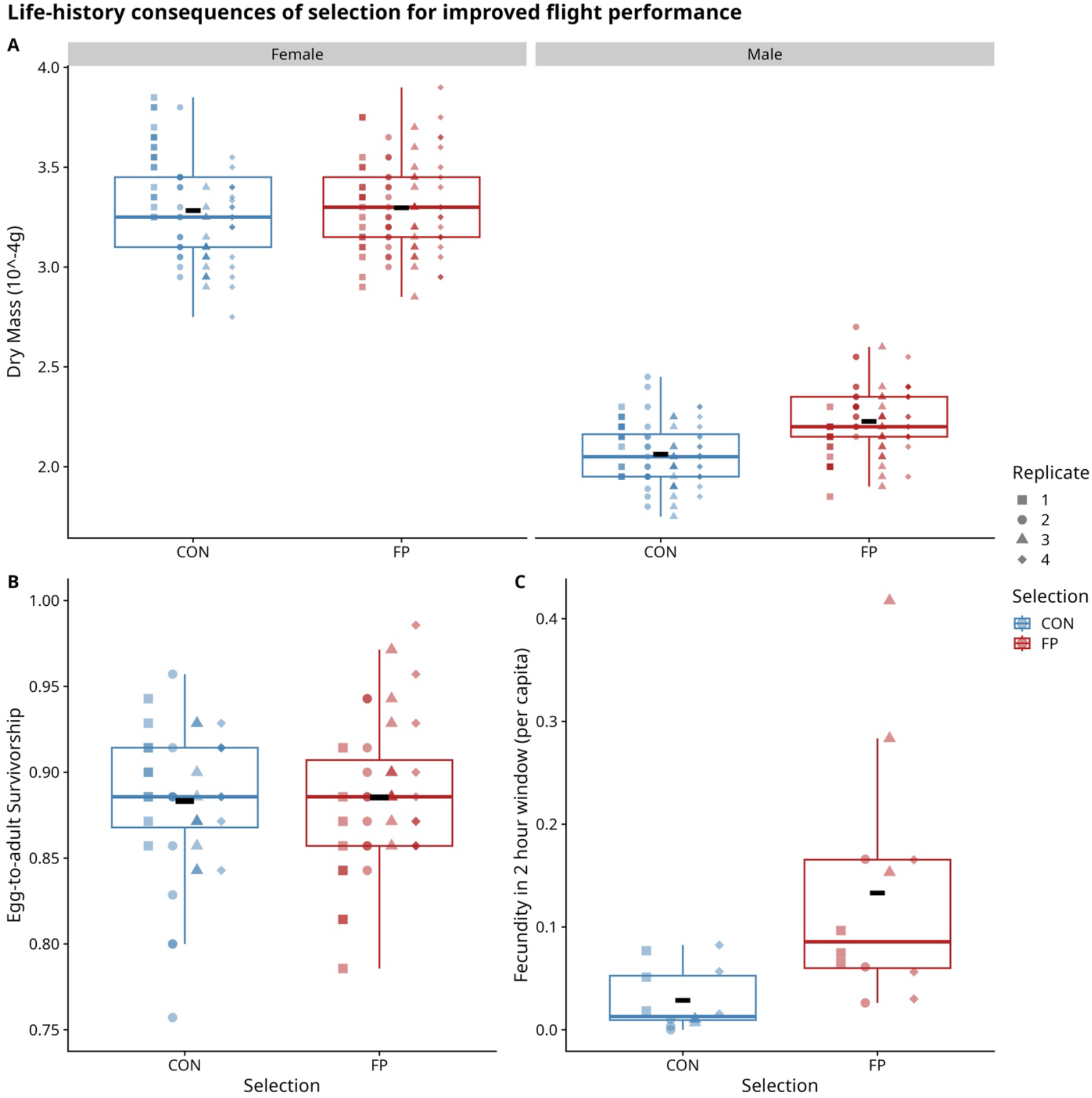
Measuring the life-history consequences of selection for improved flight performance. Across all plots, the X-axis shows Selection, either CON (control) or FP (selected). The shape of the point represents population replicate. The black ‘-’ within each box plot represents the group mean. A. Adult dry mass. The panels are separated by sex. Y-axis shows per capita dry mass (10^-4^g). While we found no significant difference in per-capita dry mass, selection had a large effect size for male flies, and a pattern of FP>CON could be observed. B. Egg to adult survivorship (proportion eggs becoming adults). The Y-axis shows proportion survival of eggs to adults. No significant difference was found between CON and FP. C. Fecundity measured within 2-hour windows. Each population’s fecundity was measured over 3 consecutive 2-hour windows. A per capita measure of fecundity was calculated from this, shown in the Y-axis. While we found no significant difference in per-capita fecundity, selection had a large effect size, and a pattern of FP>CON could be observed.

#### Dry Mass of adults

When both selection and sex were analyzed together, there was very large effect size for sex (partial η^2^ = 0.9099) due to the consistently large difference in mass of females and males. Thus, both female and male dry mass data were analyzed separately for a main effect of selection and population replicate as a random factor nested within the Selection factor. Separate analyses revealed no significant effect of Selection in females, with a negligible effect size (F_1, 6_ = 0.0217, *P* = 0.9451; partial η^2^ = 0.0004; fig. 6A). For males, a separate analysis revealed a very large effect size for the main effect of Selection (partial η^2^ = 0.59), which was marginally non-significant (F_1, 6_ = 8.4801, *P* = 0.0522; fig. 6A). Male flies from the selected (FP) populations showed a non-significant pattern of greater mean dry mass than flies from control (CON) populations (fig. 6A). The ANOVA tables are available in supp. table S17, and effect size table in supp. table S18.

#### Fecundity

The results of AICc showed the strongest support for the model fitting only the main effect of Selection (AICc weight = 0.9932; supp. table S19):

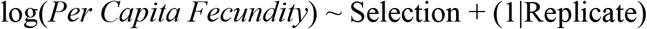

We saw a large effect size for the main effect of Selection on log transformed per capita fecundity (partial η^2^ = 0.4450) although the confidence intervals were very large (supp. table S21). The main effect of Selection was not significant (F_1, 6_ = 4.8112, *P* = 0.1197; supp. table S20). A non-significant pattern of difference was observed, with flies from FP populations have greater per-capita fecundity compared to flies from CON populations (fig. 6C). The number of females in the population was not significantly correlated with fecundity of the population, either in CON or in FP (supp. table S22). The number of females ranged from 194 to 321 (supp fig. S6A). We have shown the per capita fecundity counts by plate number in supp. fig. S6B.

ANOVA results for both transformed and untransformed per capita fecundity are in supp. table S20. Effect sizes for the same are in supp. table S21.

### Whole genome analyses

#### PCA

The biplot for the PCA showed overall differentiation between populations, as observed by the distances between different population points (supp fig. S8B). The scree plot (supp fig. S8A) shows that PC1 captured 22.37% variation, PC2 and PC3 captured variation within 1% of each other (16.68% and 15.89%, respectively).

#### SNP differentiation

A total of 42 SNPs were significantly different between the control and selected populations using the linear model analyses, with a post-hoc P-adjustment. These sites are displayed on a Manhattan Plot, plotted above the red line representing the significance threshold (fig. 7A). Looking at the allele frequencies of those significant SNPs, a majority lie in chromosome arm 3R. One significant SNP lies in chromosome arm 2R, and 4 significant SNPs lie in chromosome X. In all chromosomes but X, the average allele frequencies go closer to fixation in the selected populations compared to the control populations (fig. 7B). The read depth for these SNPs can be found in supp. fig. S9A, with the overall relationship between allele frequency and read depth plotted in supp. fig. S9B.

**Figure 7:**
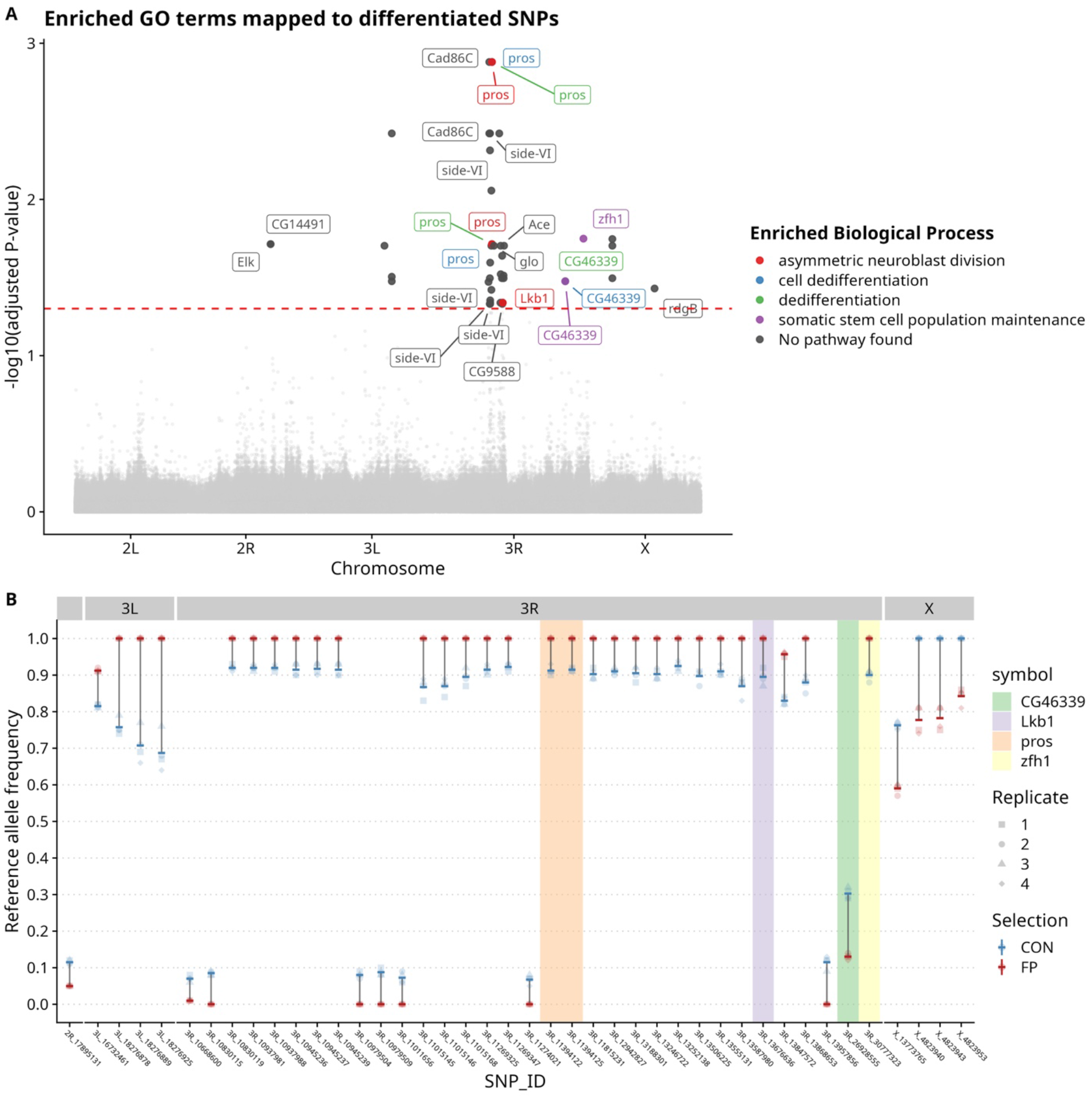
Linear model analysis of pool-seq whole genome data. A. Manhattan plot of SNP-wise *P*-values for comparisons between control (CON) and selected (FP) populations, with mapped genes labelled and colored by enriched biological process. The X-axis shows the SNP positions, divided by chromosomal arms. The Y-axis shows -log10(FDR adjusted *P*-values). The dashed red line represents the cut-off -log10(*P* = 0.05). A total of 42 sites clear these cut-offs, primarily on chromosome 3. These sites were enriched for 4 biological pathways (supp. fig. S13A). Any significantly differentiated SNP mapping to a gene associated with an enriched pathway is color-coded. Genes associated with multiple enriched pathways are labelled each time for the given pathway. Significantly differentiated SNPs within genes not associated with any pathway are colored in dark grey. B. Allele frequencies for SNPs found significant in the linear model analysis between control (CON) and selected (FP) populations, shown above in (A). The X-axis shows significant SNPs by chromosomal arms, with ID labelled. The un-labelled panel on the left represents the single significant SNP in arm 2R. The Y-axis represents reference allele frequencies. The opaque ‘-’ represents the mean allele frequency for the selection factor (CON (control) or FP (selected)). These are connected by a dark grey line to represent the magnitude of difference. The translucent points represent individual allele frequency for each population, with shape representing population replicate ID. SNPs mapped to genes associated with enriched pathways are denoted by distinct background colors.

#### Average Nucleotide Diversity (θ_π)_

Both control (CON) and selected (FP) populations had similar distributions of nucleotide diversity values (supp. fig. S10). A linear model with the following model was fit for every window:

θ_π_ ∼ Selection

The P-values were adjusted using the Benjamini-Hochberg method (p.adjust function, method ‘fdr’). We found no windows of significantly different average nucleotide diversity.

Figure 8A shows the log2 transformed θ_π_ ratio of FP/CON across genomic windows, with 1% outliers highlighted in green. These outlier containing windows did not overlap with windows containing significant SNPs from the linear model test (grey background lines, fig 8A).

**Figure 8:**
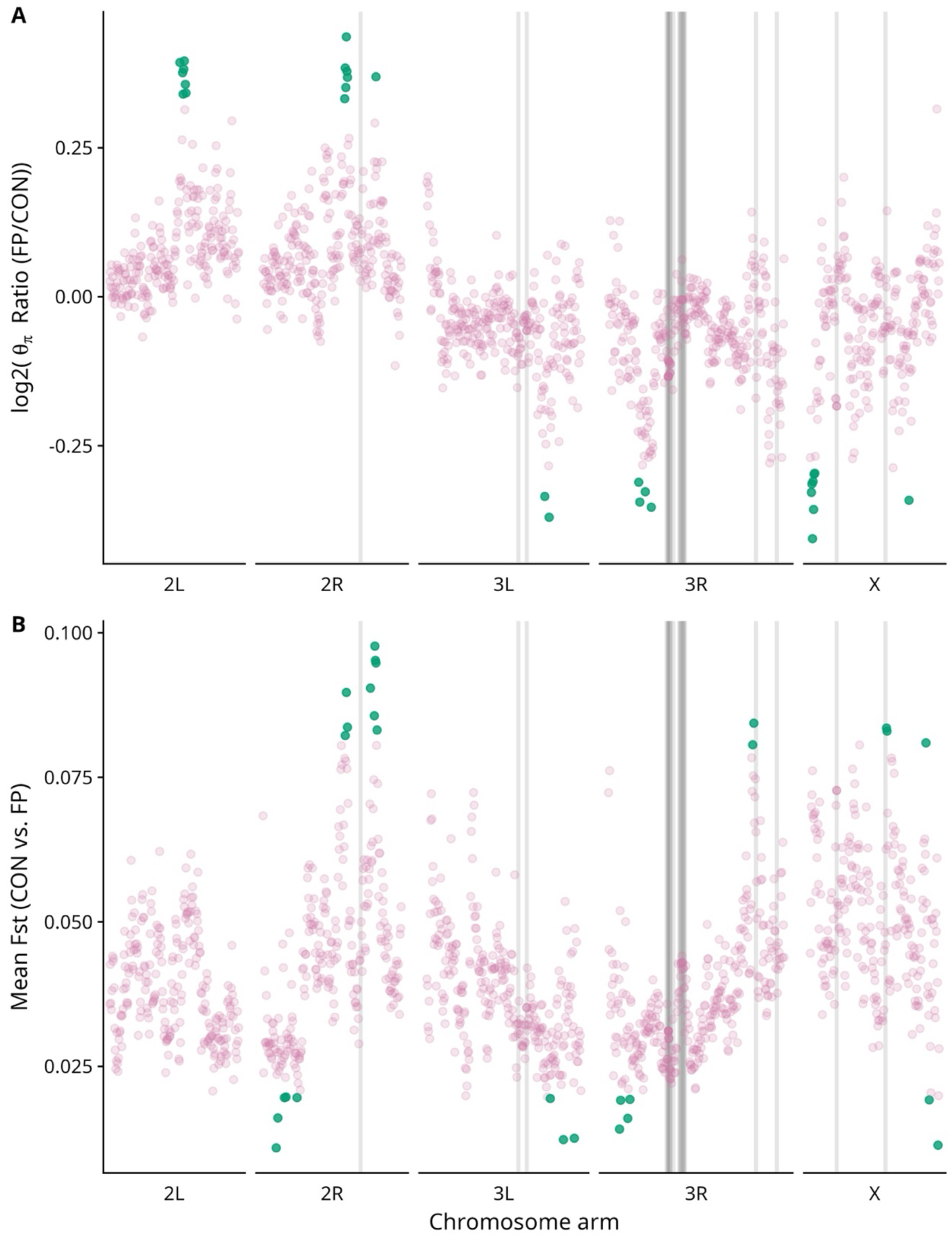
Measures of genetic diversity and population differentiation. A. Comparison of average nucleotide diversity (θ_π_) between selected (FP) and control (CON) populations. The X-axis shows window start position across chromosomal arms, where each window represents the non-overlapping interval of 100kb used for θ_π_ calculation. The Y-axis represents the log2 θ_π_ ratio of FP/CON. Ratio points at extreme 1% are colored in green. The grey lines in the background represent those windows which contains significant SNP sites found in the linear model. No outlier window for θ_π_ ratio overlaps with a window containing a significant SNP. B. Mean F_ST_ calculated between the CON and FP populations. Similar to the genetic diversity plot above, the X-axis represents starting points for 100kb windows. The Y-axis denotes mean F_ST_ (Hudson), calculated between the CON and FP populations. The green points are those which lie at the 1% extremes of F_ST_. The grey lines, as in the above plot, indicate those windows which would contain the significant SNPs from the linear model analysis. No overlaps exist between outlier F_ST_ values and windows containing significantly differentiated SNPs.

#### Population differentiation (F_ST_)

Using both Hudson and Nei estimates showed similar patterns of mean F_ST_ comparisons between all populations (supp. fig. S11, S12), with Nei estimates having lower values than Hudson estimates. For all other plots, F_ST_ was calculated using the Hudson estimate. Classifying by comparisons, CON vs. CON populations had a mean F_ST_ of 0.035, while CON vs. FP (0.041) and FP vs. FP (0.041) comparisons were marginally higher. Supp. fig. S11 shows the distributions of F_ST_ values across comparison types, for both Hudson and Nei estimates. Figure 8B shows F_ST_ values for CON vs. FP over the genomic windows, with 1% outliers marked in green. A few outlier points overlap with nucleotide diversity outliers. However, there were no overlaps between windows containing outlier F_ST_ values and windows containing significant SNP values from the linear model test (grey background lines, fig. 8B).

#### Analyses of Haplotypes

For the comparisons between the control (CON) and selected (FP) populations with respect to their founder haplotype compositions, we found a total of 119,771 sites which were significantly different. We have represented these changes across supplementary figures S14-S18, for each chromosome arm (2L, 2R, 3L, 3R, X) respectively. In each figure (S14-S18), the plot A shows the actual standardized haplotype frequency ascribed to each founder line, for each population, across genomic sites. Plot A contains 8 panels, representing each founder haplotype (A1-A8). The bottom panel displays the standardized reference allele frequency for the given population and genomic position. A dark grey background line denotes sites found to differ significantly between FP and CON populations. A yellow background line is also shown for those sites found different in the linear model test (fig. 7). For each significant genomic position, we calculated the average difference between the CON and FP populations for the frequency of each founder. We ranked each founder by magnitude of difference at each site, with the greatest magnitude of differences ranked 1, and the smallest magnitude ranked 8. Thus, on each chromosomal arm, we could calculate how many times each founder was rank 1, how many times it was rank 2 and so on. Below, we summarize the top 3 ranking founders for each chromosomal arm:

i. Arm 2L (supp. fig. S14B): the founder A4 accounted for the greatest difference (rank 1) for all the significant genomic sites, with a mean rank of 1. Between them, founders A1 (mean rank 2.52) and A5 (mean rank 3.05) accounted for the majority of both rank 2 and 3 differences.
ii. Arm 2R (supp. fig. S15B): the top ranks in significant sites were split between multiple founders: A7 (mean rank 3.57), A4 (mean rank 3.92), A6 (mean rank 3.99), and A8 (mean rank 4.02).
iii. Arm 3L (supp. fig. S16B): founder A4 accounted for the most rank 1 differences (mean rank 2.79). Another notable founder contributing to significantly different sites was A8 (mean rank 3.08).
iv. Arm 3R (supp. fig. S17B): founder A4 accounted for the most rank 1 differences, and had a mean rank of 1.59. Founder A5 accounted for the greatest number of rank 2 differences (mean rank 3.28). Although most of the significantly differentiated SNPs from the linear model test resided in chromosome 3R (fig. 7), there were few overlaps with the sites found significantly different in the haplotype analysis (supp. fig. S17A). None of the overlapping sites mapped to genes involved in enriched biological pathways (fig. 7, supp. fig. S17C).
v. Chromosome X (supp. fig. S18B): founder A6 accounted for the greatest number of rank 1 differences (mean rank 3.19), while founder A7 accounted for the greatest number of rank 2 differences (mean rank 2.83). Founder A4 was also a notable contributor to many significantly different sites (mean rank 3.06).

Across supplementary figures S14-S18, plot C highlights the significant genomic sites mapped to genes involved in enriched biological processes (supp. fig. S13B).

We have further magnified a region in chromosome arm 2R containing significantly different sites mapped to two potentially important genes: *Sdc* and *Egfr* (fig. 9, see Gene Ontology and Enrichment analyses below). The plots of founder haplotype frequencies for CON and FP populations (fig. 9A) show that in this region, founder A6 contributed to the most differences (mean rank 1.01, fig. 9B), followed by founder A1 (mean rank 2.08, fig. 9B) and founder A5 (mean rank 3.36, fig. 9B). The significant sites and their mapped genes in this region are shown in figure 9C.

**Figure 9:**
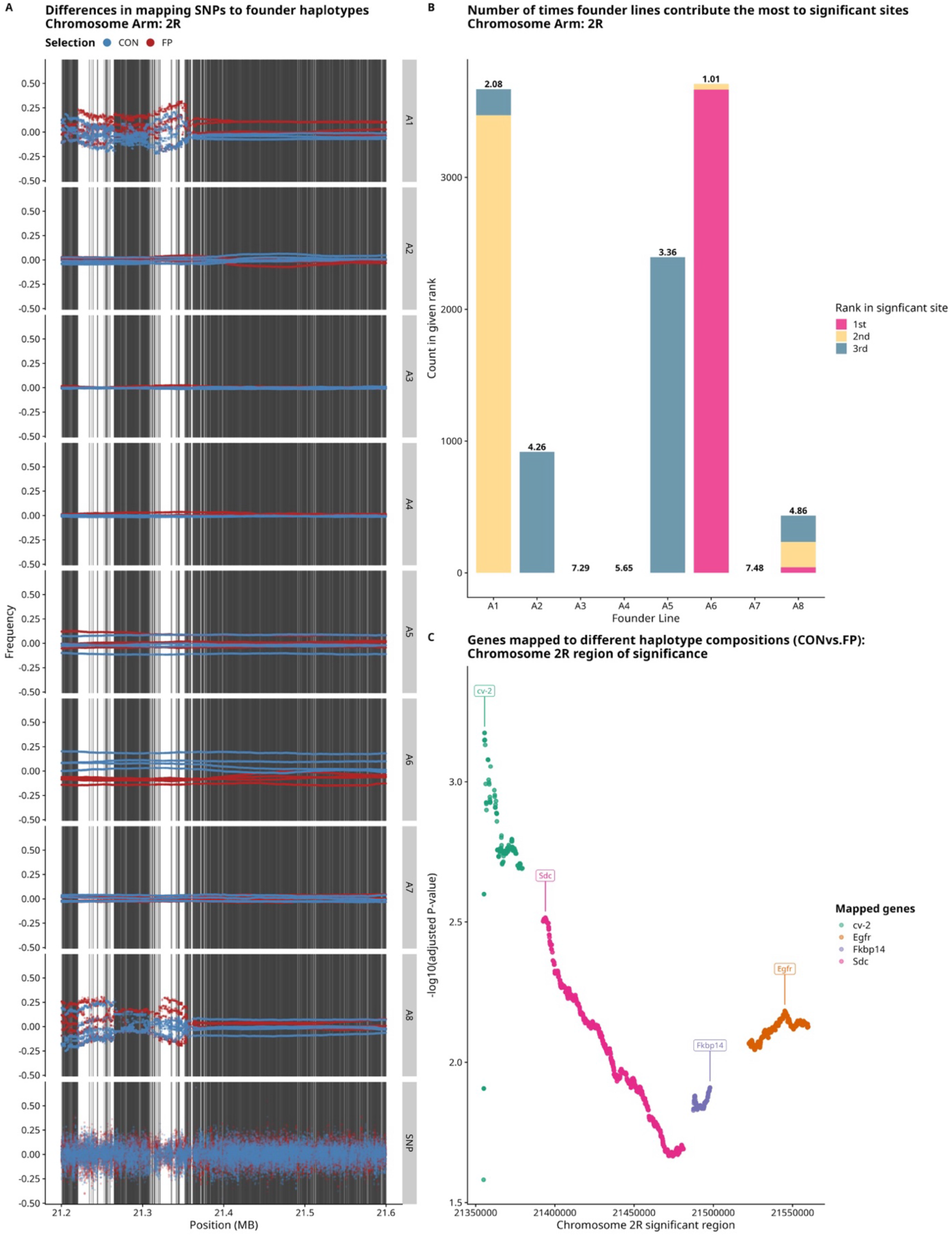
Haplotype analyses in a region of interest in chromosome arm 2R. A. Standardized haplotype frequency ascribed to each founder line, for each population, across genomic sites. The X-axis denotes genomic position. The Y-axis denotes standardized frequency for each population, colored by Selection (CON or FP). The 8 panels are labelled for each founder haplotype (A1-A8). The bottom panel marked ‘SNP’ displays the standardized reference allele frequency for the given population and genomic position. Dark grey background lines denote sites found to differ significantly between FP and CON populations. B. For each significant genomic position, we calculated the average difference between the CON and FP populations for the frequency of each founder. We ranked each founder by magnitude of difference at each site, with the greatest magnitude of differences ranked 1, and the smallest magnitude ranked 8. Thus, on each chromosomal arm, we could calculate how many times each founder was rank 1, how many times it was rank 2 and so on. The plot shows how many times a founder was rank 1, 2 or 3. Mean rank for each founder is shown on top of each respective bar. C. The genomic sites found to differ significantly in haplotype frequency between CON and FP populations. The X-axis denotes genomic position. The Y-axis shows -log10(FDR adjusted *P*-values). The mapped genes are labelled and colored accordingly. Notable genes here include *Sdc* and *Egfr* (fig. 10).

#### Gene Ontology and Enrichment

Enrichment analyses of Gene Ontology (GO) terms for Biological Processes are given in supp. fig. S13 and defined in supp. tables S25, S26. Two sets of analyses were carried out:

i. Significant SNPs of linear model analysis (fig. 7, supp. fig S13A): after enrichment analysis, the 42 SNPs altogether mapped to a total of 4 candidate genes involved in Biological Process terms. These were *pros*, *Lkb1*, CG46339 (alias *sda*) and *zfh1*. Between them, these genes, all in chromosome 3R, were significantly enriched for 4 pathways (after removal of redundant pathways using the simplify function) namely dedifferentiation, cell dedifferentiation, asymmetric neuroblast division and somatic stem cell population maintenance. The Gene Ontology and enrichment analysis table, with the respective GO IDs, pathway descriptions, adjusted P-values, q-values and gene IDs is available in supp. table S25. The Manhattan plot showing significance values for the linear model analysis (fig. 7A) also shows the enriched pathways for the significant sites, and labels the respective candidate genes.
ii. Significant SNPs from haplotype analysis: the 119,771 sites found significantly different in haplotype frequencies overlapped with 384 candidate genes involved in Biological Process terms. Enrichment of GO terms and removal of redundant pathways with the simplify function revealed 13 unique pathways across a variety of functions (supp. fig. S13B). The pathway with the most support was the negative regulation of response to stimulus (GO:0048585, adjusted P = 0.0003, q value = 0.00012). The Gene Ontology and enrichment analysis table, containing respective GO IDs, pathway descriptions, adjusted P-values, q-values and gene IDs is available in supp. table S26. The sites mapped to genes involved in enriched Biological Process terms are colored by the pathway name and labelled with the respective gene symbols on each chromosome arm (supp. fig. S14-S18, panel C).

## Discussion

To our knowledge, this is the first E&R study on direct flight performance using populations of *Drosophila melanogaster* (for the first experimental evolution study on flight performance, see (Weber 1996)). Our study approached the evolution of flight performance from a complex trait perspective, combining a wide range of phenotypic measurements with molecular evolution.

### The evolution of flight performance can be seen within 10 generations of selection

Results from the measurements of flight performance across generations indicate that there was a statistically detectable improvement in flight performance in the selected (FP) populations, while there was no evolution of flight performance in control (CON) populations (fig. 2). This is perhaps the most important and striking result of our study – that significant evolution of improved flight performance can be achieved within 10 generations of selection using the method employed in our study. The factorial test of the effects of light and wind stimuli on flight performance assay further reinforces this result, with a significant main effect of Selection, which also had a large effect size (fig. 3). This test also showed marginal significance of main effects for the stimuli provided in the flight tunnel – both light and fan had moderate effect sizes due to their presence/absence (fig. 3). While the flight tunnel method is now well proven to affect the evolution of flight performance in *D. melanogaster*, it is also a test of endurance due to the time cutoff of 15 minutes. It is possible that some flies which initially travel far into the tunnel are subsequently sent back several chambers by the increasing wind pressure (Weber 1988, 1996).

### Flight-related traits: activity levels, wing morphometrics

Results from the activity measurement assay warrant some caution in drawing inferences. This is because we observed two different patterns of results in both the assay sets (fig. 4B). While the main effect of Selection showed a large effect size and was marginally significant, the difference between activity levels in FP and CON populations was only observed in assay set 2 (fig. 4B). This might be due to high variance in the assay itself, or due to sampling at different generations. Set 2 was 2 generations after generation 10 of FP selection, whereas set 1 was 1 generation after generation 10. We also used flies sampled from set 1 for the factorial flight performance assay, which showed robust differences in flight performance between FP and CON populations (fig. 3). Although male flies consistently showed greater activity levels than female flies, this was observed in both FP and CON populations (fig. 4A). The Selection×Sex effect size was negligible.

Geometric morphometric analysis of wing size in shape in both sexes in response to selection revealed several patterns worth discussion. Sexual size dimorphism in *Drosophila* is well-established (e.g., David et al. 2003; Powell et al. 2010), and we observed the same pattern in wing centroid size: females are on average larger than males (Tables 1, 2). Selection for flight performance resulted in a significant increase in centroid size in both sexes, with a small, slightly significant interaction. These results suggest that there is an overall increase in body size resulting from 10 generations of selection for increased flight performance. Similarly, we found significant effects of selection and sex on multivariate wing shape (Table 3). After accounting for variation in shape attributable to centroid size, mean shapes were different in response to selection (P = 0.003) and between sexes (P = 0.0383). We did observe significant inter-replicate differences in wing shape as well (P = 0.0003). In contrast to size and shape, we found comparatively little evidence for directional or fluctuating asymmetry in response to selection, in contrast to studies of artificial (Garland and Freeman 2005; Castro and Garland 2018) and natural selection (Swaddle and Witter 1998).

### Life-history consequences of selection

The dry mass of adult males of FP populations showed a pattern of being greater than CON populations (fig. 6A). This potential response of selection may be due to lighter males being more susceptible to being pushed back by wind in the flight tunnel. Heavier males could potentially resist the wind pressure better at chamber levels closer to the fan. The fecundity assay displayed a visible pattern of FP populations having a greater fecundity than CON (fig. 6C). The overall per capita fecundity in the assay was very low, and we did not observe any significant linear relationship between population size and total fecundity (supp. fig. S6A, supp. table S22). This may be due to a very narrow time window for oviposition, and disturbances due to multiple plate changes. The final life-history trait assayed, egg-to-adult survivorship, showed no significant or observable patterns of differences between the selected and control populations. This may be expected, as the selection was still nascent at 10 generations. A trait as fundamental to fitness as pre-adult survivorship would likely be unaffected unless the selection pressure was strengthened many more times and applied for several more generations (e.g., (Prasad et al. 2000)).

### Associated genomic changes

The linear model test for SNP-wise comparison of allele frequencies revealed 42 sites which differed significantly between FP and CON populations. These sites were enriched for 4 Biological Process terms (supp. fig. S13A). The linear model used here was akin to a t-test with equal variance assumptions. We also carried out another, more conservative analysis by fitting a quasibinomial model (glm(family = quasibinomial(link = “logit”))), which showed no significant differences in any genomic positions between the FP and CON populations (data not shown). This is likely due to the generations of selection being too few to affect large genomic changes detectable by more conservative tests. Similarly, nucleotide diversity did not show any remarkable patterns of differences, nor was there any overlap between with the linear model results (fig. 8A). F_ST_ patterns of differences showed that the selected population FP3 primarily differed the most from the CON populations (supp. fig. S12). The outlier windows of CON vs. FP F_ST_ did not overlap with any significantly differentiated sites of the linear model (fig. 8B).

The analysis of founder haplotype frequencies revealed 119,771 significantly differentiated sites between the CON and FP populations (supp. figs. S14-S18). The 384 genes these sites mapped to were enriched for 12 pathways.

### Classifying genes of interest

Given the large number of genes found in the GO enrichment analysis, we further highlighted potentially important genes using two criteria.

1. We ranked genes by the number of significant sites present in them. The disadvantage of ranking genes thus is that large genes may contain a large number of significant sites.
2. We ranked genes by the density of significant sites present in the exon length of the gene. This accounted for smaller genes which may have a high proportion of significant sites, but may rank lower on number of significant sites due to short gene lengths.

From the genes mapped in the haplotype analysis, figure 10A shows the top 10 genes ranked by density of significant sites present in them. Figure 10B shows the top 10 genes ranked by the number of significant sites present. We have classified the genes which feature in both the top 10 lists as ‘genes of interest’. We also note those genes present in either of the top 10 lists which are involved in most enriched pathways. The genes of interest are noted in table 5.

**Figure 10:**
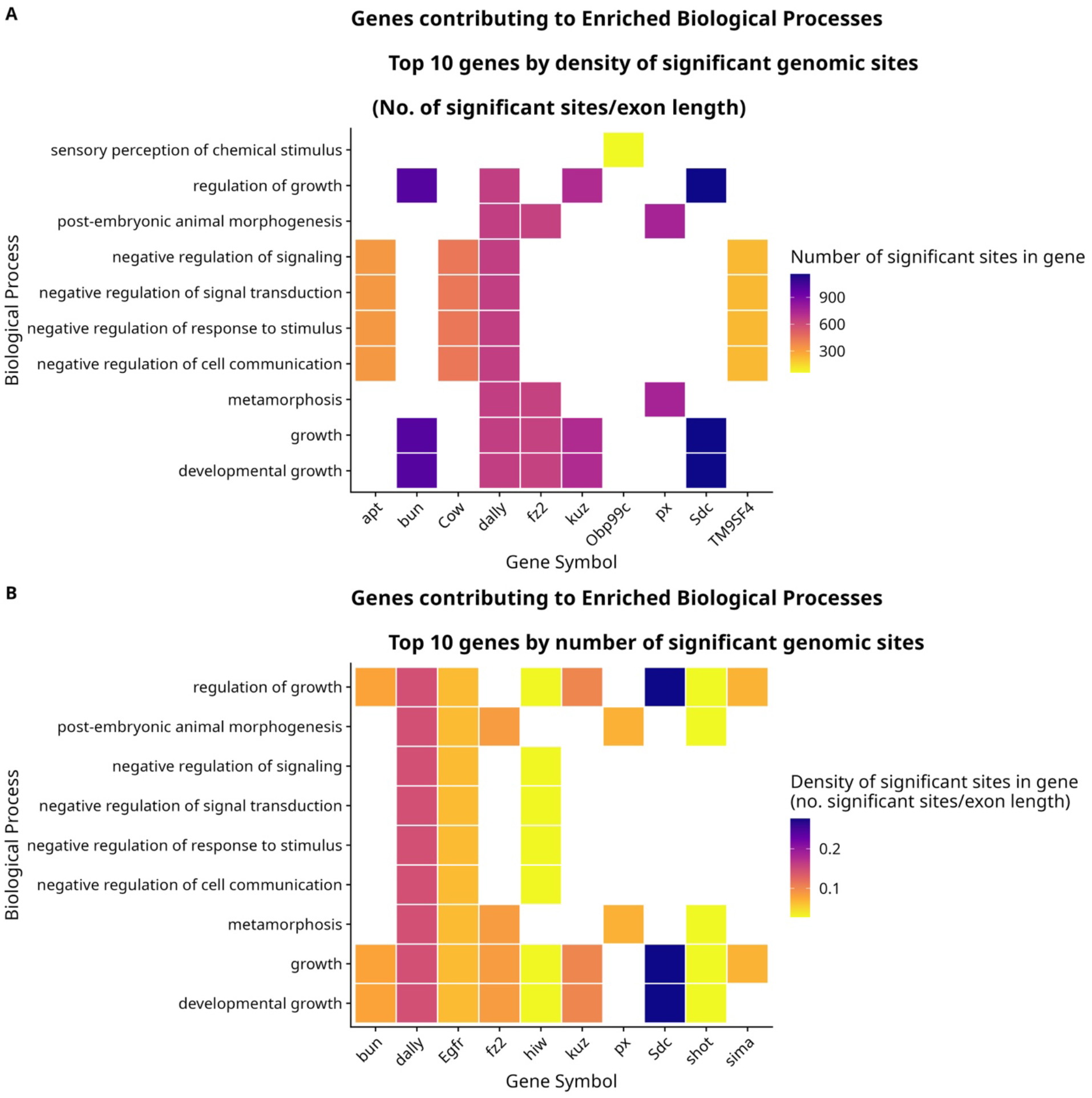
Heat plots showing top 10 genes and the pathways they are involved in. A. Top 10 genes classified by ‘density’ of significant genomic sites. Density is classified as number of significant sites divided by exon length of the gene. The X-axis shows gene symbol, and the Y-axis shows enriched pathways the gene is involved in. The heat plot is colored by number of significant genomic sites in the gene. B. Top 10 genes classified by number of significant genomic sites. The heat plot is colored by density of significant genomic sites.

**Table 5:** information on genes of interest.

| Gene symbol | Location | Notable function(s) | Flybase link |
| --- | --- | --- | --- |
| *bun* | 2L | Nervous system development | https://flybase.org/reports/FBgn0259176 |
| *dally* | 3L | Wing development, germline stem cell maintenance. | https://flybase.org/reports/FBgn0263930 |
| *fz2* | 3L | Receptor for Wnt proteins, wing margin development | https://flybase.org/reports/FBgn0016797 |
| *kuz* | 2L | Axon guidance, muscle tissue development | https://flybase.org/reports/FBgn0259984 |
| *px* | 2R | Wing vein morphogenesis | https://flybase.org/reports/FBgn0003175 |
| *Sdc* | 2R | Promotes synapse growth at larval neuromuscular junction | https://flybase.org/reports/FBgn0010415 |
| *Egfr* | 2R | Growth regulation, cell survival, developmental patterning, wing disc development | https://flybase.org/reports/FBgn0003731 |

### Gene overlaps

We performed a genome scan at both the SNP level and the haplotype level after inferring haplotype frequencies from our poolseq SNP data. The haplotype-level analysis is known to be a more powerful method in Evolve and Resequence studies (Kessner et al. 2013; Tilk et al. 2019; Linder et al. 2022; Macdonald et al. 2022), and we found many more significant sites with out haplotype-level analysis compared to the SNP-level analysis. Thus, it is not surprising that we have many sites that are significant for the haplotype analysis that do not show any significant SNPs. However, we also don’t see any overlap between our significant SNP sites and the haplotype-level analysis. One reason this occurred was for a few differentiated SNP sites, there was no haplotype information available in the founder lines and thus we did not estimate haplotype frequencies at those exact sites, though this was not the case for every significant SNP and it remains unclear why those signals are not also found in the haplotype-level analysis.

#### Finding overlaps with other molecular studies

We compared genes enriched for Biological Process terms with genes from two other notable studies, both using the Drosophila Genetic Reference Panel (DGRP) background (Mackay et al. 2012).

The first was a study of genes involved in flight performance using the gravity-based free fall tube using several DGRP lines (Spierer et al. 2021). We did not find any overlaps between the genes found in this study and the genes mapped by the significant sites from SNP differentiation analysis using the linear model (fig. 7). Between the genes mapped by the significant sites from the haplotype analysis and the compared study, we found 4 common genes between (Spierer et al. 2021) and the genes mapped by the haplotype analysis. These were *Sdc*, *Egfr*, *rhea*, *rut* (supp. table S27).

The second was a study of genes associated with wing shape (Pitchers et al. 2019). We found one overlapping gene between this study and the linear model based SNP differentiation analysis: *prospero* (*pros*), involved in neural differentiation (https://flybase.org/reports/FBgn0004595). Between the haplotype analysis and the study on wing shape (Pitchers et al. 2019), we found 18 common genes were found, and 1 common enriched pathway ‘growth’ (supp. table S28). Of these, some potentially important overlaps were *Egfr*, *Sdc*, *px* and *dally*.

### The way forward for flight performance in E&R studies

Through this study, we have shown that flight performance is very amenable to selection through the wind tunnel method. The short span of time required to evolve flight performance means that we can take two routes for future experiments:

1. Use the same experimental protocol, but with larger population sizes, stronger selection differentials, and more replicate populations.

a. Increased effective population size and replicate populations: an informative E&R study depends on having many replicate populations, as well as a large effective population size per replicate (Kofler and Schlötterer 2014). The original study (Weber 1996) used only two replicate populations, but with around 10,000 adult flies for the selection experiment. A future experiment could use similarly large numbers. Our study used four replicate populations, which could be increased to 5-6 replicates for future studies. However, larger numbers would restrict the logistics of experiments and measurements.
b. Stronger selection differentials, longer term selection: this would be a very logical next step to the successful demonstration of selection in our current study. Carrying on selection for several more generations could let us, in theory, enforce greater selection criteria (> chamber 2 in the wind tunnel, see methods).
2. Conduct several short-term selection experiments, with various changes to the original experimental protocol: this could be informed by several findings from previous studies.

a. The effect of temperature: flight performance was shown to differ based on a single generation of rearing in cold vs. warm temperatures (Frazier et al. 2008). Ambient temperatures have affected ‘free-flight performance’ in *D. melanogaster* (Lehmann 1999). Heat stress has previously changed flight behaviour (Krebs and Thompson 2006). Given the many studies on the relationship between temperature and flight performance, a future study could combine different temperature rearing and flight performance.
b. Changing the motivator in the tunnel: Light was the motivator for fly movement in our flight tunnel. Other motivators based on olfaction can be used as well. Previous studies have demonstrated a role of olfaction in guiding flight in *D. melanogaster* (Chow and Frye 2008; Bhandawat et al. 2010). In our setup, an attractive olfactory stimulus can replace light. The fan, along with providing flight resistance, could also effectively transmit the olfactory stimulus.
c. Comparing measures of flight performance: a future selection study could replace the wind tunnel method used in our study, with other methods of flight performance. Examples including a gravity-based free fall tube (Babcock and Ganetzky 2014), free-flight performance (Lehmann 1999), or a ‘flight-mill’ (Wong et al. 2018).

While we took a comprehensive approach for the study of the evolution of flight performance, ranging from stimuli in the flight tunnel (fig. 3), flight related traits such as activity (fig. 4) and wing morphology (fig. 5) and life-history related traits (fig. 6), there are many more traits that can be studied. In addition to wing morphology, both musculature (Vigoreaux 2001) and wing beat frequency (Lane et al. 2014) are relevant to flight. The latter can potentially affect male courtship songs (Ewing and Bennet-Clark 1968; Turner and Miller 2012), an important aspect of mating in flies.

## Conclusion

In our study, flight performance improved in selected populations compared to control populations. The selected populations also likely evolved activity levels, wing morphology, male dry mass, and fecundity. Whole genome analysis highlighted several enriched pathways involved in growth, development and response to stimuli, and potentially important genes for flight performance, such as *Sdc* and *dally*. Our study highlights flight performance as a robust complex trait for Evolve- and-Resequence studies using populations of *Drosophila melanogaster*.

## Acknowledgments

This study was supported by the NIH grant R35 GM149238. We thank Dr. Tara Finegan and the University of Missouri Advanced Light Microscopy Core Facility for assistance with microscopy. We thank Elliett Baca, Hailey Moore, Angie Pantoja and other members of the King lab for their help with experiments. DNA samples were sequenced by the Genomics Technology Core at the University of Missouri. The computation for this work was performed on the high-performance computing infrastructure operated by Research Support Solutions in the Division of IT at the University of Missouri, Columbia MO (https://doi.org/10.32469/10355/97710).

## Note on AI usage

No part of this manuscript was written using AI. No code was written by AI or copied from prompts. An author consulted Google Gemini 3 models for genomic data analyses and making plots, primarily for PCA, SNP differentiation, nucleotide diversity, F_ST_ and GO enrichment analyses.

## References

Adams, D. C., and E. Otárola-Castillo. 2013. geomorph: An R package for the collection and analysis of geometric morphometric shape data. Methods Ecol. Evol. 4:393–399. Wiley.

Allaire, J. J., Y. Xie, C. Dervieux, J. McPherson, J. Luraschi, K. Ushey, A. Atkins, H. Wickham, J. Cheng, W. Chang, and R. Iannone. 2026. rmarkdown: Dynamic Documents for R.

Allard, R. W., and A. D. Bradshaw. 1964. Implications of Genotype-Environmental Interactions in Applied Plant Breeding. Crop Science 4:cropsci1964.0011183X000400050021x.

Arel-Bundock, V. 2022. modelsummary: Data and Model Summaries in R. Journal of Statistical Software 103:1–23.

Babcock, D. T., and B. Ganetzky. 2014. An Improved Method for Accurate and Rapid Measurement of Flight Performance in Drosophila. JoVE 51223.

Baken, E. K., M. L. Collyer, A. Kaliontzopoulou, and D. C. Adams. 2021. geomorph v4.0 and gmShiny: enhanced analytics and a new graphical interface for a comprehensive morphometric experience. Methods Ecol. Evol., doi: 10.1111/2041-210x.13723. Wiley.

Bakka, H., H. Rue, G.-A. Fuglstad, A. I. Riebler, D. Bolin, J. Illian, E. Krainski, D. P. Simpson, and F. K. Lindgren. 2018. Spatial modelling with INLA: A review. WIRES (Invited extended review) xx:xx–xx.

Barghi, N., R. Tobler, V. Nolte, and C. Schlötterer. 2017. *Drosophila simulans* : A Species with Improved Resolution in Evolve and Resequence Studies. G3 Genes|Genomes|Genetics 7:2337–2343.

Barnes, P. T., and C. C. Laurie-Ahlberg. 1986. GENETIC VARIABILITY OF FLIGHT METABOLISM IN DROSOPHILA MELANOGASTER. III. EFFECTS OF Gpdh ALLOZYMES AND ENVIRONMENTAL TEMPERATURE ON POWER OUTPUT. Genetics 112:267–294.

Bates, D., M. Mächler, B. Bolker, and S. Walker. 2015. Fitting Linear Mixed-Effects Models Using lme4. Journal of Statistical Software 67:1–48.

Ben-Shachar, M. S., D. Lüdecke, and D. Makowski. 2020. effectsize: Estimation of Effect Size Indices and Standardized Parameters. Journal of Open Source Software 5:2815. The Open Journal.

Bhandawat, V., G. Maimon, M. H. Dickinson, and R. I. Wilson. 2010. Olfactory modulation of flight in *Drosophila* is sensitive, selective and rapid. Journal of Experimental Biology 213:4313–4313.

Bhatia, G., N. Patterson, S. Sankararaman, and A. L. Price. 2013. Estimating and interpreting FST: The impact of rare variants. Genome Research 23:1514–1521. Cold Spring Harbor Laboratory Press.

Bioconductor Core Team, and Bioconductor Package Maintainer. 2020. TxDb.Dmelanogaster.UCSC.dm6.ensGene: Annotation package for TxDb object(s).

Bonnett, O. T. 1964. Morphogenesis in the Organism and the Expression of Complex Traits. Crop Science 4:cropsci1964.0011183X000400050020x.

Brembs, B., F. Christiansen, H. J. Pflüger, and C. Duch. 2007. Flight Initiation and Maintenance Deficits in Flies with Genetically Altered Biogenic Amine Levels. J. Neurosci. 27:11122– 11131.

Burghardt, L. T., B. Epstein, J. Guhlin, M. S. Nelson, M. R. Taylor, N. D. Young, M. J. Sadowsky, and P. Tiffin. 2018. Select and resequence reveals relative fitness of bacteria in symbiotic and free-living environments. Proc. Natl. Acad. Sci. U.S.A. 115:2425–2430.

Burke, M. K., J. P. Dunham, P. Shahrestani, K. R. Thornton, M. R. Rose, and A. D. Long. 2010. Genome-wide analysis of a long-term evolution experiment with Drosophila. Nature 467:587–590.

Burke, M. K., E. G. King, P. Shahrestani, M. R. Rose, and A. D. Long. 2014a. Genome-wide association study of extreme longevity in *Drosophila melanogaster*. Genome Biology and Evolution 6:1–11.

Burke, M. K., G. Liti, and A. D. Long. 2014b. Standing Genetic Variation Drives Repeatable Experimental Evolution in Outcrossing Populations of Saccharomyces cerevisiae. Molecular Biology and Evolution 31:3228–3239.

Burke, M. K., and A. D. Long. 2012. What paths do advantageous alleles take during short-term evolutionary change? Molecular Ecology 21:4913–4916.

Burke, M. K., and M. R. Rose. 2009. Experimental evolution with *Drosophila*. American Journal of Physiology-Regulatory, Integrative and Comparative Physiology 296:R1847–R1854.

Carlson, M. 2025. org.Dm.eg.db: Genome wide annotation for Fly.

Castro, A. A., and T. Garland Jr. 2018. Evolution of hindlimb bone dimensions and muscle masses in house mice selectively bred for high voluntary wheel-running behavior. J. Morphol., doi: 10.1002/jmor.20809.

Cellini, B., and J.-M. Mongeau. 2020. Hybrid visual control in fly flight: insights into gaze shift via saccades. Current Opinion in Insect Science 42:23–31.

Chow, D. M., and M. A. Frye. 2008. Context-dependent olfactory enhancement of optomotor flight control in *Drosophila*. Journal of Experimental Biology 211:2478–2485.

Collyer, M. L., and D. C. Adams. 2018. RRPP: An R package for fitting linear models to high-dimensional data using residual randomization. Methods Ecol. Evol. 9:1772–1779. Wiley.

Conery, M., and S. F. A. Grant. 2023. Human height: a model common complex trait. Annals of Human Biology 50:258–266. Taylor & Francis.

Czech, L., Y. Peng, J. P. Spence, P. L. M. Lang, T. Bellagio, J. Hildebrandt, K. Fritschi, R. Schwab, B. A. Rowan, GrENE-net consortium, D. Weigel, J. F. Scheepens, F. Vasseur, and M. Exposito-Alonso. 2022. Monitoring rapid evolution of plant populations at scale with Pool-Sequencing. Evolutionary Biology.

Czech, L., J. P. Spence, and M. Expósito-Alonso. 2024. grenedalf: population genetic statistics for the next generation of pool sequencing. Bioinformatics 40:btae508.

Da Silva Ribeiro, T., J. A. Galván, and J. E. Pool. 2022. Maximum SNP *FST* Outperforms Full-Window Statistics for Detecting Soft Sweeps in Local Adaptation. Genome Biology and Evolution 14:evac143.

Darwin, L. J., F. A. Lemieux, R. Z. Bachtel, J. H. Blocker, C. P. Brown, A. D. Griffin, J. D. Lerman, O. C. Maule, Y. Raynes, and D. M. Rand. 2025. Genetic and environmental interactions outweigh mitonuclear coevolution for complex traits in Drosophila. bioRxiv 2025–11. Cold Spring Harbor Laboratory.

David, J. R., P. Gibert, S. Mignon-Grasteau, Hé. Legout, G. PÉtavy, C. Beaumont, and B. Moreteau. 2003. Genetic variability of sexual size dimorphism in a natural population ofDrosophila melanogaster: An isofemale-line approach. J Genet 82:79–88.

Deora, T., S. S. Sane, and S. P. Sane. 2021. Wings and halteres act as coupled dual oscillators in flies. eLife 10:e53824.

Dickson, W. B., A. D. Straw, and M. H. Dickinson. 2008. Integrative model of *Drosophila* flight. AIAA Journal 46:2150–2164.

Durinck, S., Y. Moreau, A. Kasprzyk, S. Davis, B. De Moor, A. Brazma, and W. Huber. 2005. BioMart and Bioconductor: a powerful link between biological databases and microarray data analysis. Bioinformatics 21:3439–3440.

Durinck, S., P. T. Spellman, E. Birney, and W. Huber. 2009. Mapping identifiers for the integration of genomic datasets with the R/Bioconductor package biomaRt. Nat Protoc 4:1184–1191.

Everman, E. R., S. J. Macdonald, and J. K. Kelly. 2023. The genetic basis of adaptation to copper pollution in Drosophila melanogaster. Front. Genet. 14. Frontiers.

Ewing, A. W., and H. C. Bennet-Clark. 1968. The Courtship Songs of Drosophila. Behaviour 31:288–301. Brill.

Ferretti, L., S. E. Ramos-Onsins, and M. Pérez-Enciso. 2013. Population genomics from pool sequencing. Molecular Ecology 22:5561–5576.

Fisher, R. A., F. R. Immer, and O. Tedin. 1932. The Genetical Interpretation of Statistics of the Third Degree in the Study of Quantitative Inheritance. Genetics 17:107–124.

Flaibani, N., V. E. Ortiz, J. J. Fanara, and V. P. Carreira. 2024. The relationship between morphology and flight in Drosophila: a study of two pairs of sibling species from a natural population. Insect Science 31:885–900.

Frazer, K. A., S. S. Murray, N. J. Schork, and E. J. Topol. 2009. Human genetic variation and its contribution to complex traits. Nat Rev Genet 10:241–251.

Frazier, M. R., J. F. Harrison, S. D. Kirkton, and S. P. Roberts. 2008. Cold rearing improves cold-flight performance in *Drosophila via* changes in wing morphology. Journal of Experimental Biology 211:2116–2122.

Frye, M. A., and M. H. Dickinson. 2004. Closing the loop between neurobiology and flight behavior in *Drosophila*. Current Opinion in Neurobiology 14:729–736.

Gamboa-Santarosa, K. A., G. A. Crestani, A. Moran, D. Modha, H. S. Dugo, M. Abdoli, M. K. Burke, and P. Shahrestani. 2026. Parallel adaptive responses to postponed reproduction increase lifespan and immune defense. Evolutionary Biology.

Garland, T., Jr, and P. W. Freeman. 2005. Selective breeding for high endurance running increases hindlimb symmetry. Evolution 59:1851–1854.

Garland, T., Jr., M. Zhao, and W. Saltzman. 2016. Hormones and the Evolution of Complex Traits: Insights from Artificial Selection on Behavior. Integr Comp Biol 56:207–224.

Garland, T., and M. R. Rose. 2009. Experimental Evolution: Concepts, Methods, and Applications of Selection Experiments. University of California Press.

Gidaszewski, N. A., M. Baylac, and C. P. Klingenberg. 2009. Evolution of sexual dimorphism of wing shape in the *Drosophila melanogaster* subgroup. BMC Evol. Biol. 9:110.

Glazier, A. M., J. H. Nadeau, and T. J. Aitman. 2002. Finding Genes That Underlie Complex Traits. Science 298:2345–2349. American Association for the Advancement of Science.

Gohel, D., S. Moog, and M. Heckmann. 2026. officer: Manipulation of Microsoft Word and PowerPoint Documents.

Gomes, D. G. E. 2022. Should I use fixed effects or random effects when I have fewer than five levels of a grouping factor in a mixed-effects model? PeerJ 10:e12794. PeerJ Inc.

Graves, J. L., Jr, K. L. Hertweck, M. A. Phillips, M. V. Han, L. G. Cabral, T. T. Barter, L. F. Greer, M. K. Burke, L. D. Mueller, and M. R. Rose. 2017. Genomics of Parallel Experimental Evolution in Drosophila. Molecular Biology and Evolution 34:831–842.

Graves, J. L., L. S. Luckinbill, and A. Nichols. 1988. Flight duration and wing beat frequency in long- and short-lived *Drosophila melanogaster*. Journal of Insect Physiology 34:1021– 1026.

Greenspan, Z. S., T. T. Barter, M. A. Phillips, J. M. Ranz, M. R. Rose, and L. D. Mueller. 2024. Genomewide architecture of adaptation in experimentally evolved Drosophila characterized by widespread pleiotropy. J Genet 103:8.

Hadler, N. M. 1964. GENETIC INFLUENCE ON PHOTOTAXIS IN DROSOPHILA MELANOGASTER. The Biological Bulletin 126:264–273.

Hamlin, V. L., J. Baumann, R. Heffern, A. Franke, and E. G. King. 2025. Directional selection failed to produce changes in olfactory Y-maze learning and memory but induced changes in climbing phenotypes. bioRxiv 2025.05.18.654242.

Harrisson, K. A., A. Pavlova, M. Telonis-Scott, and P. Sunnucks. 2014. Using genomics to characterize evolutionary potential for conservation of wild populations. Evolutionary Applications 7:1008–1025.

Hedrick, T. L., S. A. Combes, and L. A. Miller. 2015. Recent developments in the study of insect flight. Can. J. Zool. 93:925–943.

Hesselberg, T., and F.-O. Lehmann. 2009. The role of experience in flight behaviour of *Drosophila*. Journal of Experimental Biology 212:3377–3386.

Hillis, D. A., and T. Garland. 2023. Multiple solutions at the genomic level in response to selective breeding for high locomotor activity. GENETICS 223:iyac165.

Hirsch, J., and J. C. Boudreau. 1958. Studies in experimental behavior genetics: I. The heritability of phototaxis in a population of Drosophila Melanogaster. Journal of Comparative and Physiological Psychology 51:647–651. American Psychological Association, US.

Hoedjes, K. M., J. van den Heuvel, M. Kapun, L. Keller, T. Flatt, and B. J. Zwaan. 2019. Distinct genomic signals of lifespan and life history evolution in response to postponed reproduction and larval diet in Drosophila. Evol Lett 3:598–609.

Houle, D., J. Mezey, P. Galpern, and A. Carter. 2003. Automated measurement of *Drosophila* wings. BMC Evol Biol 3:25.

Hudson, R. R., M. Slatkin, and W. P. Maddison. 1992. Estimation of levels of gene flow from DNA sequence data. Genetics 132:583–589.

Hvitfeldt, E. 2021. paletteer: Comprehensive Collection of Color Palettes.

Iannone, R., J. Cheng, B. Schloerke, S. Haughton, E. Hughes, A. Lauer, R. François, J. Seo, K. Brevoort, and O. Roy. 2026. gt: Easily Create Presentation-Ready Display Tables.

Kawecki, T. J., R. E. Lenski, D. Ebert, B. Hollis, I. Olivieri, and M. C. Whitlock. 2012. Experimental evolution. Trends in Ecology & Evolution 27:547–560.

Kessner, D., T. L. Turner, and J. Novembre. 2013. Maximum Likelihood Estimation of Frequencies of Known Haplotypes from Pooled Sequence Data. Mol Biol Evol 30:1145–1158.

King, E. G., S. J. Macdonald, and A. D. Long. 2012a. Properties and power of the *Drosophila* Synthetic Population Resource for the routine dissection of complex traits. Genetics 191:935–949.

King, E. G., C. M. Merkes, C. L. McNeil, S. R. Hoofer, S. Sen, K. W. Broman, A. D. Long, and S. J. Macdonald. 2012b. Genetic dissection of a model complex trait using the *Drosophila* Synthetic Population Resource. Genome Res. 22:1558–1566.

King, E. G., C. M. Merkes, C. L. McNeil, S. R. Hoofer, S. Sen, K. W. Broman, A. D. Long, and S. J. Macdonald. 2012c. Genetic dissection of a model complex trait using the *Drosophila* Synthetic Population Resource. Genome Res 22:1558–1566.

Klingenberg, C. P., M. Barluenga, and A. Meyer. 2002. Shape analysis of symmetric structures: quantifying variation among individuals and asymmetry. Evolution 56:1909–1920.

Klingenberg, C. P., and G. S. McIntyre. 1998. Geometric morphometrics of developmental instability: Analyzing patterns of fluctuating asymmetry with Procrustes methods. Evolution 52:1363–1375.

Kofler, R., and C. Schlötterer. 2014. A Guide for the Design of Evolve and Resequencing Studies. Molecular Biology and Evolution 31:474–483.

Krassowski, M. 2020. ComplexUpset.

Krebs, R. A., and K. A. Thompson. 2006. Direct and correlated effects of selection on flight after exposure to thermal stress in Drosophila melanogaster. Genetica 128:217–225.

Krishna, S., M. Cho, H.-N. Wehmann, T. Engels, and F.-O. Lehmann. 2020. Wing design in flies: properties and aerodynamic function. Insects 11:466. Multidisciplinary Digital Publishing Institute.

Kuznetsova, A., P. B. Brockhoff, and R. H. B. Christensen. 2017. lmerTest package: Tests in linear mixed effects models. J. Stat. Softw. 82:1–26. Foundation for Open Access Statistic.

Lander, E. S., and N. J. Schork. 1994. Genetic Dissection of Complex Traits. 265.

Lane, S. J., W. A. Frankino, M. M. Elekonich, and S. P. Roberts. 2014. The effects of age and lifetime flight behavior on flight capacity in *Drosophila melanogaster*. Journal of Experimental Biology 217:1437–1443.

Laughlin, H. H. 1933. The General Formula of Heredity. Proc. Natl. Acad. Sci. U.S.A. 19:787–801.

Lawrence, M., W. Huber, H. Pagès, P. Aboyoun, M. Carlson, R. Gentleman, M. Morgan, and V. Carey. 2013. Software for Computing and Annotating Genomic Ranges. PLoS Computational Biology 9.

Lehmann, F.-O. 1999. Ambient temperature affects free-flight performance in the fruit fly Drosophila melanogaster. Journal of Comparative Physiology B: Biochemical, Systemic, and Environmental Physiology 169:165–171.

Lehmann, F.-O., and M. H. Dickinson. 1997. The Changes in Power Requirements and Muscle Efficiency During Elevated Force Production in the Fruit Fly *Drosophila Melanogaster*. Journal of Experimental Biology 200:1133–1143.

Lenth, R. V., and J. Piaskowski. 2026. emmeans: Estimated Marginal Means, aka Least-Squares Means.

Lex, A., N. Gehlenborg, H. Strobelt, R. Vuillemot, and H. Pfister. 2014. UpSet: Visualization of Intersecting Sets,. IEEE Transactions on Visualization and Computer Graphics 20:1983– 1992.

Linder, R. A., B. Zabanavar, A. Majumder, H. C.-S. Hoang, V. G. Delgado, R. Tran, V. T. La, S. W. Leemans, and A. D. Long. 2022. Adaptation in Outbred Sexual Yeast is Repeatable, Polygenic and Favors Rare Haplotypes. Mol Biol Evol 39:msac248.

Long, A. D., S. J. Macdonald, and E. G. King. 2014. Dissecting complex traits using the *Drosophila* Synthetic Population Resource. Trends in Genetics 30:488–495.

Long, A., G. Liti, A. Luptak, and O. Tenaillon. 2015. Elucidating the molecular architecture of adaptation via evolve and resequence experiments. Nat Rev Genet 16:567–582.

Loshchilov, I., and F. Hutter. 2019. Decoupled Weight Decay Regularization. arXiv.

Macdonald, S. J., K. M. Cloud-Richardson, D. J. Sims-West, and A. D. Long. 2022. Powerful, efficient QTL mapping in Drosophila melanogaster using bulked phenotyping and pooled sequencing. Genetics 220:iyab238.

Mackay, T. F. C. 2009. The genetic architecture of complex behaviors: lessons from Drosophila. Genetica 136:295–302.

Mackay, T. F. C., and R. R. H. Anholt. 2006. Of Flies and Man: *Drosophila* as a Model for Human Complex Traits. Annu. Rev. Genom. Hum. Genet. 7:339–367.

Mackay, T. F. C., S. Richards, E. A. Stone, A. Barbadilla, J. F. Ayroles, D. Zhu, S. Casillas, Y. Han, M. M. Magwire, J. M. Cridland, M. F. Richardson, R. R. H. Anholt, M. Barrón, C. Bess, K. P. Blankenburg, M. A. Carbone, D. Castellano, L. Chaboub, L. Duncan, Z. Harris, M. Javaid, J. C. Jayaseelan, S. N. Jhangiani, K. W. Jordan, F. Lara, F. Lawrence, S. L. Lee, P. Librado, R. S. Linheiro, R. F. Lyman, A. J. Mackey, M. Munidasa, D. M. Muzny, L. Nazareth, I. Newsham, L. Perales, L.-L. Pu, C. Qu, M. Ràmia, J. G. Reid, S. M. Rollmann, J. Rozas, N. Saada, L. Turlapati, K. C. Worley, Y.-Q. Wu, A. Yamamoto, Y. Zhu, C. M. Bergman, K. R. Thornton, D. Mittelman, and R. A. Gibbs. 2012. The Drosophila melanogaster Genetic Reference Panel. Nature 482:173–178.

Marden, J. H. 2000. Variability in the Size, Composition, and Function of Insect Flight Muscles. Annu. Rev. Physiol. 62:157–178.

Marden, J. H., M. R. Wolf, and K. E. Weber. 1997. Aerial performance of *Drosophila melanogaster* from populations selected for upwind flight ability. Journal of Experimental Biology 200:2747–2755.

Mardia, K. 2000. Statistical assessment of bilateral symmetry of shapes. Biometrika 87:285–300. Oxford University Press (OUP).

Mathis, A., T. Biasi, S. Schneider, M. Yüksekgönül, B. Rogers, M. Bethge, and M. W. Mathis. 2019. Pretraining boosts out-of-domain robustness for pose estimation.

Mathis, A., P. Mamidanna, K. M. Cury, T. Abe, V. N. Murthy, M. W. Mathis, and M. Bethge. 2018. DeepLabCut: markerless pose estimation of user-defined body parts with deep learning. Nat. Neurosci. 21:1281–1289.

Mazerolle, M. J. 2023. AICcmodavg: Model selection and multimodel inference based on (Q)AIC(c).

Nath, T., A. Mathis, A. C. Chen, A. Patel, M. Bethge, and M. W. Mathis. 2019. Using DeepLabCut for 3D markerless pose estimation across species and behaviors. Nat. Protoc. 14:2152– 2176.

Nei, M. 1986. Definition and Estimation of Fixation Indices. Evolution 40:643–645. Oxford University Press.

Neuwirth, E. 2022. RColorBrewer: ColorBrewer Palettes.

Obenchain, V., M. Lawrence, V. Carey, S. Gogarten, P. Shannon, and M. Morgan. 2014. VariantAnnotation: a Bioconductor package for exploration and annotation of genetic variants. Bioinformatics 30:2076–2078.

Pagès, H., M. Carlson, S. Falcon, and N. Li. 2026. AnnotationDbi: Manipulation of SQLite-based annotations in Bioconductor.

Paszke, A., S. Gross, F. Massa, A. Lerer, J. Bradbury, G. Chanan, T. Killeen, Z. Lin, N. Gimelshein, L. Antiga, A. Desmaison, A. Köpf, E. Yang, Z. DeVito, M. Raison, A. Tejani, S. Chilamkurthy, B. Steiner, L. Fang, J. Bai, and S. Chintala. 2019. PyTorch: An Imperative Style, High-Performance Deep Learning Library. arXiv.

Pedersen, T. L. 2025. patchwork: The Composer of Plots.

Petukhov, V., T. van den Brand, and E. Biederstedt. 2023. ggrastr: Rasterize Layers for ‘ggplot2’.

Phillips, M. A., I. C. Kutch, A. D. Long, and M. K. Burke. 2020. Increased time sampling in an evolve-and-resequence experiment with outcrossing Saccharomyces cerevisiae reveals multiple paths of adaptive change. Molecular Ecology 29:4898–4912.

Pitchers, W., J. Nye, E. J. Márquez, A. Kowalski, I. Dworkin, and D. Houle. 2019. A multivariate genome-wide association study of wing shape in *Drosophila melanogaster*. Genetics 211:1429–1447.

Plomin, R., M. J. Owen, and P. McGuffin. 1994. The Genetic Basis of Complex Human Behaviors. Science 264:1733–1739. American Association for the Advancement of Science.

Powell, A. M., M. Davis, and J. R. Powell. 2010. Phenotypic plasticity across 50MY of evolution: *Drosophila* wing size and temperature. Journal of Insect Physiology 56:380–382.

Prasad, N. G., M. Shakarad, V. M. Gohil, V. Sheeba, M. Rajamani, and A. Joshi. 2000. Evolution of reduced pre-adult viability and larval growth rate in laboratory populations of *Drosophila melanogaster* selected for shorter development time. Genet. Res. 76:249–259.

R Core Team. 2026. R: A Language and Environment for Statistical Computing. R Foundation for Statistical Computing, Vienna, Austria.

Rêgo, A., J. Baur, C. Girard-Tercieux, M. De La Paz Celorio-Mancera, R. Stelkens, and D. Berger. 2025. Repeatability of evolution and genomic predictions of temperature adaptation in seed beetles. Nat Ecol Evol 9:1061–1074.

Robertson, F. W., and J. H. Sang. 1944. The Ecological Determinants of Population Growth in a Drosophila Culture. I. Fecundity of Adult Flies. Proceedings of the Royal Society of London. Series B, Biological Sciences 132:258–277. Royal Society.

Rodriguez-Sanchez, F., and C. P. Jackson. 2025. grateful: Facilitate citation of R packages.

Rue, H., S. Martino, and N. Chopin. 2009. Approximate Bayesian Inference for Latent Gaussian Models Using Integrated Nested Laplace Approximations (with discussion). Journal of the Royal Statistical Society B 71:319–392.

Sadaf, S., O. V. Reddy, S. P. Sane, and G. Hasan. 2015. Neural Control of Wing Coordination in Flies. Current Biology 25:80–86.

Sane, S. P. 2003. The aerodynamics of insect flight. J Exp Biol 206:4191–4208.

Santos, M. A., A. Carromeu-Santos, A. S. Quina, M. A. Antunes, T. N. Kristensen, M. Santos, M. Matos, I. Fragata, and P. Simões. 2024. Experimental Evolution in a Warming World: The *Omics* Era. Molecular Biology and Evolution 41:msae148.

Schlötterer, C., R. Tobler, R. Kofler, and V. Nolte. 2014. Sequencing pools of individuals — mining genome-wide polymorphism data without big funding. Nat Rev Genet 15:749–763.

Sella, G., and N. H. Barton. 2019. Thinking About the Evolution of Complex Traits in the Era of Genome-Wide Association Studies. Annu. Rev. Genom. Hum. Genet. 20:461–493.

Shahrestani, P., E. King, R. Ramezan, M. Phillips, M. Riddle, M. Thornburg, Z. Greenspan, Y. Estrella, K. Garcia, P. Chowdhury, G. Malarat, M. Zhu, S. M. Rottshaefer, S. Wraight, M. Griggs, J. Vandenberg, A. D. Long, A. G. Clark, and B. P. Lazzaro. 2021. The molecular architecture of *Drosophila melanogaster* defense against *Beauveria bassiana* explored through evolve and resequence and quantitative trait locus mapping. G3 Genes|Genomes|Genetics 11:jkab324.

Slowikowski, K. 2026. ggrepel: Automatically Position Non-Overlapping Text Labels with “ggplot2.”, doi: 10.32614/CRAN.package.ggrepel.

Spierer, A. N., J. A. Mossman, S. P. Smith, L. Crawford, S. Ramachandran, and D. M. Rand. 2021. Natural variation in the regulation of neurodevelopmental genes modifies flight performance in Drosophila. PLOS Genetics 17:e1008887. Public Library of Science.

Sun, K., B. Xiao, D. Liu, and J. Wang. 2019. Deep High-Resolution Representation Learning for Human Pose Estimation. Pp. 5686–5696 in 2019 IEEE/CVF Conference on Computer Vision and Pattern Recognition (CVPR).

Sun, M. 2014. Insect flight dynamics: Stability and control. Rev. Mod. Phys. 86:615–646.

Swaddle, J. P., and M. S. Witter. 1998. Cluttered habitats reduce wing asymmetry and increase flight performance in European starlings. Behavioral Ecology and Sociobiology 42:281– 287.

Tammero, L. F., and M. H. Dickinson. 2002. Collision-avoidance and landing responses are mediated by separate pathways in the fruit fly, *Drosophila melanogaster*. Journal of Experimental Biology 205:2785–2798.

Tautz, D., L. F. Pallares, L. Andersson, N. Barghi, N. Barton, R. Bay, Y. F. Chan, A. Hancock, T. S. Kaiser, D. Koenig, Z. Kontarakis, M. Liedvogel, J. de Meaux, M. Nordborg, A. A. Palmer, M. Purugganan, C. Schlötterer, K. Schmid, D. Y. R. Stainier, D. Weigel, J. B. W. Wolf, D. Ebert, and G. Gibson. 2026. Beyond Mendel: a call to revisit the genotype-phenotype map through new experimental paradigms. Genetics 232:iyag024. Oxford University Press (OUP).

Taylor, G. K. 2001. Mechanics and aerodynamics of insect flight control. Biological Reviews 76:449–471.

Tenaillon, O., J. E. Barrick, N. Ribeck, D. E. Deatherage, J. L. Blanchard, A. Dasgupta, G. C. Wu, S. Wielgoss, S. Cruveiller, C. Médigue, D. Schneider, and R. E. Lenski. 2016. Tempo and mode of genome evolution in a 50,000-generation experiment. Nature 536:165–170.

Tilk, S., A. Bergland, A. Goodman, P. Schmidt, D. Petrov, and S. Greenblum. 2019. Accurate Allele Frequencies from Ultra-low Coverage Pool-Seq Samples in Evolve-and-Resequence Experiments. G3 Genes|Genomes|Genetics 9:4159–4168.

Tobler, R., S. U. Franssen, R. Kofler, P. Orozco-terWengel, V. Nolte, J. Hermisson, and C. Schlötterer. 2014. Massive Habitat-Specific Genomic Response in D. melanogaster Populations during Experimental Evolution in Hot and Cold Environments. Molecular Biology and Evolution 31:364–375.

Turner, T. L., and P. M. Miller. 2012. Investigating Natural Variation in *Drosophila* Courtship Song by the Evolve and Resequence Approach. Genetics 191:633–642.

Vigoreaux, J. O. 2001. Genetics of the Drosophila flight muscle myofibril: a window into the biology of complex systems. BioEssays 23:1047–1063.

Vlachos, C., and R. Kofler. 2019. Optimizing the Power to Identify the Genetic Basis of Complex Traits with Evolve and Resequence Studies. Molecular Biology and Evolution 36:2890– 2905.

Vogel, S. 1967a. Flight in *Drosophila*. Journal of Experimental Biology 46:383–392.

Vogel, S. 1967b. Flight in Drosophila: II. Variations in Stroke Parameters and Wing Contour. J Exp Biol 46:383–392.

Wang, J., K. Sun, T. Cheng, B. Jiang, C. Deng, Y. Zhao, D. Liu, Y. Mu, M. Tan, X. Wang, W. Liu, and B. Xiao. 2020. Deep High-Resolution Representation Learning for Visual Recognition. arXiv.

Ward, M. J., S. A. Ellsworth, E. G. King, E. Ng’oma, G. S. Nystrom, K. C. Lawrence, L. Maquet-Diafouka, A. Oliver, M. J. Margres, C. L. Parkinson, K. A. Hughes, and D. R. Rokyta. 2026. The genetic architecture of venom resistance: a novel approach to target identification and coevolutionary discovery. Mol Biol Evol 43:msag048.

Weber, K. E. 1988. An apparatus for selection on flying speed. Drosophila Information Services 67:93–95.

Weber, K. E. 1996. Large genetic change at small fitness cost in large populations of *Drosophila melanogaster* selected for wind tunnel flight: rethinking fitness surfaces. Genetics 144:205–213.

Wickham, H., M. Averick, J. Bryan, W. Chang, L. McGowan, R. François, G. Grolemund, A. Hayes, L. Henry, J. Hester, M. Kuhn, T. Pedersen, E. Miller, S. Bache, K. Müller, J. Ooms, D. Robinson, D. Seidel, V. Spinu, K. Takahashi, D. Vaughan, C. Wilke, K. Woo, and H. Yutani. 2019. Welcome to the tidyverse. J. Open Source Softw. 4:1686. The Open Journal.

Wickham, H., and J. Bryan. 2026. readxl: Read Excel Files.

Wightman, R., N. Raw, A. Soare, A. Arora, C. Ha, C. Reich, F. Guan, J. Kaczmarzyk, mrT23, Mike, SeeFun, contrastive, M. Rizin, H. Kim, C. Kertész, D. Mehta, G. Cucurull, K. Singh, hankyul, Y. Tatsunami, A. Lavin, J. Zhuang, M. Hollemans, M. Rashad, S. Sameni, V. Shults, Lucain, X. Wang, Y. Kwon, and Y. Uchida. 2023. rwightman/pytorch-image-models: v0.8.10dev0 Release., doi: 10.5281/zenodo.7618837. Zenodo.

Wilke, C. O. 2025a. cowplot: Streamlined Plot Theme and Plot Annotations for “ggplot2.”

Wilke, C. O. 2025b. ggridges: Ridgeline Plots in “ggplot2.”, doi: 10.32614/CRAN.package.ggridges.

Williams-Simon, P. A., C. Oster, J. A. Moaton, R. Ghidey, E. Ng’oma, K. M. Middleton, and E. G. King. 2024. Naturally segregating genetic variants contribute to thermal tolerance in a *Drosophila melanogaster* model system. Genetics 227:iyae040.

Wong, J. S., A. C. Cave, D. M. Lightle, W. F. Mahaffee, S. E. Naranjo, N. G. Wiman, J. M. Woltz, and J. C. Lee. 2018. Drosophila suzukii flight performance reduced by starvation but not affected by humidity. J Pest Sci 91:1269–1278.

Wright, K. 2025. pals: Color Palettes, Colormaps, and Tools to Evaluate Them.

Wu, X., T. Bellagio, Y. Peng, L. Czech, M. Lin, P. Lang, R. Epstein, M. Abdelaziz, J. Alexander, C. Alonso-Blanco, H. L. Andersen, M. Berbel, J. Bergelson, O. Bossdorf, L. Burghardt, M. Caton-Darby, R. Colautti, C. Delker, P. G. Dimitrakopoulos, K. Donohue, W. Durka, G. Escribano-Avila, S. J. Franks, F. B. Fritschi, A. Galanidis, A. Garcia-Fernández, A. García-Muñoz, E. Hamann, A. Hutt, J. M. Iriondo, T. E. Juenger, S. R. Keller, K. Koehl, A. Korte, P. Korte, A. Kutschera, C. Lara-Romero, L. Leventhal, D. Maag, A. Marcer, M. March-Salas, J. De Meaux, B. Méndez-Vigo, J. Morente-López, T. C. Morton, Z. Münzbergova, A. Muola, H. A. Nomoto, M. Pärtel, F. X. Picó, B. Quarles-Chidyagwai, M. Quint, N. Reichelt, A. Rudak, J. Schmitt, G. Schmitz, M. Seifan, B. L. Snoek, R. Stam, M. Stift, J. R. Stinchcombe, M. A. Taylor, P. Tiffin, I. Till-Bottraud, A. Traveset, J.-G. Valay, M. Van Zanten, V. Vandvik, C. Violle, D. Weigel, M. Wódkiewicz, F. Vasseur, J. F. Scheepens, and M. Exposito-Alonso. 2026. Rapid adaptation and extinction in synchronized outdoor evolution experiments of *Arabidopsis*. Science 391:eadz0777.

Xiao, N. 2026. ggsci: Scientific Journal and Sci-Fi Themed Color Palettes for “ggplot2.”, doi: 10.32614/CRAN.package.ggsci.

Xie, Y., J. J. Allaire, and G. Grolemund. 2018. R Markdown: The Definitive Guide. Chapman and Hall/CRC, Boca Raton, Florida.

Yengo, L., J. Sidorenko, K. E. Kemper, Z. Zheng, A. R. Wood, M. N. Weedon, T. M. Frayling, J. Hirschhorn, J. Yang, P. M. Visscher, and the GIANT Consortium. 2018. Meta-analysis of genome-wide association studies for height and body mass index in ∼700000 individuals of European ancestry. Hum Mol Genet 27:3641–3649.

Yu, G. 2026. enrichplot: Visualization of Functional Enrichment Result.

Yu, G. 2024. Thirteen years of clusterProfiler. The Innovation 5:100722.

Yu, G., L.-G. Wang, Y. Han, and Q.-Y. He. 2012. clusterProfiler: an R package for comparing biological themes among gene clusters. OMICS: A Journal of Integrative Biology 16:284– 287.

