## Supplemental methods, figures, tables for "Complex-trait dissection with Evolve-and-Resequence: Selection for improved flight performance in laboratory fruit fly populations"

*Results for Haplotype Inference Simulations*

Several previous methods have recognized that it is possible to estimate the most likely set of haplotype frequencies that would give rise to the observed set of allele frequencies for a given genomic region. These methods all perform well and they consistently show that it is possible to obtain accurate haplotype frequency estimates even for modest coverage because of the method's ability to leverage SNP information across multiple sites (Kessner et al. 2013; Tilk et al. 2019; Linder et al. 2022; Macdonald et al. 2022) For any of these estimation methods, the experimenter must decide on the genomic window size for the estimate as well as the sequencing effort to employ, affecting coverage. Here, we compare two methods for estimating haplotype frequencies from pooled sequencing data which rely only on a SNP table data rather than the raw sequencing reads. We explore different window sizes and different coverage to determine how each affects the accuracy of the haplotype estimation and to provide justification for the experimental choices we made for this study.

(i) Estimation methods

We developed and compared two different methods for estimating the set of haplotype frequencies from pooled allele frequencies. We refer to these methods as "optimality" and "likelihood" throughout. Both methods rely on two sources of data: 1) the allele counts from pooled sequencing at the set of variant positions and 2) the allele assignments for the set of haplotypes that contributed to the pool. For our application, these consist of 1) a set of simulated poolseq allele counts from an *in silico* sampled DSPR RIL pool and 2) the allele calls for the set of DSPR founder lines.

Optimality method:

At each focal position, we used the optim function in R to identify the set of founder haplotype frequencies that minimize the following quantity (similar to a sum of squares):

$$\sum_{i=1}^{nsnp} \sqrt{c_i} \cdot \left( MAF_i - \sum_{j=1}^{8} \left( f_{i,j} * h_{i,j} \right) \right)^2 + \left( 100 \cdot \left( \left( \sum_{j=1}^{8} h_{i,j} \right) - 1 \right)^2 \right)$$

where $c_i$ is the sequence coverage at the *i*th SNP in the window consisting of *nsnp* total SNPs, $MAF_i$ is the alternative allele frequency at the *i*th SNP, $f_{i,j}$ is the allelic state of the *j*th founder at the *i*th SNP (0/1), and $h_{i,j}$ is the haplotype frequency of the *j*th founder at the *i*th SNP. The second part of the above equation serves as a penalty when the sum of the 8 founder haplotype frequencies exceeds 1 to prevent this occurrence. In addition, each individual haplotype frequency is bounded by 0 and 0.95.

We have previously used this method in an xQTL study of extreme lifespan (Burke et al. 2014) and in an experimental evolution application on venom resistance (Ward et al. 2026).

Likelihood method:

The likelihood method is similar to the optimality method in that it relies on the optimization of a function, but it explicitly calculates the likelihood of observing a given set of allele counts given a set of haplotype frequencies. Here, we again used the optim function in R to identify the set of founder haplotype frequencies that minimize the following quantity:

$$-\sum_{i=1}^{nsnp} \ln \left( \binom{c_i}{a_i} p_i^{a_i} (1 - p_i)^{c_i - a_i} \right)$$

where

$$p_i = \sum_{j=1}^{8} f_{i,j} * h_{i,j}$$

This method uses calculates the likelihood of observing $a_i$ reads (i.e. count of reference allele reads) out of a total of $c_i$ reads (i.e. coverage) at the *ith* SNP using the dbinom function in R where the probability is the expected pooled SNP frequency ($p_i$) calculated based on the set of founder haplotype frequencies ($h_{i,j}$ is the frequency of the *jth* founder haplotype at the *ith* SNP) and the allele calls for each founder haplotype ($f_{i,j}$ is the allele call of the *jth* founder haplotype at the *ith* SNP) for a given window containing *nsnp* number of SNPs. The calculated pooled allele frequencies are constrained to be between 0 and 1, and each founder haplotype frequency is also bounded to be between 0 and 1.

(ii) Simulations & Parameters

We used the DSPR to simulate pooled allele frequencies. The DSPR is a multiparent population consisting of an 8-way, 50 generation intercross, followed by 25 generations of inbreeding to produce stable recombinant inbred lines (RIL). Each RIL has a mosaic genome consisting of segments of the original 8 founder lines. Previous full genome sequencing of the founder lines and dense genotyping of the RILs has produced a resource in which the haplotype identity of each genomic segment in each RIL is known with a high degree of confidence (King et al. 2012a,b). Thus, we can create a pool of RILs *in silico* and calculate a set of "pooled" genome-wide allele frequencies. Then, we can simulate different sequencing coverage using these frequencies and sampling the binomial distribution. Once we have generated these *in silico* pool-seq datasets, we can analyze these datasets using different window sizes to estimate haplotype frequencies at a given position using each of our methods described above.

We simulated poolseq datasets by sampling 800 DSPR RILs with replacement. We then randomly selected 500 positions on each of the five major chromosome arms (*X, 2L, 2R, 3L, 3R*) to perform haplotype inference. We then created poolseq data for a window centered on each position by first calculating the pooled allele frequency from the known SNP calls for the set of sampled 800 RILs. We simulated window sizes for the following sizes: 500bp, 1kb, 2kb, 5kb, 10kb, 20kb, 50kb, 100kb, 200kb, 500kb. Each of these occurred at the same position for the same sampled set of RILs, allowing direct comparison. Then, we used these frequencies to simulate coverage for the following different coverage values: 1, 2, 5, 10, 25, 50, 75, 100, 200 by sampling from the binomial distribution to produce two allele frequency counts at each position. We also considered the case where we did not simulate coverage but used the calculated frequencies directly (i.e. the ideal case of perfect frequency estimation). We then used this pooled frequency data to estimate the haplotype frequencies using each of our estimation methods described above.

Given we simulated these pooled frequencies from a known set of RILs, we have ground truth information for the true set of frequencies and can compare the estimated haplotype frequencies to the true set to estimate the error rates for each method for each parameter combination. Here, we used the mean absolute difference between the true haplotype frequencies and the estimated haplotype frequencies as our metric of error.

(iii) Performance of estimation methods for different coverage and window sizes

Once coverage is greater than ~25, the improvement in the error rate in both estimation methods is less dramatic (Figure S1a). This effect is due to the leveraging of many positions into the estimate, leading to the method being tolerant of error at individual sites. Thus, the benefit of increased coverage beyond ~25 is greatest when the window size is also small (i.e.  fewer SNPs to leverage).

The window sizes with the lowest error are 100 and 200kb and both have very similar error rates(Figure S1a). The window size with the lowest error also depends on coverage, with 200kb having a slightly lower error rate when coverage is modest but 100kb having a lower error rate once coverage is greater than 75. Smaller window sizes have higher error rates due to fewer SNPs contributing to the estimation while larger window sizes begin to encompass recombination breakpoints which add error to the estimation of a given haplotype frequency.

Both methods perform well and similarly; however, the likelihood method does slightly outperform the optimality method across the parameter space (Figure S1b,c). This comparison is easiest to see for parameter spaces restricted to show a smaller range (Figure S1b,c)

(iv) We based our haplotype estimation methods for this study on these simulation results. We chose to use the likelihood method due to its more formal estimation of likelihood and higher performance in our simulations. We targeted a sequencing coverage of 100 and used a window size of 100kb to ensure accurate haplotype frequency estimation from our dataset.

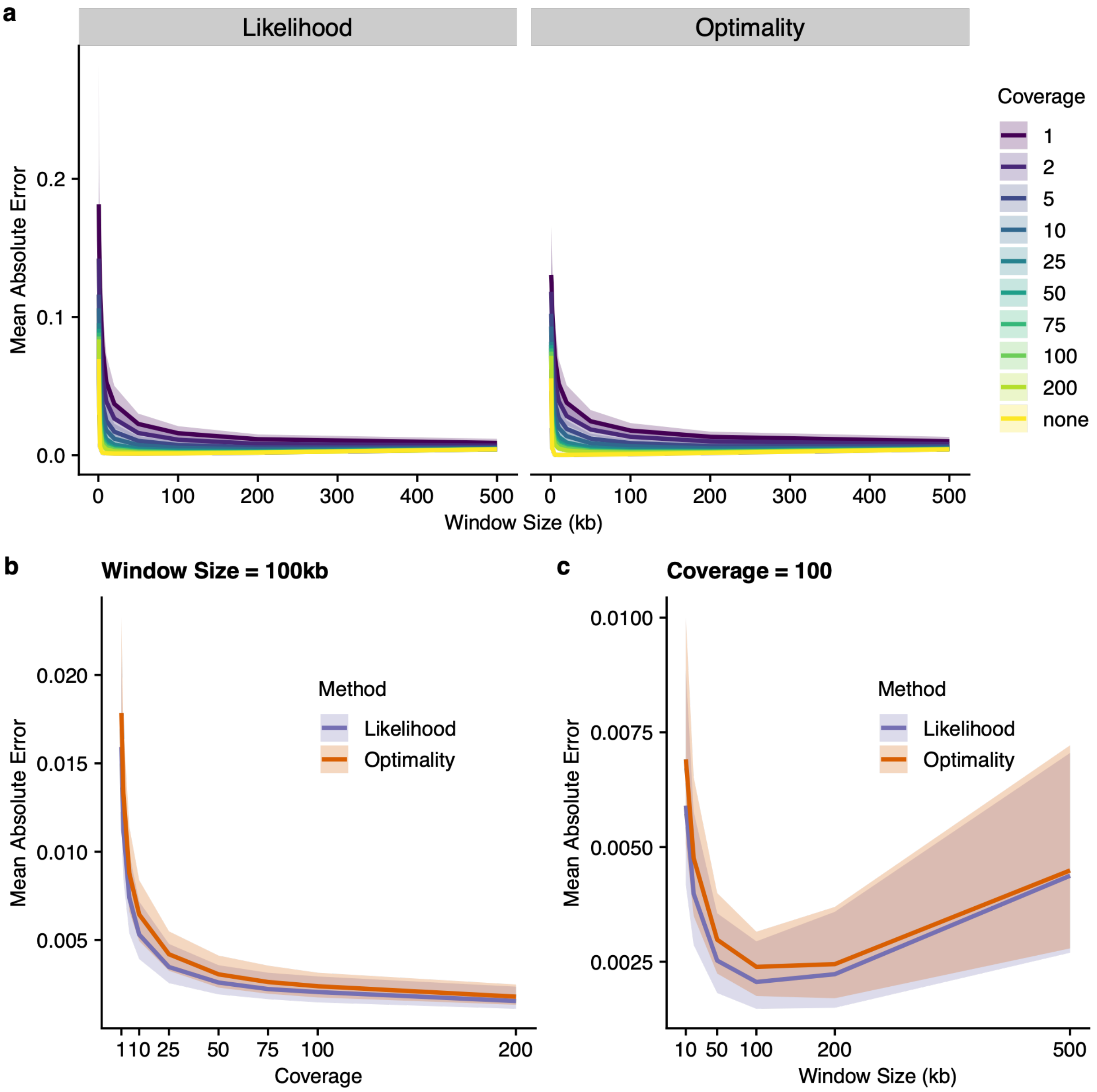


Figure S1: The mean absolute error for each of our estimation methods (likelihood and optimality) for a range of simulated coverage and window sizes. Note that each panel has its own range for the y-axis. a) The full range of parameter combinations with window size on the x-axis and different coverage values denoted by color. The "none" category of coverage is the case where coverage is not simulated. b) The effect of coverage for a window size of 100kb with the estimation method denoted by different colors. c) The effect of window size with coverage held constant at 100 with the estimation method denoted by different colors. Here, only window sizes greater than 10kb are shown.

Burke, M. K., E. G. King, P. Shahrestani, M. R. Rose, and A. D. Long. 2014. Genome-wide association study of extreme longevity in *Drosophila melanogaster*. Genome Biol. Evol. 6:1–11.

Kessner, D., T. L. Turner, and J. Novembre. 2013. Maximum Likelihood Estimation of Frequencies of Known Haplotypes from Pooled Sequence Data. Mol. Biol. Evol. 30:1145–1158.

King, E. G., S. J. Macdonald, and A. D. Long. 2012a. Properties and power of the *Drosophila* Synthetic Population Resource for the routine dissection of complex traits. Genetics 191:935–949.

King, E. G., C. M. Merkes, C. L. McNeil, S. R. Hoofer, S. Sen, K. W. Broman, A. D. Long, and S. J. Macdonald. 2012b. Genetic dissection of a model complex trait using the *Drosophila* Synthetic Population Resource. Genome Res. 22:1558–1566.

Linder, R. A., B. Zabanavar, A. Majumder, H. C.-S. Hoang, V. G. Delgado, R. Tran, V. T. La, S. W. Leemans, and A. D. Long. 2022. Adaptation in Outbred Sexual Yeast is Repeatable, Polygenic and Favors Rare Haplotypes. Mol. Biol. Evol. 39:msac248.

Macdonald, S. J., K. M. Cloud-Richardson, D. J. Sims-West, and A. D. Long. 2022. Powerful, efficient QTL mapping in Drosophila melanogaster using bulked phenotyping and pooled sequencing. Genetics 220:iyab238.

Tilk, S., A. Bergland, A. Goodman, P. Schmidt, D. Petrov, and S. Greenblum. 2019. Accurate Allele Frequencies from Ultra-low Coverage Pool-Seq Samples in Evolve-and-Resequence Experiments. G3 GenesGenomesGenetics 9:4159–4168.

Ward, M. J., S. A. Ellsworth, E. G. King, E. Ng'oma, G. S. Nystrom, K. C. Lawrence, L. Maquet-Diafouka, A. Oliver, M. J. Margres, C. L. Parkinson, K. A. Hughes, and D. R.

Rokyta. 2026. The genetic architecture of venom resistance: a novel approach to target identification and coevolutionary discovery. Mol. Biol. Evol. 43:msag048.

### Additional supplementary figures

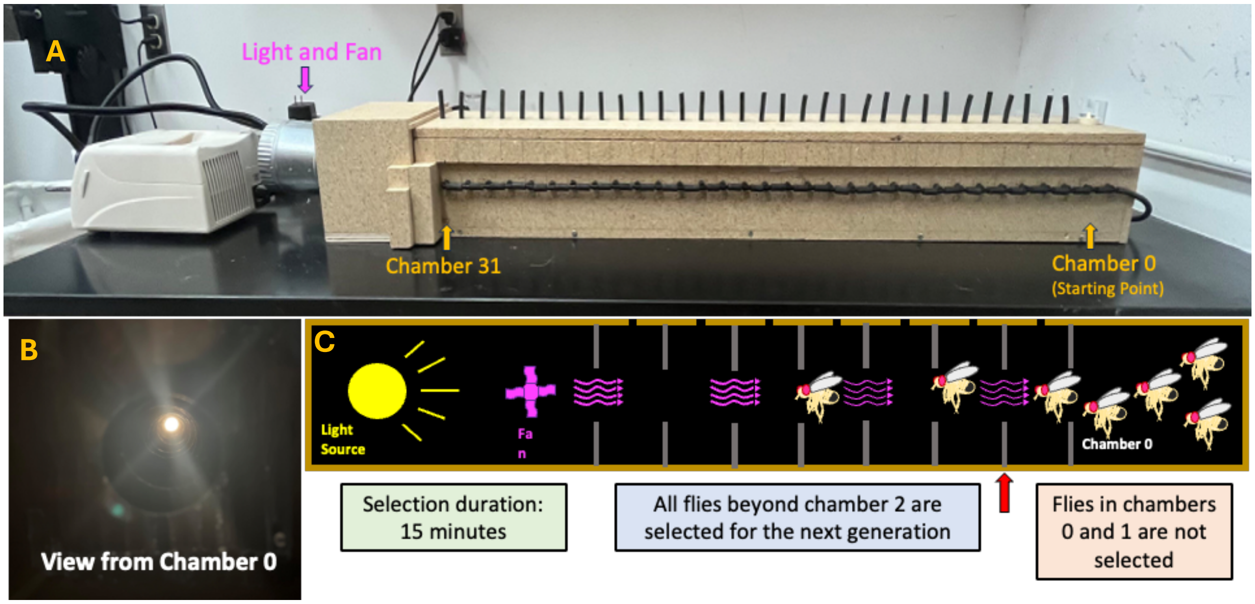


Supplementary figure S2: Custom built wind tunnel. A. Photo of the setup. B. Photo from inside the setup, with lights on. C. Schematic of selection process.

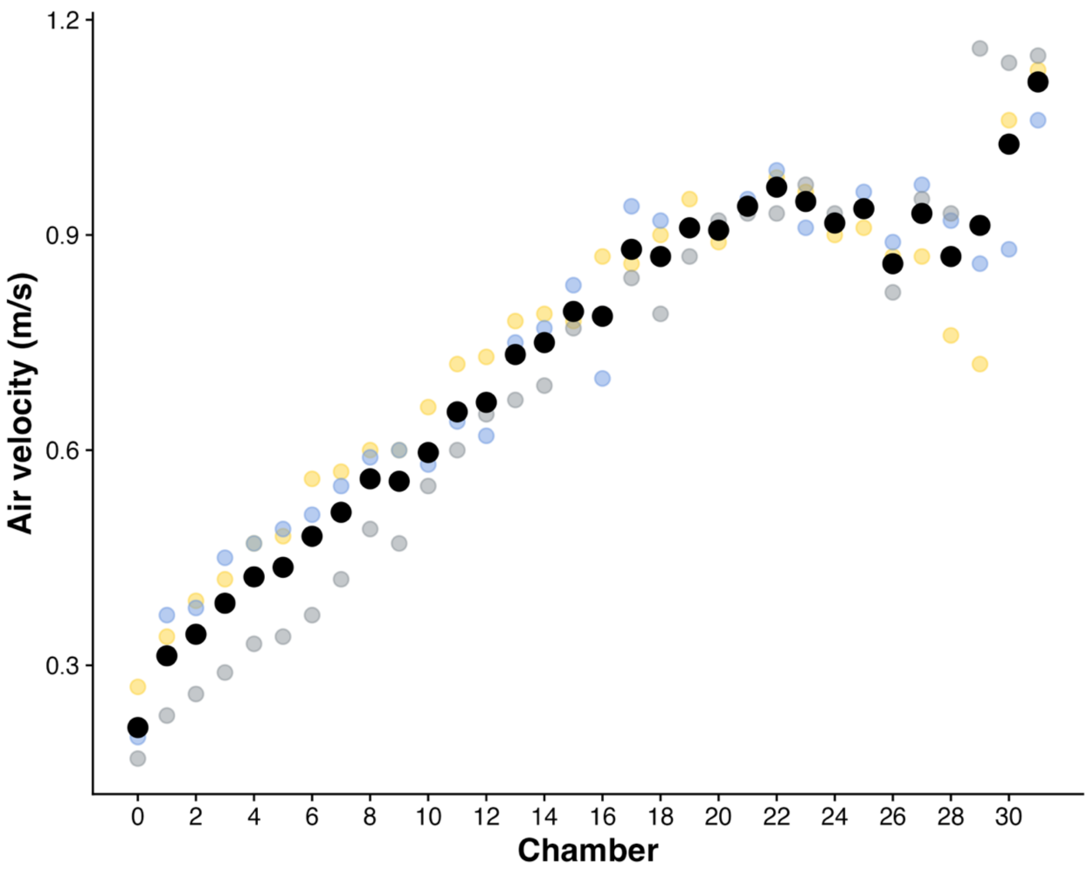


Supplementary figure S3. Air velocity per chamber within the flight tunnel setup. Flies are added in chamber 0. Chamber 32 contains the lights and fan.

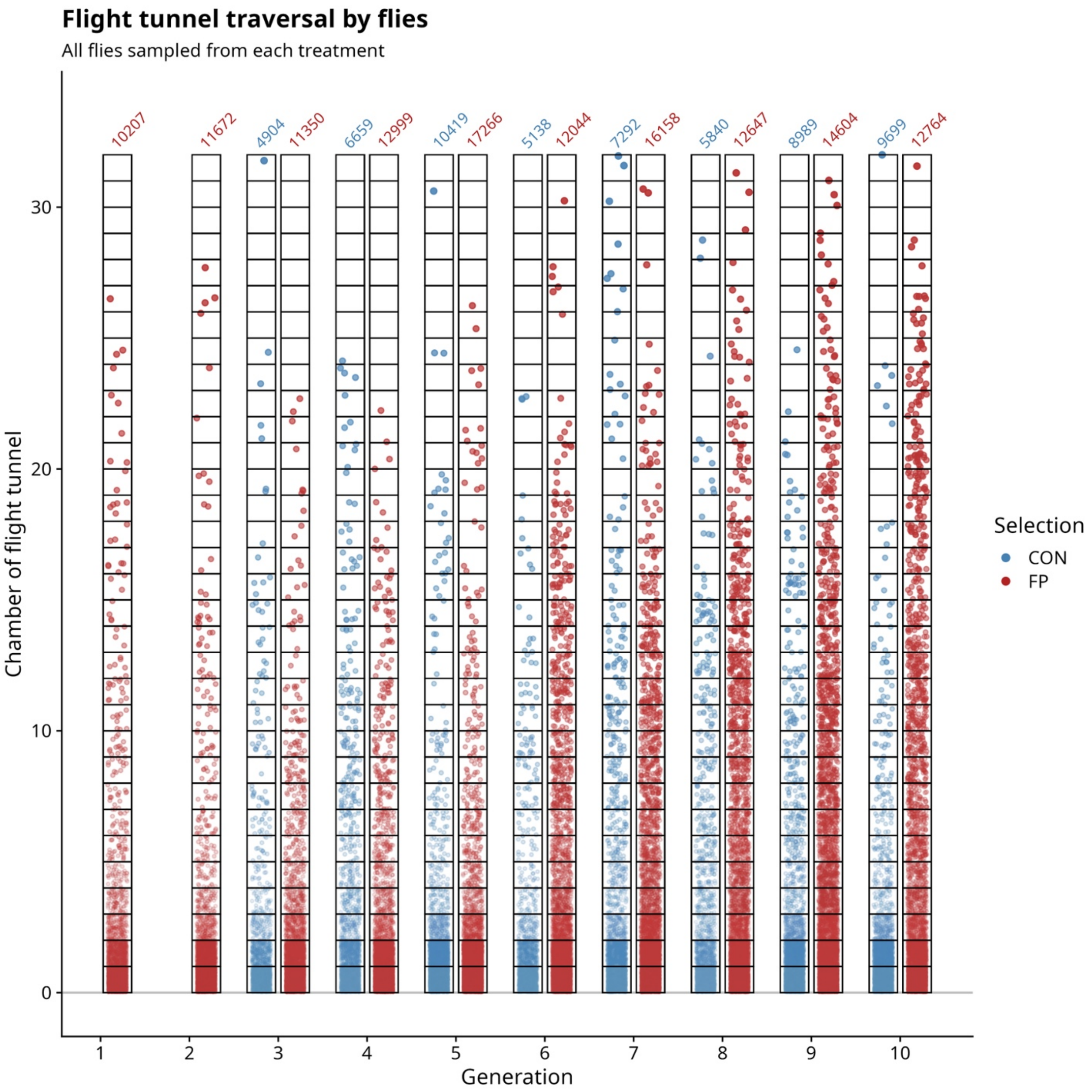


Supplementary figure S4. The evolution of flight performance across 10 generations of selection: The distance traversed by flies within the flight tunnel, pooling across replicate populations and runs. The X-axis shows generation number of selection. The Y-axis denotes the chamber position for all sampled flies each combination of Selection × Generation. Above each scatter, the number denotes the sample size.

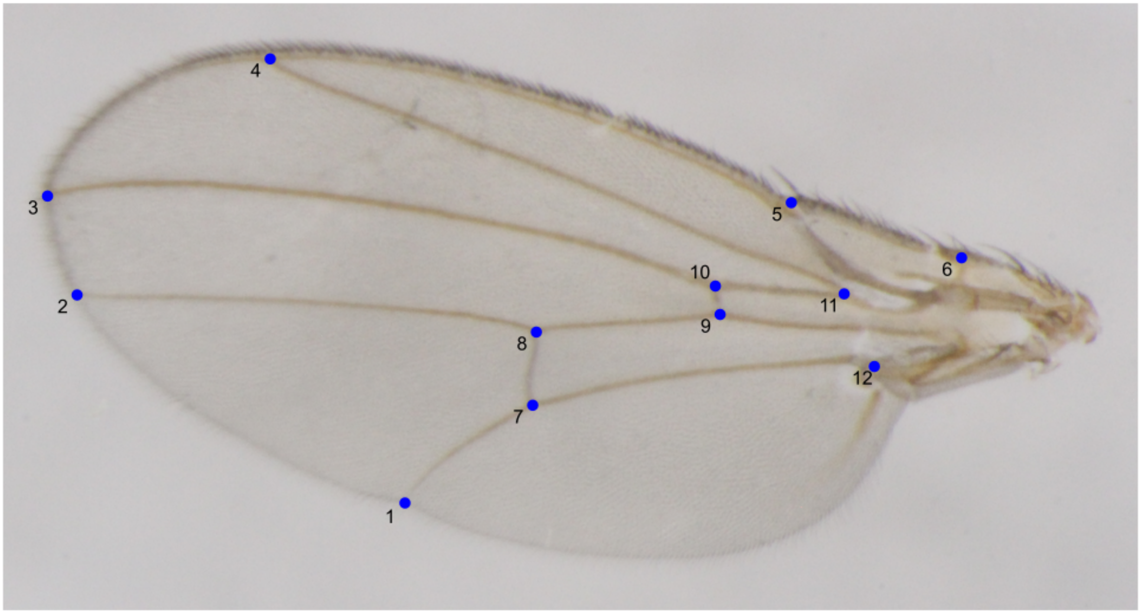


Supplementary figure S5. Landmarks used for wing landmarking. Landmarks were assigned following those used by Houle et al. (2003), which cover most of the longitudinal and cross veins. We chose not to use semilandmarks around the outer edge of the wing.

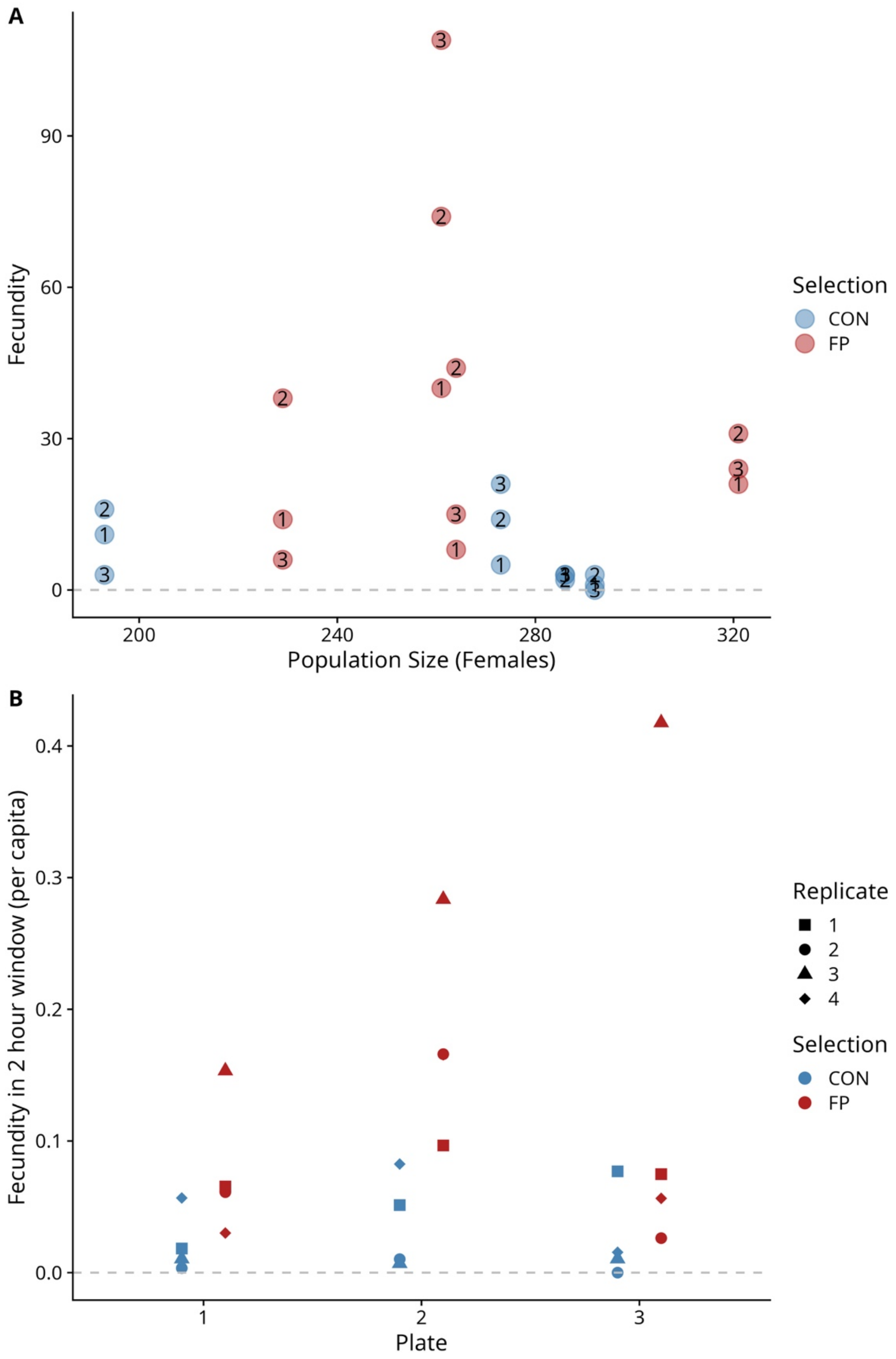


Supplementary figure S6. A. Relationship between Fecundity and populations size of females. X-axis shows female populations size. Y-axis shows total measured fecundity. B. Relationship between per-capita fecundity and plate number. X-axis shows plate number, Y-axis shows per-capita fecundity, measured in a 2-hour window.

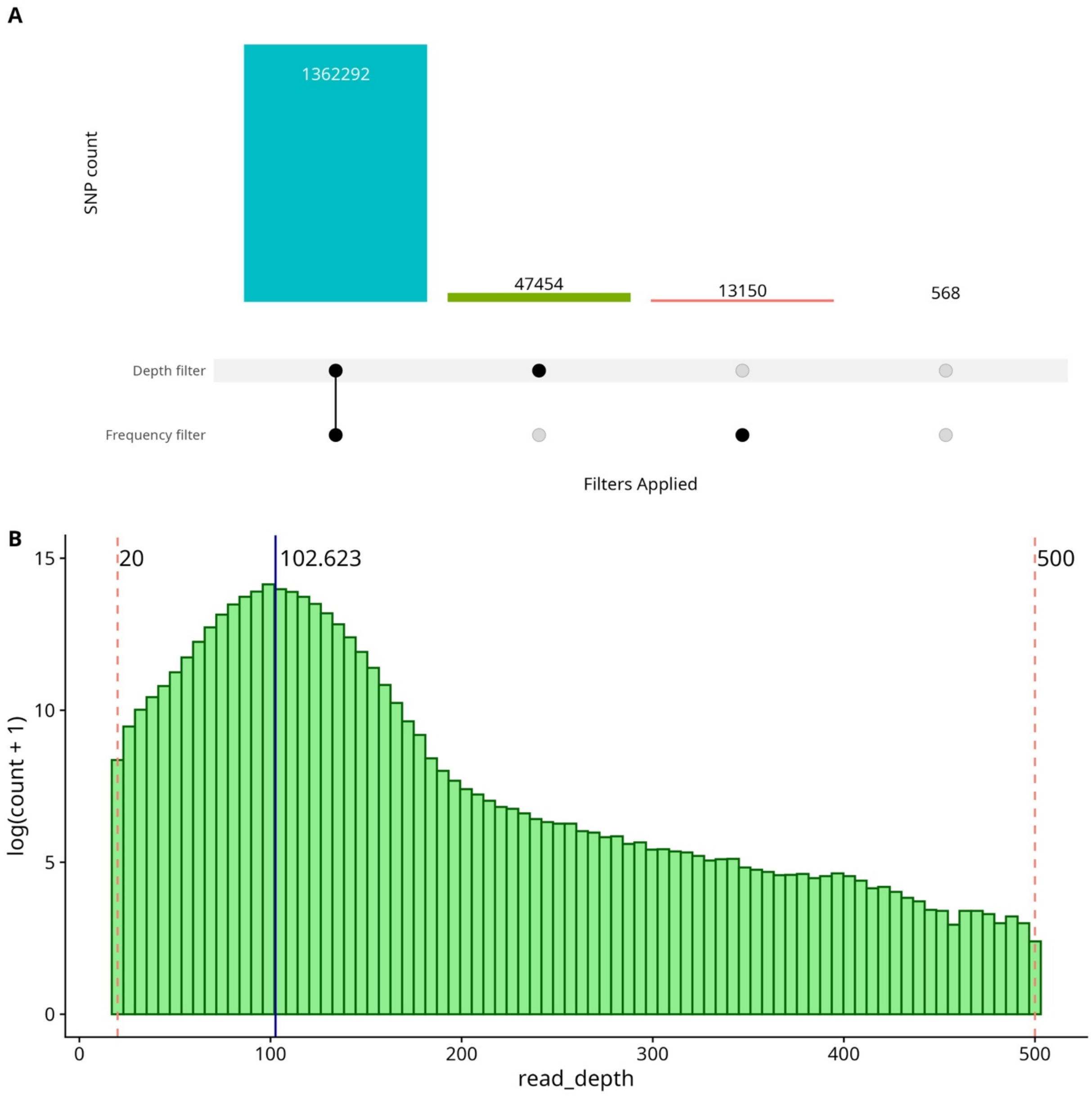


Supplementary figure S7. A. Upset plot on filtering counts. The bar plots respectively show number of SNPs when both read depth and frequency filter are applied, only depth filter applied, only frequency filter applied, and the counts which are excluded from both read depth and frequency filters. The SNPs used are the ones from both depth and frequency filters applied (1,362,292). B. Histogram of read depths after filtering. The X-axis shows read depth, and Y-axis shows the log of counts for each given read depth. The red vertical dashed lines show the minimum and maximum filtering criteria, respectively. The dark blue vertical line shows the mean read depth after filtering (102.623).

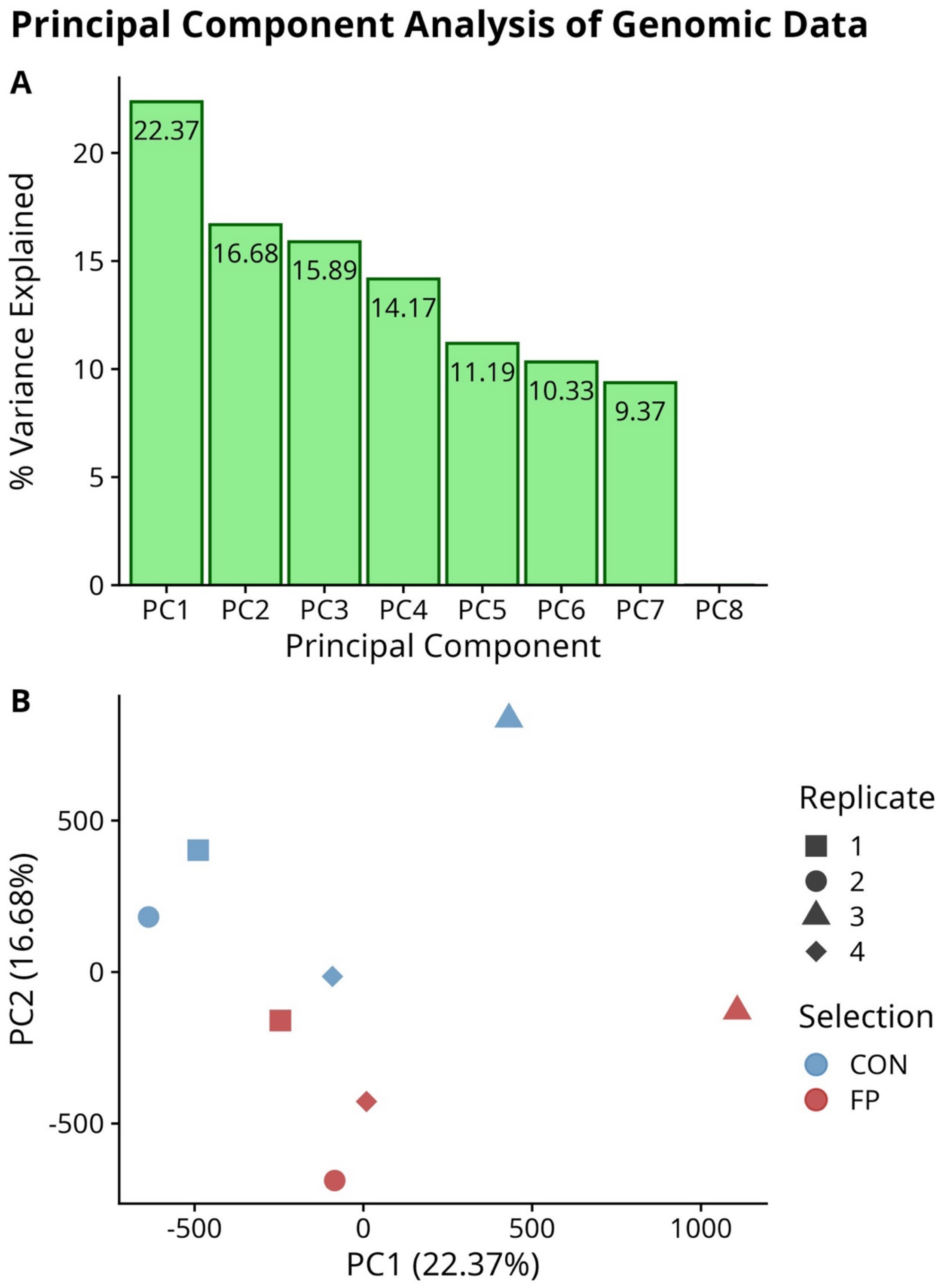


Supplementary figure S8. Results from Principal Component Analysis on SNP frequency data for the 8 populations (4 CON, 4 FP). A. Scree plot showing percentage of variance captured by each principal component. B. Biplot of PC1 and PC2, showing the scatter of populations. Points close to each other denote closeness in genome sequences.

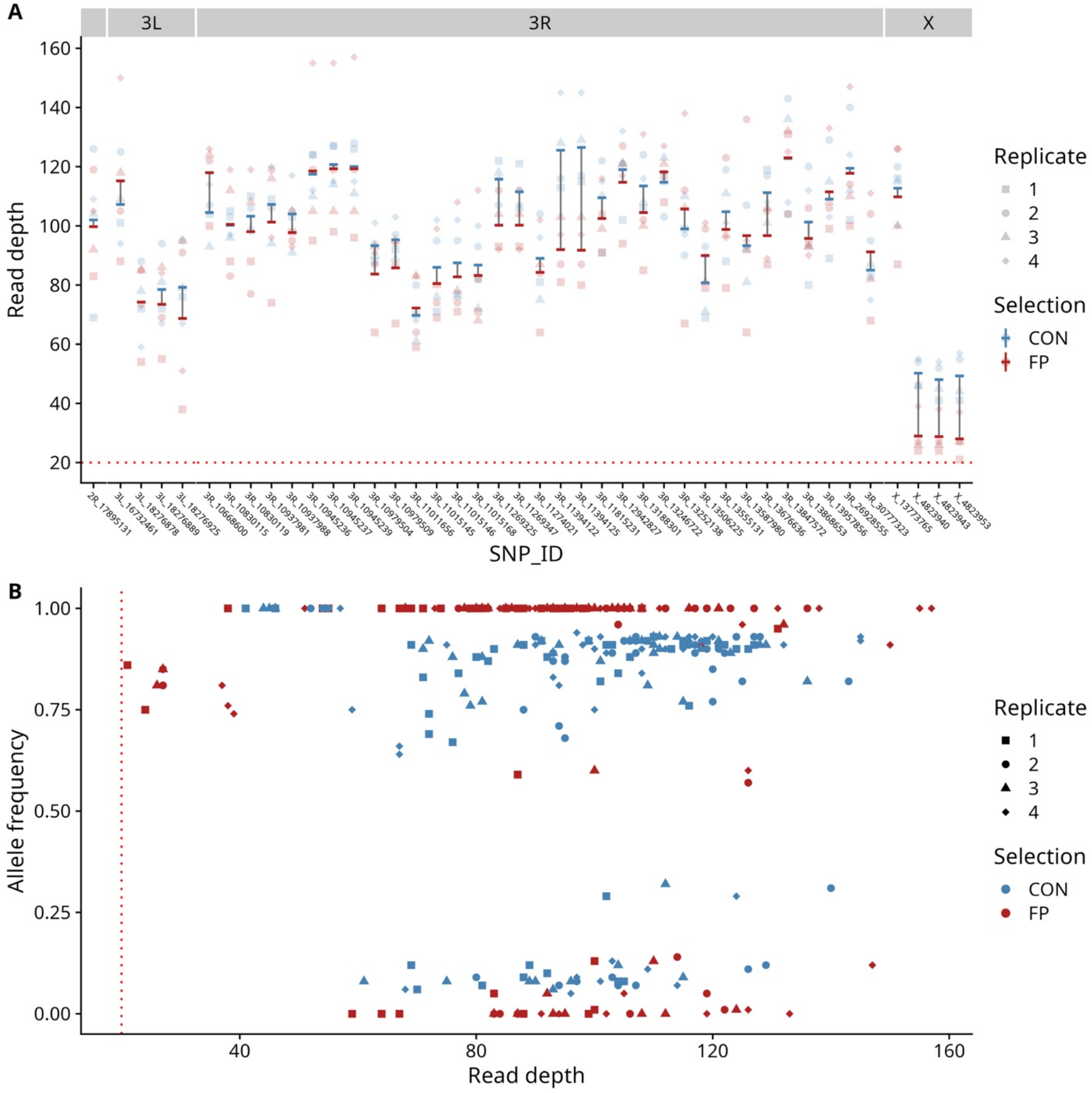


Supplementary figure S9. A. Read depth distribution for each significantly differentiated site from the linear model analysis. The X-axis shows The SNP ID, and the Y-axis shows the read depth. Populations are colored by Selection (CON or FP) and shape denotes population replicate. The opaque '-' points denote mean for Selection at the given SNP ID, and the vertical grey line denotes difference between CON and FP. B. Scatter plot between reference allele frequency and read depth of significant sites. The vertical red dotted line denotes the minimum cutoff for read depth (20).

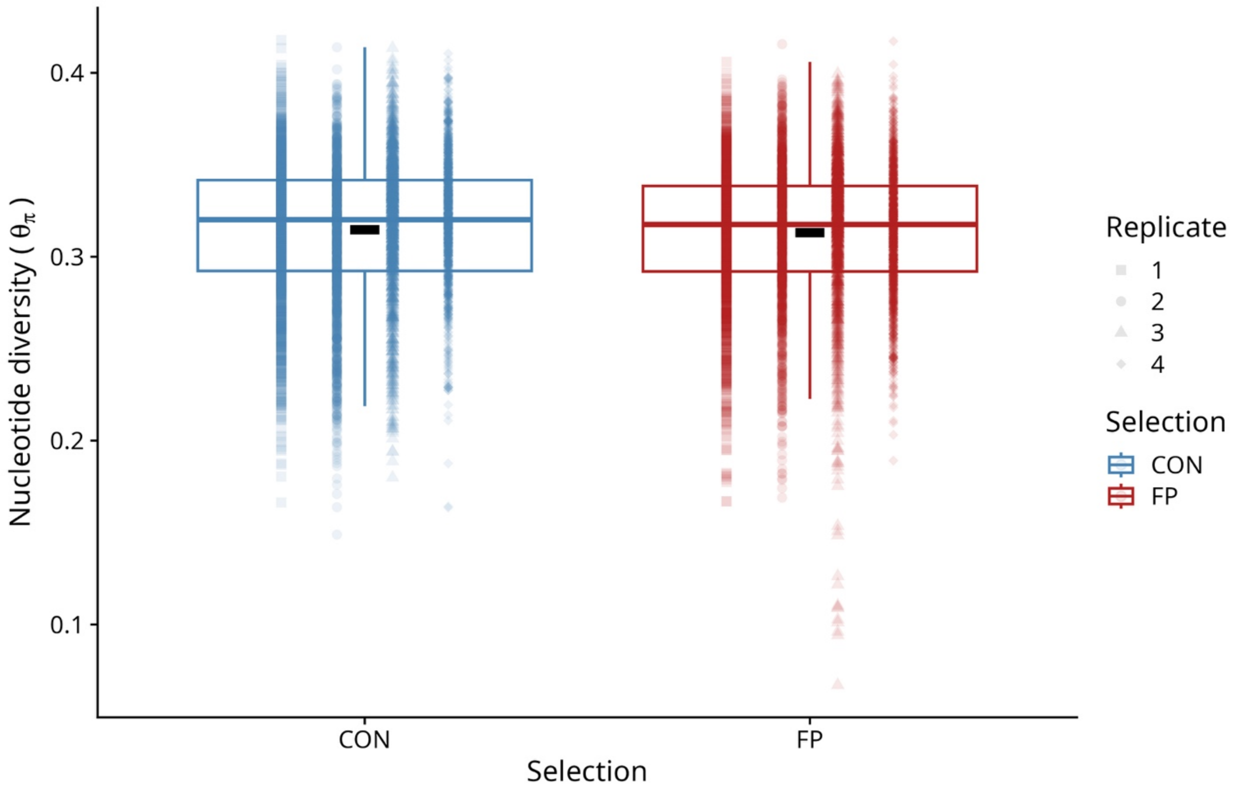


Supplementary figure S10. Boxplots of nucleotide diversity, for CON and FP populations. Y-axis shows nucleotide diversity, X-axis shows Selection (CON or FP). The black '-' denotes overall mean for CON or FP.

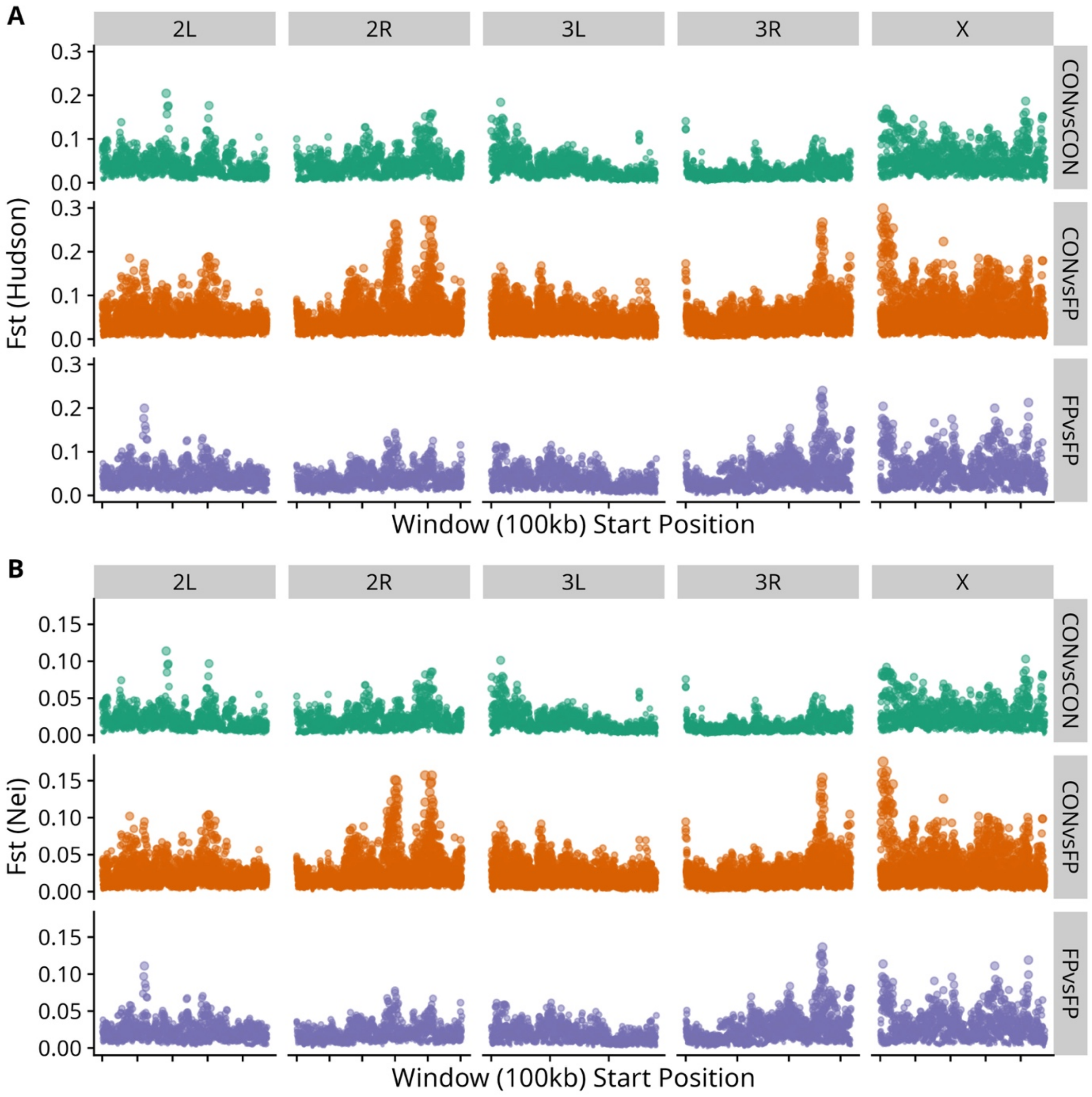


Supplementary figure S11. Fst values for different comparison categories. Y-axis shows Fst value, X-axis shows window position (start of window). A. Fst Hudson estimate. B. Fst Nei estimate.

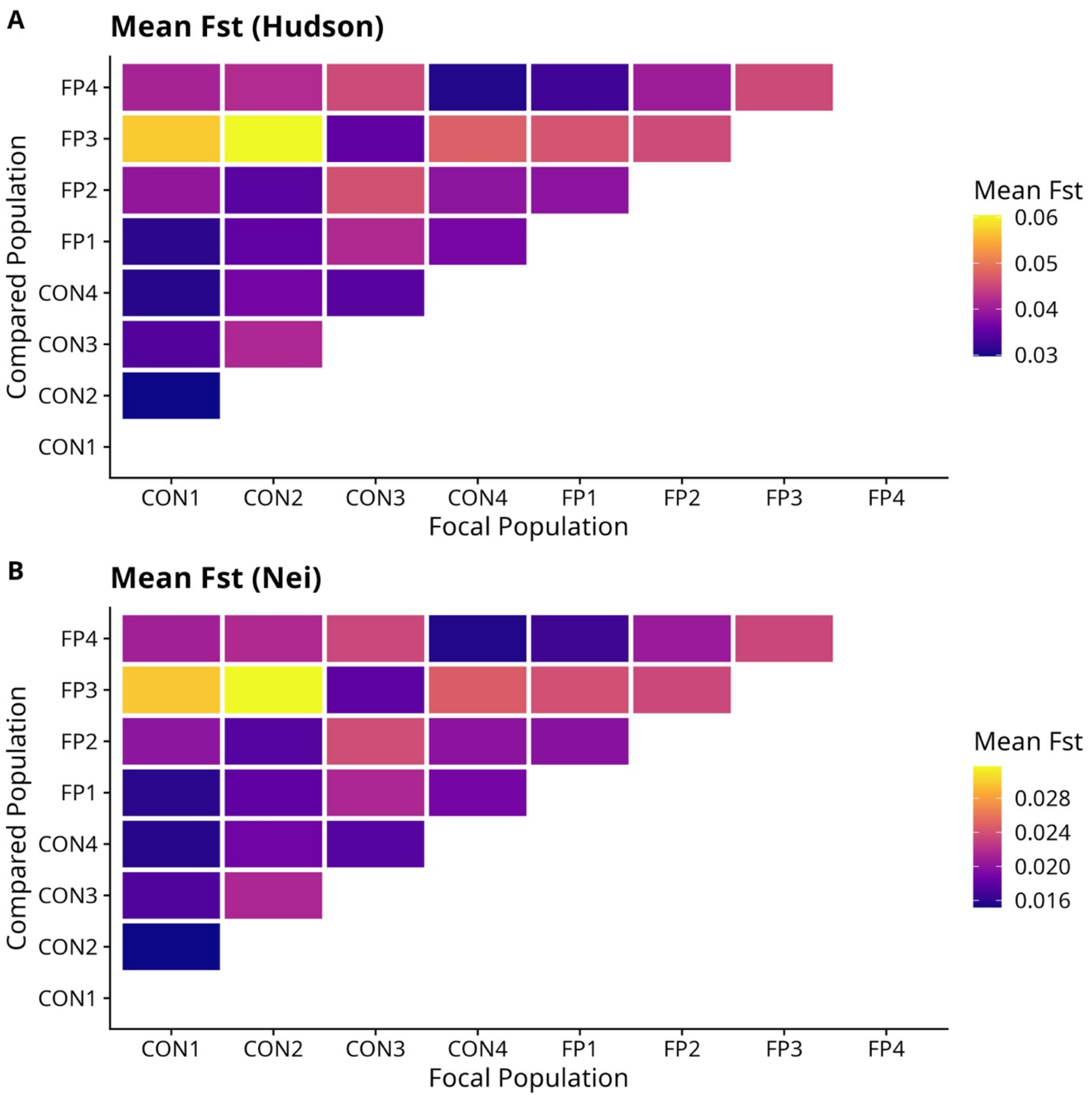


Supplementary figure S12. Mean Fst value heatplots for population-wise comparisons. A. Fst Hudson estimate. B. Fst Nei estimate.

Gene ontology and enrichment

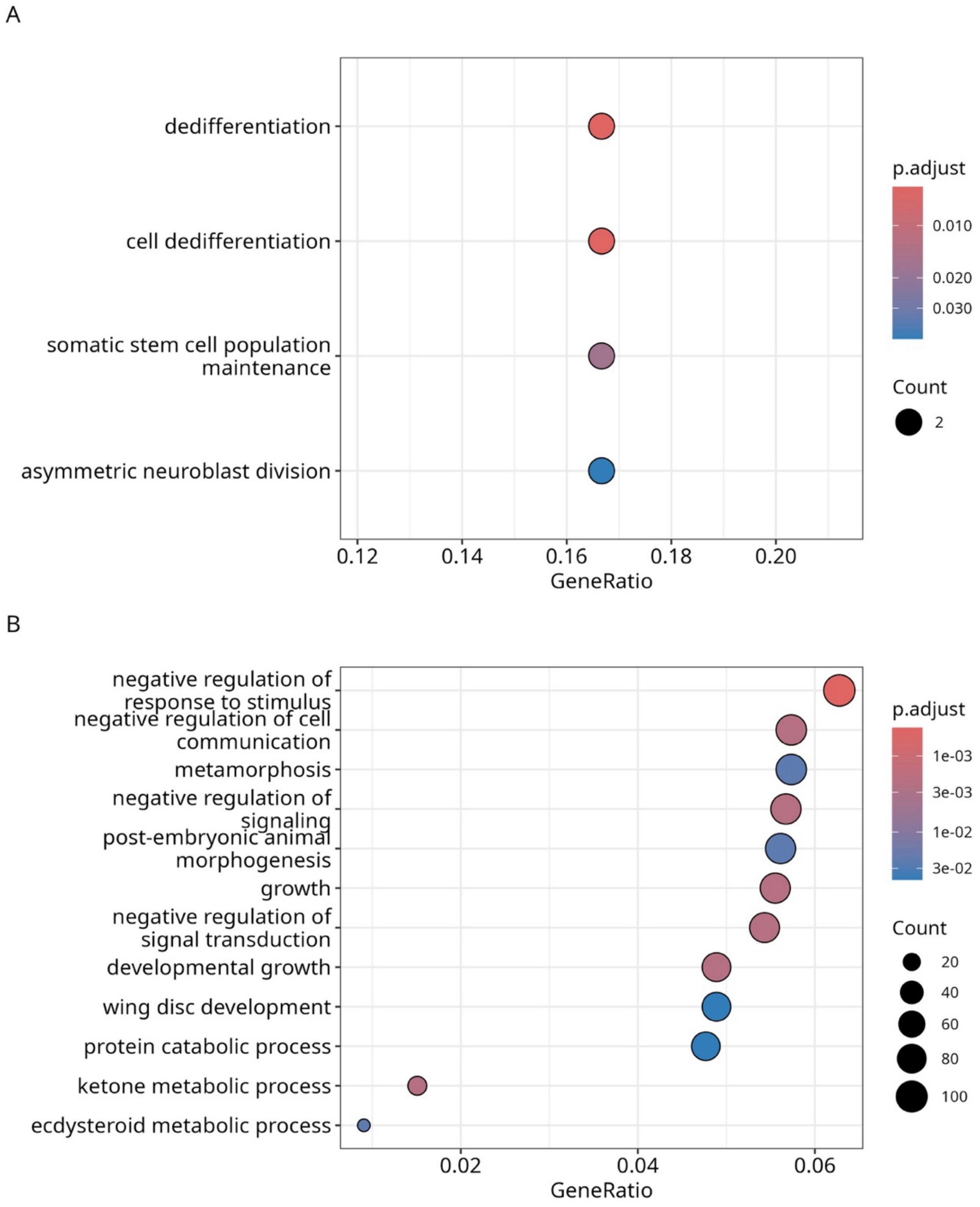


Supplementary figure S13: Gene Ontology terms enriched for Biological Process. The size of the dot indicates number of genes associated with the pathway. The color is the adjusted P-value after Benjamini Hochberg correction. The X-axis shows the Gene Ratio, which is the gene count for a given dot divided by the total number of genes analyzed. The Y-axis shows the different pathways enriched from the GO terms. A. Shows the pathways enriched from significant sites of the linear model test (fig. 7). B. Shows the pathways enriched from significant sites based on haplotype frequency differences (fig. S14-18).

**The following supplementary figures (S14-18) denote haplotype analyses results for entire chromosome arms.**

A. Standardized haplotype frequency ascribed to each founder line, for each population, across genomic sites. The X-axis denotes genomic position. The Y-axis denotes standardized frequency for each population, colored by Selection (CON or FP). The 8 panels are labelled for each founder haplotype (A1-A8). The bottom panel marked 'SNP' displays the standardized reference allele frequency for the given population and genomic position. Dark grey background lines denote sites found to differ significantly between FP and CON populations. Yellow background lines show for those sites found different in the linear model test (fig. 7). B. For each significant genomic position, we calculated the average difference between the CON and FP populations for the frequency of each founder. We ranked each founder by magnitude of difference at each site, with the greatest magnitude of differences ranked 1, and the smallest magnitude ranked 8. Thus, on each chromosomal arm, we could calculate how many times each founder was rank 1, how many times it was rank 2 and so on. Mean rank for each founder is shown on top of each respective bar. C. The genomic sites found to differ significantly in haplotype frequency between CON and FP populations. The X-axis denotes genomic position. The Y-axis shows -log10(FDR adjusted *P*-values). The mapped genes are labelled and colored according to enriched pathways (1 random pathway in case a gene maps to multiple pathways).

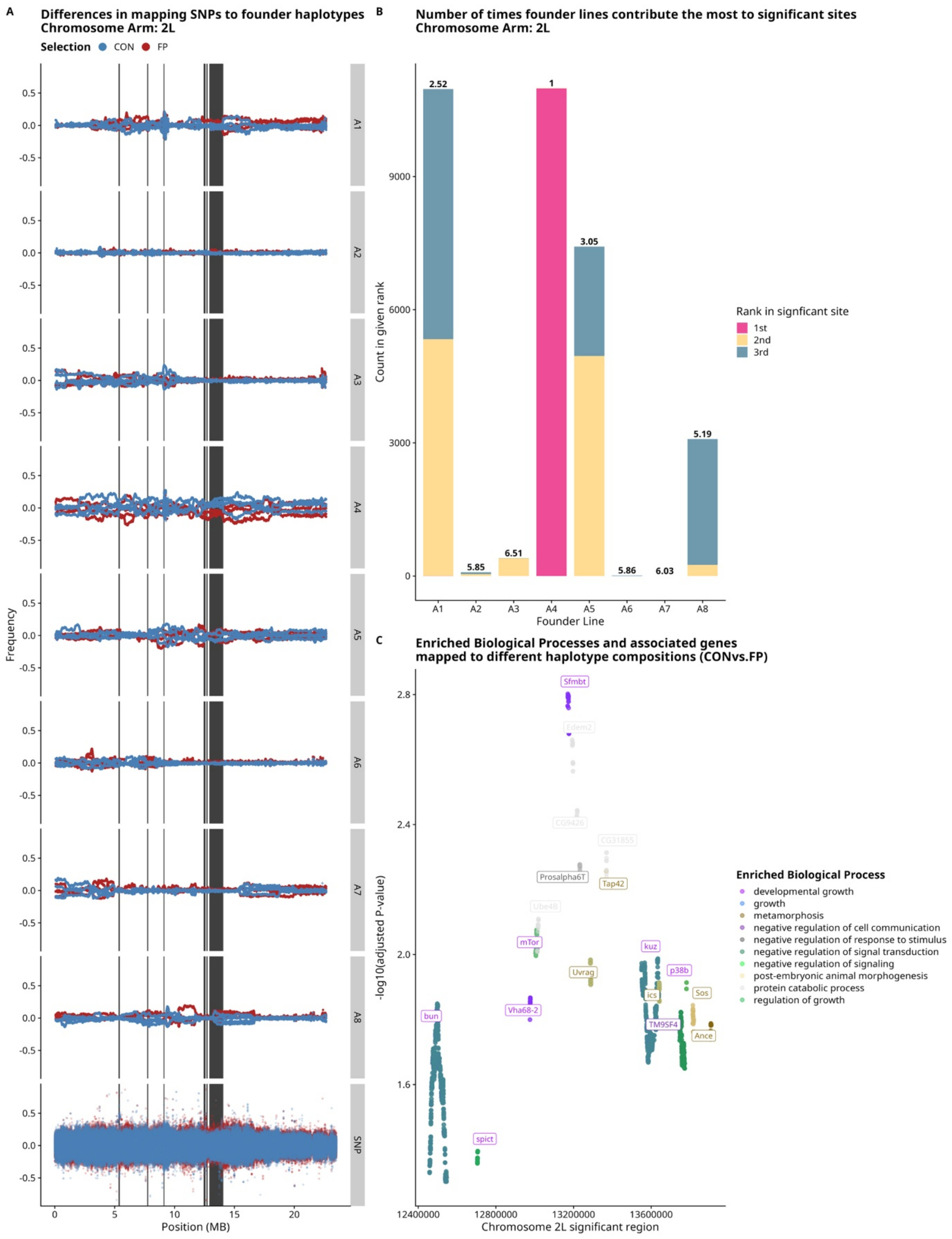


Supplementary figure S14. Haplotype analyses for chromosome arm 2L.

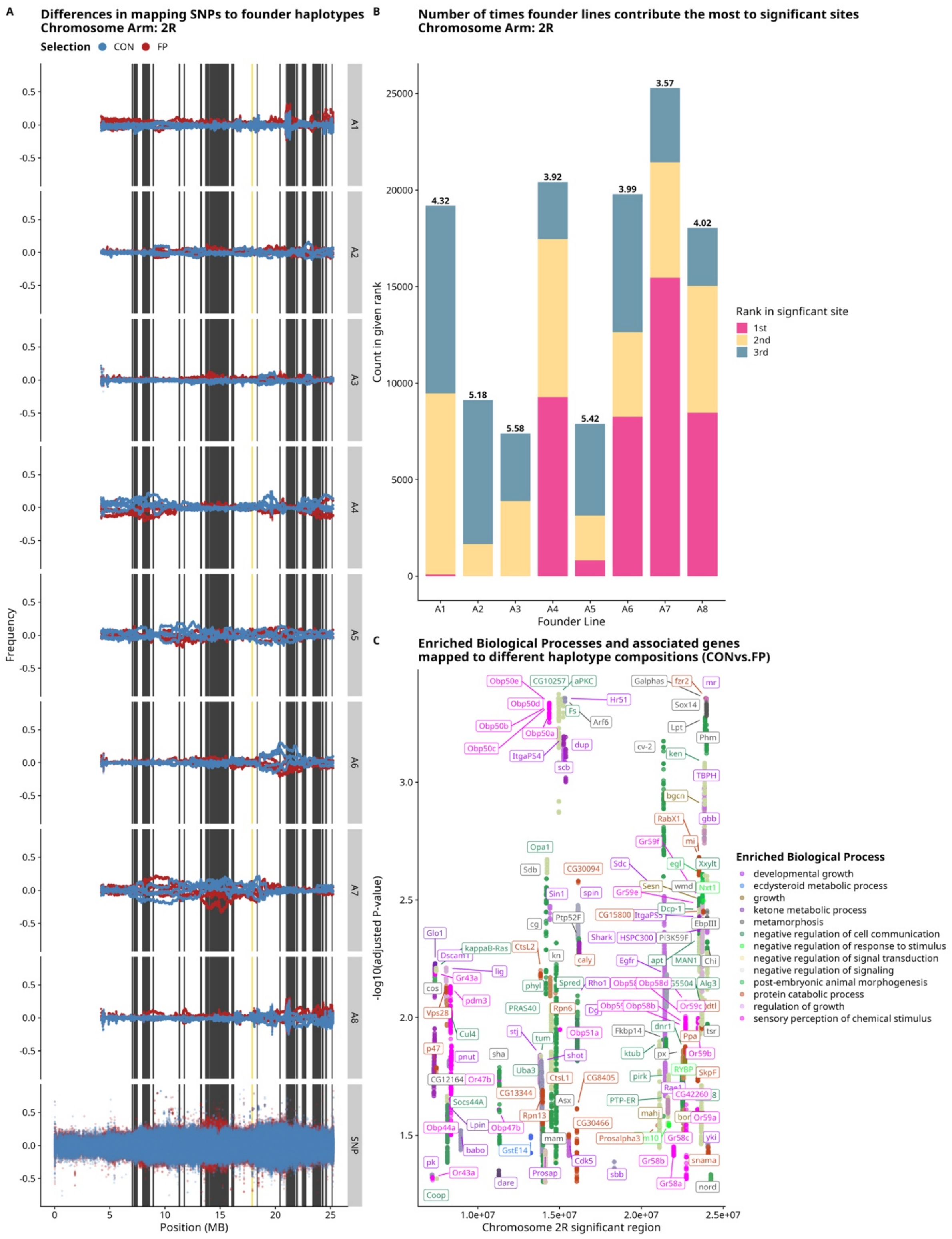


Supplementary figure S15. Haplotype analyses for chromosome arm 2R.

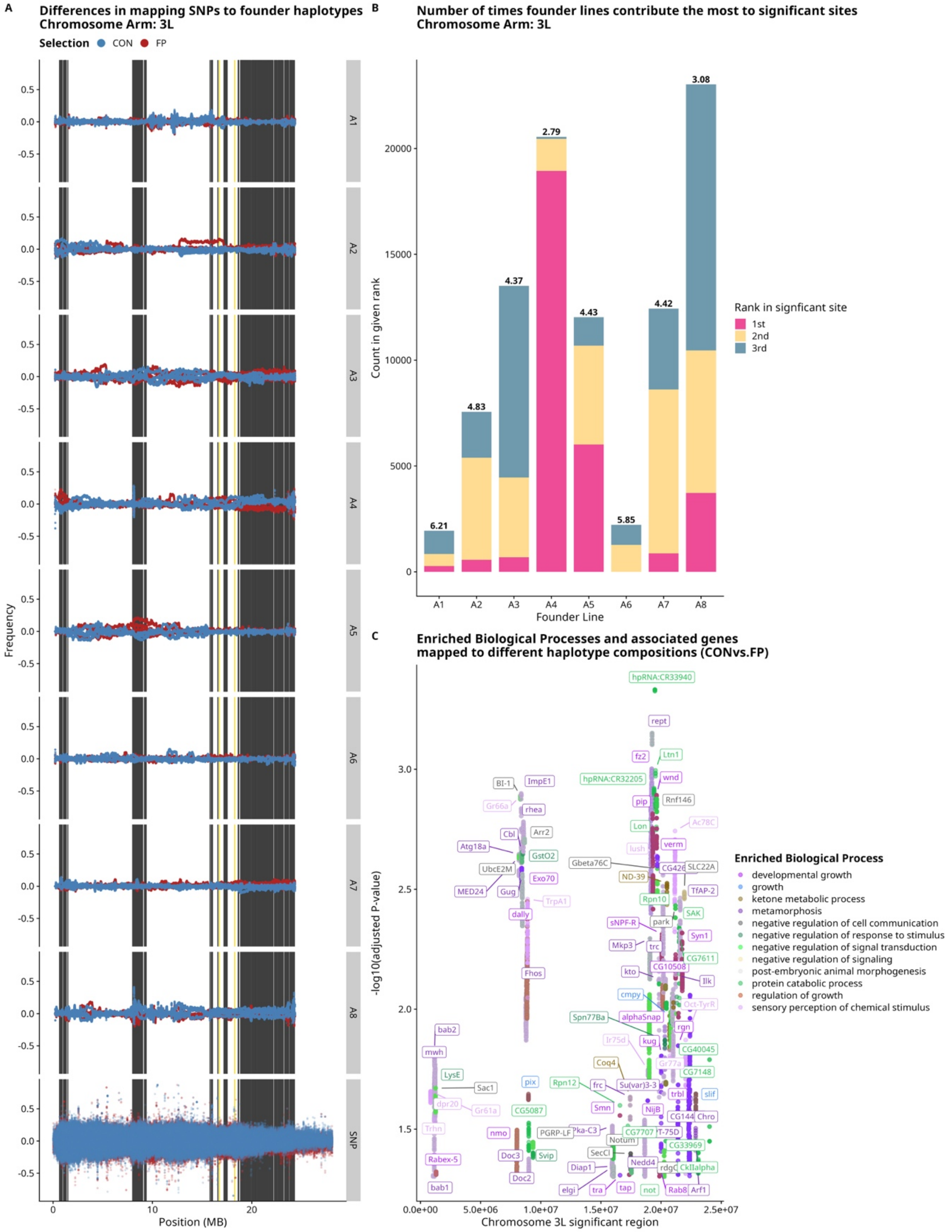


Supplementary figure S16. Haplotype analyses for chromosome arm 3L.

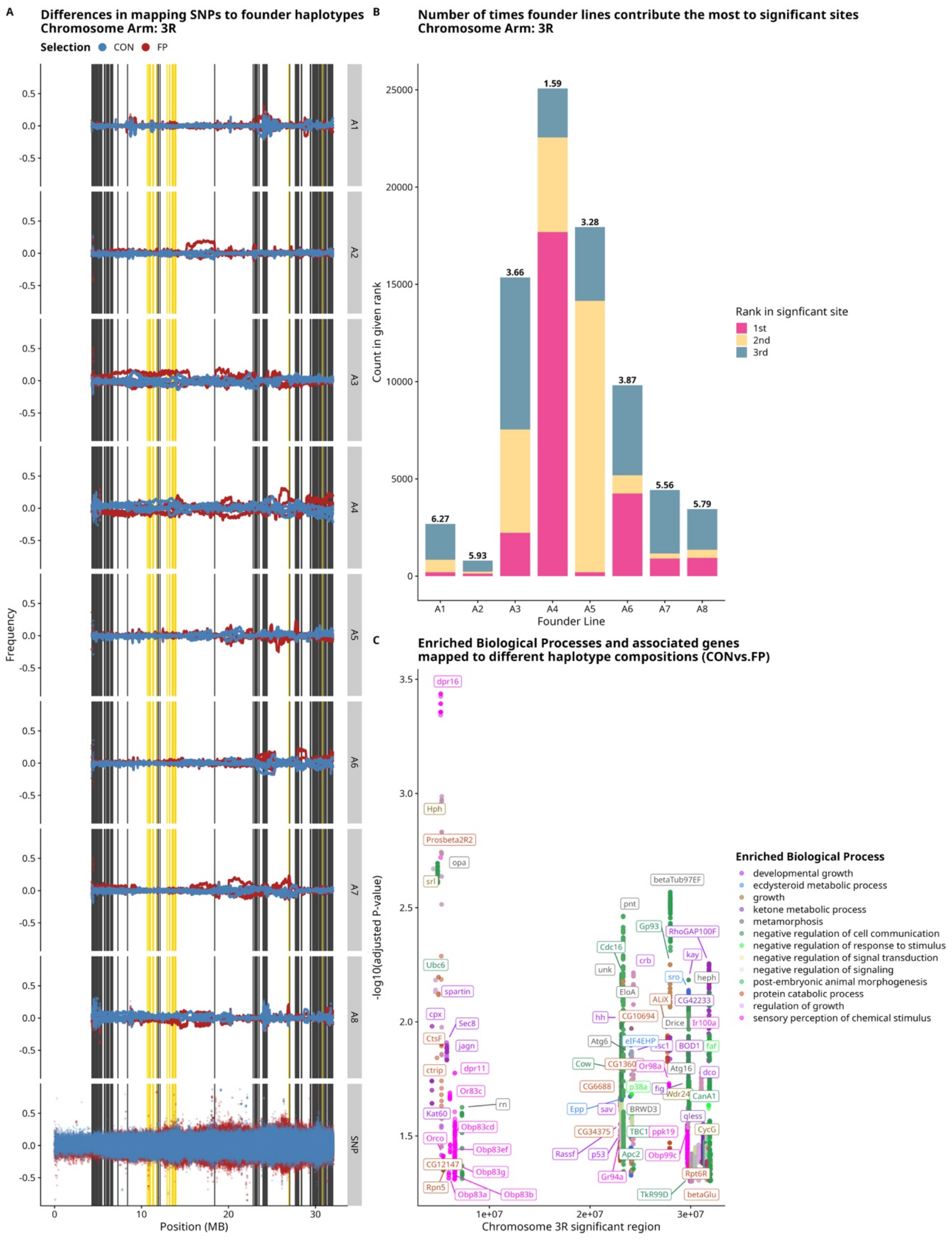


Supplementary figure S17. Haplotype analyses for chromosome arm 3R.

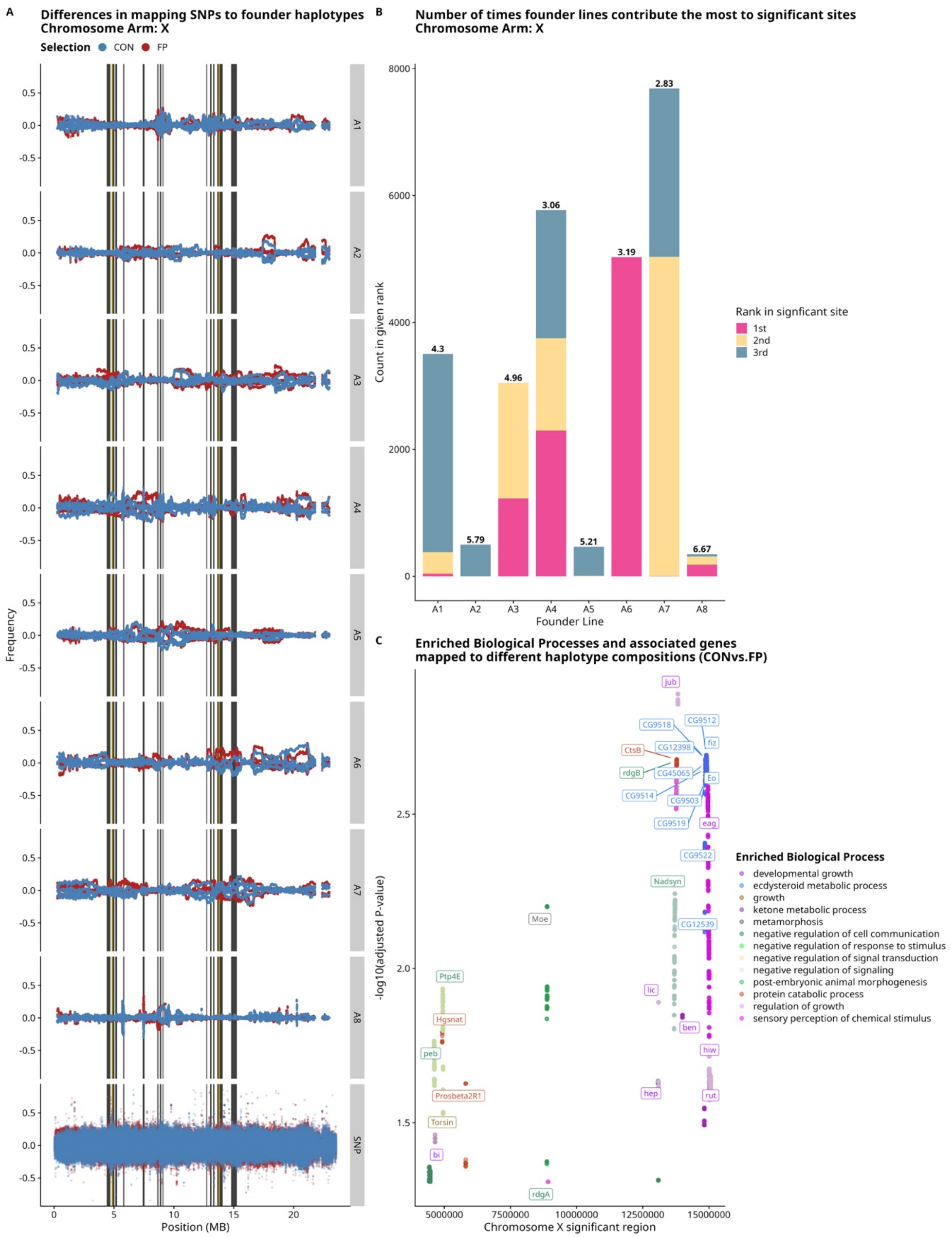


Supplementary figure S17. Haplotype analyses for chromosome arm X.

**Supplemental tables**

| | | | | |
|---|---|---|---|---|
| 11004 | 11009 | 11012 | 11016 | 11037 |
| 11038 | 11044 | 11052 | 11062 | 11074 |
| 11110 | 11117 | 11122 | 11149 | 11153 |
| 11155 | 11159 | 11162 | 11185 | 11192 |
| 11196 | 11216 | 11228 | 11232 | 11239 |
| 11255 | 11265 | 11298 | 11299 | 11300 |
| 11303 | 11305 | 11310 | 11316 | 11321 |
| 11328 | 11355 | 11358 | 11360 | 11361 |
| 11369 | 11374 | 11375 | 11376 | 11381 |
| 11410 | 11416 | 11420 | 11464 | 11465 |
| 11466 | 11481 | 12002 | 12010 | 12014 |
| 12021 | 12022 | 12028 | 12039 | 12043 |
| 12050 | 12075 | 12080 | 12097 | 12114 |
| 12123 | 12130 | 12138 | 12145 | 12159 |
| 12169 | 12190 | 12191 | 12234 | 12243 |
| 12253 | 12258 | 12270 | 12272 | 12278 |
| 12279 | 12285 | 12300 | 12301 | 12304 |
| 12316 | 12333 | 12348 | 12357 | 12363 |
| 12372 | 12381 | 12390 | | |

Supplementary table S1: The DSPR RIL IDs used to create the outbred populations in the study.

| *Model* | *Effect* | *SS* | *MS* | *dfnum* | *dfden* | *F* | *P* | *adj. P* |
|---|---|---|---|---|---|---|---|---|
| Selection × Gen | Selection | 0.7576 | 0.7576 | 1.0000 | 8.1899 | 22.8834 | 0.0013 | 0.0036 |
| Selection × Gen | Generation | 2.3888 | 0.2654 | 9.0000 | 48.0175 | 8.0166 | 0.0000 | 0.0000 |
| Selection × Gen | Selection:Generation | 1.0410 | 0.1487 | 7.0000 | 47.8321 | 4.4916 | 0.0007 | 0.0019 |
| Gen (only CON) | Generation | 0.2479 | 0.0354 | 7.0000 | 24.0000 | 1.0325 | 0.4349 | 0.5415 |
| Gen (only FP) | Generation | 3.1609 | 0.3512 | 9.0000 | 26.0778 | 12.0759 | 0.0000 | 0.0000 |

Supplementary table S2: ANOVA tables for evolution of flight performance

| *Model* | *Parameter* | *Eta2_partial* | *CI* | *CI_low* | *CI_high* |
|---|---|---|---|---|---|
| Selection × Gen | Selection | 0.7364 | 0.9500 | 0.3650 | 1.0000 |
| Selection × Gen | Generation | 0.6004 | 0.9500 | 0.4013 | 1.0000 |
| Selection × Gen | Selection:Generation | 0.3966 | 0.9500 | 0.1507 | 1.0000 |
| Gen (only CON) | Generation | 0.2314 | 0.9500 | 0.0000 | 1.0000 |
| Gen (only FP) | Generation | 0.8065 | 0.9500 | 0.6502 | 1.0000 |

Supplementary table S3: partial eta-squared tables for evolution of flight performance

| *Model formula* | *K* | *AICc* | *Delta_AICc* | *AICc weight* | *LL* |
|---|---|---|---|---|---|
| 4.Additive | 6.0000 | 126.8987 | 0.0000 | 0.8071 | -56.7125 |
| 5.SelCross | 8.0000 | 131.1905 | 4.2918 | 0.0944 | -56.2862 |
| 1.mainSel | 4.0000 | 131.5248 | 4.6262 | 0.0799 | -61.4234 |
| 3.mainFan | 4.0000 | 136.4729 | 9.5743 | 0.0067 | -63.8975 |
| 6.Fullmod | 10.0000 | 136.6868 | 9.7881 | 0.0060 | -56.2679 |
| 2.mainLight | 4.0000 | 136.7532 | 9.8546 | 0.0058 | -64.0376 |

Supplementary table S4: AICc results for models fit for factorial flight performance assay

| *Effect* | *SS* | *MS* | *dfnum* | *dfden* | *F* | *P* | *adj. P* |
|---|---|---|---|---|---|---|---|
| Selection | 4.3668 | 4.3668 | 1.0000 | 6.0000 | 13.9243 | 0.0097 | 0.0214 |
| Light_Status | 1.5052 | 1.5052 | 1.0000 | 54.0000 | 4.7997 | 0.0328 | 0.0577 |
| Fan_Status | 1.5978 | 1.5978 | 1.0000 | 54.0000 | 5.0947 | 0.0281 | 0.0529 |

Supplementary table S5: ANOVA table for factorial flight performance assay

| *Parameter* | *Eta2_partial* | *CI* | *CI_low* | *CI_high* |
|---|---|---|---|---|
| Selection | 0.6989 | 0.9500 | 0.2043 | 1.0000 |
| Light_Status | 0.0816 | 0.9500 | 0.0037 | 1.0000 |
| Fan_Status | 0.0862 | 0.9500 | 0.0052 | 1.0000 |

Supplementary table S6: partial eta-squared table for factorial flight performance assay

| *Model formula* | *K* | *AICc* | *Delta_AICc* | *AICc weight* | *LL* |
|---|---|---|---|---|---|
| 5.Sel+Sex+Assay | 7.0000 | 19513.8632 | 0.0000 | 0.5518 | -9749.8921 |
| 6.Sel*Sex+Assay | 8.0000 | 19514.7181 | 0.8549 | 0.3599 | -9749.3082 |
| 7.Sel*Sex*Assay | 10.0000 | 19517.5569 | 3.6937 | 0.0870 | -9748.7007 |
| 8.Assay_random | 7.0000 | 19525.9507 | 12.0875 | 0.0013 | -9755.9359 |
| 3.Sel+Sex | 5.0000 | 19619.7363 | 105.8731 | 0.0000 | -9804.8470 |
| 4.Sel*Sex | 6.0000 | 19620.6260 | 106.7627 | 0.0000 | -9804.2834 |
| 2.Only_Sex | 4.0000 | 19624.8713 | 111.0081 | 0.0000 | -9808.4216 |
| 1.Only_Selection | 4.0000 | 19819.8782 | 306.0150 | 0.0000 | -9905.9250 |

Supplementary table S7: AICc results for models fit for activity assay

| *Effect* | *SS* | *MS* | *dfnum* | *dfden* | *F* | *P* | *adj. P* |
|---|---|---|---|---|---|---|---|
| Selection | 435128.1862 | 435128.1862 | 1.0000 | 5.9986 | 8.6488 | 0.0259 | 0.0518 |
| Sex | 11736375.9584 | 11736375.9584 | 1.0000 | 1414.0071 | 233.2784 | 0.0000 | 0.0000 |
| Assay_set | 439173.0442 | 439173.0442 | 1.0000 | 1414.0142 | 8.7292 | 0.0032 | 0.0078 |
| Selection:Sex | 58613.2337 | 58613.2337 | 1.0000 | 1414.0071 | 1.1650 | 0.2806 | 0.4116 |
| Selection:Assay_set | 5295268.2011 | 5295268.2011 | 1.0000 | 1414.0142 | 105.2515 | 0.0000 | 0.0000 |

Supplementary table S8: ANOVA table for activity assay

| *Parameter* | *Eta2_partial* | *CI* | *CI_low* | *CI_high* |
|---|---|---|---|---|
| Selection | 0.5905 | 0.9500 | 0.0722 | 1.0000 |
| Sex | 0.1416 | 0.9500 | 0.1151 | 1.0000 |
| Assay_set | 0.0061 | 0.9500 | 0.0012 | 1.0000 |
| Selection:Sex | 0.0008 | 0.9500 | 0.0000 | 1.0000 |
| Selection:Assay_set | 0.0693 | 0.9500 | 0.0495 | 1.0000 |

Supplementary table S9: partial eta-squared table for activity assay

| *contrast* | *estimate* | *SE* | *t.ratio* | *p.value* | *adj_pval* |
|---|---|---|---|---|---|
| CON Set1 - FP Set1 | -2.1826 | 43.8450 | -0.0498 | 1.0000 | 1.0000 |
| CON Set1 - CON Set2 | 86.7811 | 16.8010 | 5.1652 | 0.0000 | 0.0000 |
| CON Set1 - FP Set2 | -159.1644 | 43.8222 | -3.6321 | 0.0332 | 0.0577 |
| FP Set1 - CON Set2 | 88.9637 | 43.8314 | 2.0297 | 0.2635 | 0.3952 |
| FP Set1 - FP Set2 | -156.9818 | 16.8014 | -9.3434 | 0.0000 | 0.0000 |
| CON Set2 - FP Set2 | -245.9454 | 43.8086 | -5.6141 | 0.0035 | 0.0082 |

Supplementary table S10: Tukey HSD results for activity assay ANOVA

| | Left | | Right | | |
|---|---|---|---|---|---|
| | Female | Male | Female | Male | Total |
| Control | | | | | |
| C-1 | 18 | 20 | 19 | 20 | 77 |
| C-2 | 19 | 19 | 16 | 20 | 74 |
| C-3 | 17 | 19 | 17 | 18 | 71 |
| C-4 | 18 | 20 | 17 | 19 | 74 |
| Mean | 18.0 | 19.5 | 17.2 | 19.2 | 74.0 |
| Flight Performance | | | | | |
| FP-1 | 19 | 20 | 20 | 19 | 78 |
| FP-2 | 17 | 14 | 17 | 20 | 68 |
| FP-3 | 19 | 19 | 19 | 20 | 77 |
| FP-4 | 20 | 19 | 20 | 20 | 79 |
| Mean | 18.8 | 18.0 | 19.0 | 19.8 | 75.5 |
| Total | 147 | 150 | 145 | 156 | 598 |

Supplementary table S11: Counts of wings analyzed by sex, selection, and replicate.

| *Effect* | *df* | *SS* | *MS* | $R^2$ | *F* | *Z* | *P-value* |
|---|---|---|---|---|---|---|---|
| Female-Control | | | | | | | |
| Individual Variation | 65 | 1.4394892 | 0.0221460 | 0.990 | 97.58 | 16.89 | 0.0003 |
| DA | 1 | 0.0000321 | 0.0000321 | 0.000 | 0.14 | -0.56 | 0.8185 |
| FA | 65 | 0.0147521 | 0.0002270 | 0.010 | | | |
| Female-Flight Performance | | | | | | | |
| Individual Variation | 71 | 1.5409696 | 0.0217038 | 0.984 | 61.03 | 15.24 | 0.0003 |
| DA | 1 | 0.0001525 | 0.0001525 | 0.000 | 0.43 | 0.00 | 0.6181 |
| FA | 71 | 0.0252480 | 0.0003556 | 0.016 | | | |
| Male-Control | | | | | | | |
| Individual Variation | 74 | 1.5877971 | 0.0214567 | 0.992 | 133.81 | 17.81 | 0.0003 |
| DA | 1 | 0.0001417 | 0.0001417 | 0.000 | 0.88 | 0.44 | 0.4837 |
| FA | 74 | 0.0118659 | 0.0001604 | 0.007 | | | |
| Male-Flight Performance | | | | | | | |
| Individual Variation | 70 | 1.3992685 | 0.0199895 | 0.990 | 101.24 | 18.60 | 0.0003 |
| DA | 1 | 0.0000002 | 0.0000002 | 0.000 | 0.00 | -1.97 | 1.0000 |
| FA | 70 | 0.0138212 | 0.0001974 | 0.010 | | | |

Supplementary table S12: Results of analyses of bilateral symmetry in wing centroid size analyzed groupwise. Individual Variation represents the variation explained by individuals. DA is variation attributable to directional asymmetry, and FA to fluctuating asymmetry.

| *Effect* | *df* | *SS* | *MS* | $R^2$ | *F* | *Z* | *P-value* |
|---|---|---|---|---|---|---|---|
| Female-Control | | | | | | | |
| Individual Variation | 65 | 0.0539900 | 0.0008306 | 0.794 | 4.04 | 14.91 | 0.0003 |
| DA | 1 | 0.0005968 | 0.0005968 | 0.009 | 2.90 | 2.91 | 0.0037 |
| FA | 65 | 0.0133700 | 0.0002057 | 0.197 | | | |
| Female-Flight Performance | | | | | | | |
| Individual Variation | 71 | 0.0645405 | 0.0009090 | 0.754 | 3.12 | 13.00 | 0.0003 |
| DA | 1 | 0.0004230 | 0.0004230 | 0.005 | 1.45 | 1.06 | 0.2279 |
| FA | 71 | 0.0206658 | 0.0002911 | 0.241 | | | |
| Male-Control | | | | | | | |
| Individual Variation | 74 | 0.0624708 | 0.0008442 | 0.762 | 3.25 | 15.77 | 0.0003 |
| DA | 1 | 0.0002795 | 0.0002795 | 0.003 | 1.08 | 0.35 | 0.4922 |
| FA | 74 | 0.0191994 | 0.0002595 | 0.234 | | | |
| Male-Flight Performance | | | | | | | |
| Individual Variation | 70 | 0.0620164 | 0.0008859 | 0.782 | 3.73 | 17.23 | 0.0003 |
| DA | 1 | 0.0006759 | 0.0006759 | 0.009 | 2.84 | 2.95 | 0.0041 |
| FA | 70 | 0.0166316 | 0.0002376 | 0.210 | | | |

Supplementary table S13: Results of analyses of bilateral symmetry in wing shape analyzed groupwise. Individual Variation represents the variation explained by individuals. DA is variation attributable to directional asymmetry, and FA to fluctuating asymmetry.

| **Term** | **Estimate** | **SE** | **df** | **t** | **P-value** |
|---|---|---|---|---|---|
| DA: Centroid size | | | | | |
| Selection (FP) | -0.0012 | 0.0049 | 11.3 | -0.25 | 0.9035 |
| Sex (M) | 0.0029 | 0.0036 | 274.1 | 0.81 | 0.5298 |
| Selection x Sex | -0.0010 | 0.0051 | 274.1 | -0.19 | 0.9184 |
| FA: Centroid size | | | | | |
| Selection (FP) | 0.0020 | 0.0041 | 9.0 | 0.48 | 0.7546 |
| Sex (M) | -0.0027 | 0.0026 | 273.7 | -1.03 | 0.4266 |
| Selection x Sex | -0.0002 | 0.0036 | 273.7 | -0.05 | 1.0000 |
| FA: Shape | | | | | |
| Selection (FP) | 0.0024 | 0.0028 | 8.1 | 0.88 | 0.5265 |
| Sex (M) | 0.0012 | 0.0014 | 274.0 | 0.83 | 0.5265 |
| Selection x Sex | -0.0027 | 0.0020 | 274.0 | -1.35 | 0.2739 |

Supplementary table S14: Summary of multilevel ANOVA models with replicate as a random effect comparing directional asymmetry in centroid size and fluctuating asymmetry in centroid size and multivariate shape. It is not possible to test for overall patterns of directional asymmetry in this way.

| *Effect* | *SS* | *MS* | *dfnum* | *dfden* | *F* | *P* | *adj. P* |
|---|---|---|---|---|---|---|---|
| Selection | 1.748e-04 | 1.748e-04 | 1.0000 | 5.8290 | 0.0414 | 0.8457 | 0.9184 |

Supplementary table S15: ANOVA tables for survivorship assay

| *Parameter* | *Eta2_partial* | *CI* | *CI_low* | *CI_high* |
|---|---|---|---|---|
| Selection | 0.0070 | 0.9500 | 0.0000 | 1.0000 |

Supplementary table S16: partial eta-squared tables for survivorship assay

| *Model* | *Effect* | *SS* | *MS* | *dfnum* | *dfden* | *F* | *P* | *adj. P* |
|---|---|---|---|---|---|---|---|---|
| Females | Selection | 0.0009 | 0.0009 | 1.0000 | 6.0000 | 0.0217 | 0.8878 | 0.9451 |
| Males | Selection | 0.1672 | 0.1672 | 1.0000 | 6.0000 | 8.4801 | 0.0269 | 0.0522 |

Supplementary table S17: ANOVA tables for dry mass assay

| *Model* | *Parameter* | *Eta2_partial* | *CI* | *CI_low* | *CI_high* |
|---|---|---|---|---|---|
| Selection × Sex | Selection | 0.2357 | 0.9500 | 0.0000 | 1.0000 |
| Selection × Sex | Sex | 0.9099 | 0.9500 | 0.8967 | 1.0000 |
| Selection × Sex | Selection:Sex | 0.0414 | 0.9500 | 0.0127 | 1.0000 |

Supplementary table S18: partial eta-squared table for dry mass assay

| *Model formula* | *K* | *AICc* | *Delta_AICc* | *AICc weight* | *LL* |
|---|---|---|---|---|---|
| Selection | 4.0000 | 102.7577 | 0.0000 | 0.9932 | -46.3262 |
| Selection * Plate | 8.0000 | 112.7367 | 9.9789 | 0.0068 | -43.5683 |

Supplementary table S19: AICc results for models fit for fecundity assay (log fecundity)

| *Model* | *Effect* | *SS* | *MS* | *dfnum* | *dfden* | *F* | *P* | *adj. P* |
|---|---|---|---|---|---|---|---|---|
| percapita_Fecundity | Selection | 0.0152 | 0.0152 | 1.0000 | 6.0000 | 4.0144 | 0.0920 | 0.1480 |
| log_percapita_Fecundity | Selection | 10.3139 | 10.3139 | 1.0000 | 6.0000 | 4.8112 | 0.0707 | 0.1197 |

Supplementary table S20: ANOVA tables for fecundity assay (1. per-capita fecundity; 2. log-transformed per-capita fecundity)

| *Model* | *Parameter* | *Eta2_partial* | *CI* | *CI_low* | *CI_high* |
|---|---|---|---|---|---|
| percapita_Fecundity | Selection | 0.4009 | 0.9500 | 0.0000 | 1.0000 |
| log_percapita_Fecundity | Selection | 0.4450 | 0.9500 | 0.0000 | 1.0000 |

Supplementary table S21: partial eta-squared table for fecundity assay (1. per-capita fecundity; 2. log-transformed per-capita fecundity)

| *Selection* | *estimate* | *statistic* | *p.value* | *parameter* | *conf.low* | *conf.high* | *method* | *alternative* |
|---|---|---|---|---|---|---|---|---|
| CON | -0.3972 | -1.3687 | 0.2010 | 10.0000 | -0.7908 | 0.2288 | Pearson's product-moment correlation | two.sided |
| FP | -0.0404 | -0.1277 | 0.9009 | 10.0000 | -0.6004 | 0.5462 | Pearson's product-moment correlation | two.sided |

Supplementary table S22: correlation tables for fecundity assay for number of females vs. fecundity

**Variant calling pipeline**

| **Filter** | **Annotation** | **Description** |
|---|---|---|
| QD < 2.0 | QualByDepth | Variant confidence divided by read depth. Low values suggest the call is poorly supported relative to coverage, often indicating a false positive. |
| QUAL < 30.0 | QualityScore | Phred-scaled probability that the variant is real. Below 30 means less than 99.9% confidence in the call. |
| SOR > 3.0 | StrandOddsRatio | Measures strand bias (whether the variant is supported more by reads on one strand than the other). High values suggest sequencing artifact. |
| FS > 60.0 | FisherStrand | Strand bias measured by Fisher’s exact test. High values indicate reads supporting the variant are disproportionately on one strand. |
| MQ < 40.0 | MappingQuality | Average mapping quality of reads supporting the variant. Low values mean reads are poorly mapped and the call may be unreliable. |
| MQRankSum < −12.5 | MappingQualityRankSumTest | Compares mapping quality of reads supporting the reference vs. the alternate allele. Strongly negative values mean alt-supporting reads are mapped much worse, suggesting a false positive. |
| ReadPosRankSum < −8.0 | ReadPositionRankSumTest | Compares the position of the variant within reads supporting ref vs. alt. Strongly negative values mean the variant tends to appear near read ends, a common artifact. |

Supplementary table S23: SNP hard filter thresholds and descriptions.

| Filter | Annotation | Description |
|---|---|---|
| QD < 2.0 | QualByDepth | Same as SNPs; low confidence relative to depth. |
| QUAL < 30.0 | QualityScore | Same as SNPs; low overall call confidence. |
| FS > 200.0 | FisherStrand | Same strand bias metric as SNPs, but with a more lenient threshold as indels naturally show higher strand bias. |
| ReadPosRankSum < −20.0 | ReadPositionRankSumTest | Same read position test as SNPs, with a more lenient threshold as indels near read ends are more common even in true calls. |

Supplementary table S24: Indel hard filter thresholds and descriptions.

**Gene Ontology and Enrichment analyses results are in Zenodo (https://doi.org/10.5281/zenodo.23356991)**

Supplementary table S25: enrichment analysis results for significantly differentiated SNPs based on linear model.

Supplementary table S26: enrichment analysis results for significantly differentiated SNPs based on haplotype analysis.

Supplementary table S27: genes overlapping with Spierer et al. 2021.

Supplementary table S28: genes overlapping with Pitcher et al. 2019.
